# TC10 regulates breast cancer invasion and metastasis by controlling membrane type-1 matrix metalloproteinase at invadopodia

**DOI:** 10.1101/2020.11.17.386854

**Authors:** M. Hülsemann, S.K. Donnelly, P.V. Verkhusha, S.P.H. Mao, J.E. Segall, L. Hodgson

## Abstract

During breast cancer metastasis, cancer cell invasion is driven by actin-rich protrusions called invadopodia, which mediate the extracellular matrix degradation required for the success of the invasive cascade. In this study, we demonstrated that TC10, a member of a Cdc42 subfamily of p21 small GTPases, regulates the membrane type 1 matrix metalloproteinase (MT1-MMP)-driven extracellular matrix degradation at invadopodia. We show that TC10 is required for the plasma membrane surface exposure of MT1-MMP at invadopodia. By utilizing our new Förster resonance energy transfer (FRET) biosensor, we demonstrated the p190RhoGAP-dependent regulation of spatiotemporal TC10 activity at invadopodia. We identified a pathway that regulates TC10 activity and function at invadopodia through the activation of p190RhoGAP and the downstream interacting effector Exo70 at the invadopodia sites. Our findings reveal the role of a previously unknown regulator of vesicular fusion at invadopodia, TC10, on the invasive potential of breast cancer cells during invasion and metastasis.

## Introduction

Cancer metastasis represents a multistep process, during which cells escape from a primary tumor and disseminate throughout the body, establishing new tumors at distant sites. To achieve this dissemination, cancer cells form actin-rich protrusions called invadopodia. Mature invadopodia degrade the extracellular matrix (ECM) by recruiting membrane type 1 matrix metalloproteinases (MT1-MMP), a transmembrane protease that has been associated with ECM degradation during mammary adenocarcinoma invasion^1^. Invadopodia structures are spatially and temporally regulated^2^ and are necessary to breach the basement membrane and degrade the ECM during the intra/extravasation process^3–8^. Although the therapeutic efficacy of proteinase inhibitors has not been successfully established in clinical applications^9,10^, *in vitro* and *in vivo* studies have indicated that MT1-MMP-mediated functions play important roles during the breast tumor metastatic cascade. These issues highlight our lack of clear understanding regarding the mechanisms that underlie the tumor invasion and dissemination processes, preventing the clear delineation of the contributions made by proteinase-dependent^11–13^ and -independent^14^ processes during tumor invasion and metastasis.

TC10 is a p21 small GTPase that belongs to the Rho family and is closely related to Cdc42, a canonical small GTPase. The role played by Cdc42 in the regulation of invadopodia generation has previously been demonstrated^12,15^. A minor paralog of Cdc42, TC10, has not yet been established as a key player in tumor invasion and metastasis^16–19^, although the involvement of TC10 has been recognized in other disease contexts, including diabetes^20^. In general, Rho-family GTPases serve as molecular switches that cycle between the GTP-bound on state and the GDP-bound off state. GTPases are regulated by guanine nucleotide exchange factors (GEFs), which exchange GDP for GTP, GTPase-activating proteins (GAPs), which induce GTP hydrolysis, and guanine nucleotide dissociation inhibitors (GDIs), which can prevent the GDP to GTP exchange. Unlike other canonical Rho GTPases, TC10 has a relatively low binding affinity for Mg^2+^, suggesting that wild-type (WT) TC10 may act as a fast-cycling GTPase, remaining in an activated state unless acted upon by GTPase regulators, such as GAPs^21,22^. TC10 is highly active on exocytic vesicles and recycling endosomes, and the TC10-mediated hydrolysis of GTP is necessary to promote vesicular fusion at the plasma membrane^23^. TC10 interacts with Exo70 as part of a conserved, octameric exocyst complex that recruits TC10-loaded vesicles to the plasma membrane^24,25^. This function of TC10 is conserved in neurites^26^, suggesting that TC10 activity is broadly important for exocytosis. The docking of the exocyst complex at invadopodia has been observed in breast cancer cells, where it appears to control the exocytic presentation of MT1-MMP^27^. These observations indicate the likely involvement of a yet unknown vesicular fusion regulator that may be necessary to complete the final step of MT1-MMP surface presentation at tumor invadopodia^28^.

In this study, we showed that endogenous TC10 is localized at invadopodia and that TC10 depletion markedly reduced ECM degradation and the *in vitro* invasion of mammary adenocarcinoma cells through Matrigel-coated filters. We identified an important control node for the TC10 GTPase function involving p190RhoGAP, which is necessary for the regulation of TC10 activity at invadopodia. We observed the activation dynamics of TC10 at invadopodia using our new, sensitive, Förster resonance energy transfer (FRET)-based TC10 biosensor. Importantly, we demonstrated that the TC10-mediated hydrolysis of GTP, which is promoted by p190RhoGAP, was required for matrix degradation and the surface exposure of MT1-MMP at invadopodia. Moreover, we showed that TC10 significantly impacts breast tumor metastasis to the lungs in an *in vivo* mouse orthotopic model of breast cancer metastasis. Taken together, our results indicated an important role for TC10 as a regulator of exocytic vesicular control at invadopodia, involved in matrix degradation, invasion, and metastasis in breast cancer.

## Results

### TC10 is localized at invadopodia and is necessary for matrix degradation

TC10 is known to function in vesicular trafficking, especially during glucose receptor transport in diabetes^29^; however, its role in cancer has not yet been elucidated. We hypothesized that TC10 might impact cancer invasion and metastasis by regulating the functions of tumor invadopodia. We found that endogenous TC10 was localized at invadopodia in two different breast cancer cell lines: rat adenocarcinoma MTLn3 (Fig. 1a) and human triple-negative MDA–MB-231 (Supplementary Fig. S1). Endogenous TC10 at invadopodia displayed two distinct localization patterns, either laterally at the sides of invadopodia or within the core of invadopodia, overlapping with the cortactin/Tks5 core marker proteins (Fig. 1b and c; Supplementary Fig. S1). Under steady-state conditions, TC10 localized predominantly to the invadopodia core (Fig. 1b). To characterize TC10 localization during the early phases of invadopodia formation, we serum-starved MTLn3 cells and then stimulated them with epidermal growth factor (EGF) to induce the synchronous formation of invadopodium precursors, which are unable yet to degrade the ECM. We found that TC10 was initially partitioned equally between the core and the regions surrounding the core, whereas 5 min after EGF stimulation, TC10 was observed to accumulate at the core (Fig. 1d). This timing relative to EGF stimulation coincided with previous observations regarding β1 integrin activation dynamics during EGF-stimulated invadopodia precursor formation^30^ and with the activation of various pathways associated with this important adhesion molecule^31^. These observations may indicate the possible involvement of a β1 integrin-adhesion-mediated pathway in the modulation of TC10 activity and functions at invadopodia.

**Figure 1:**
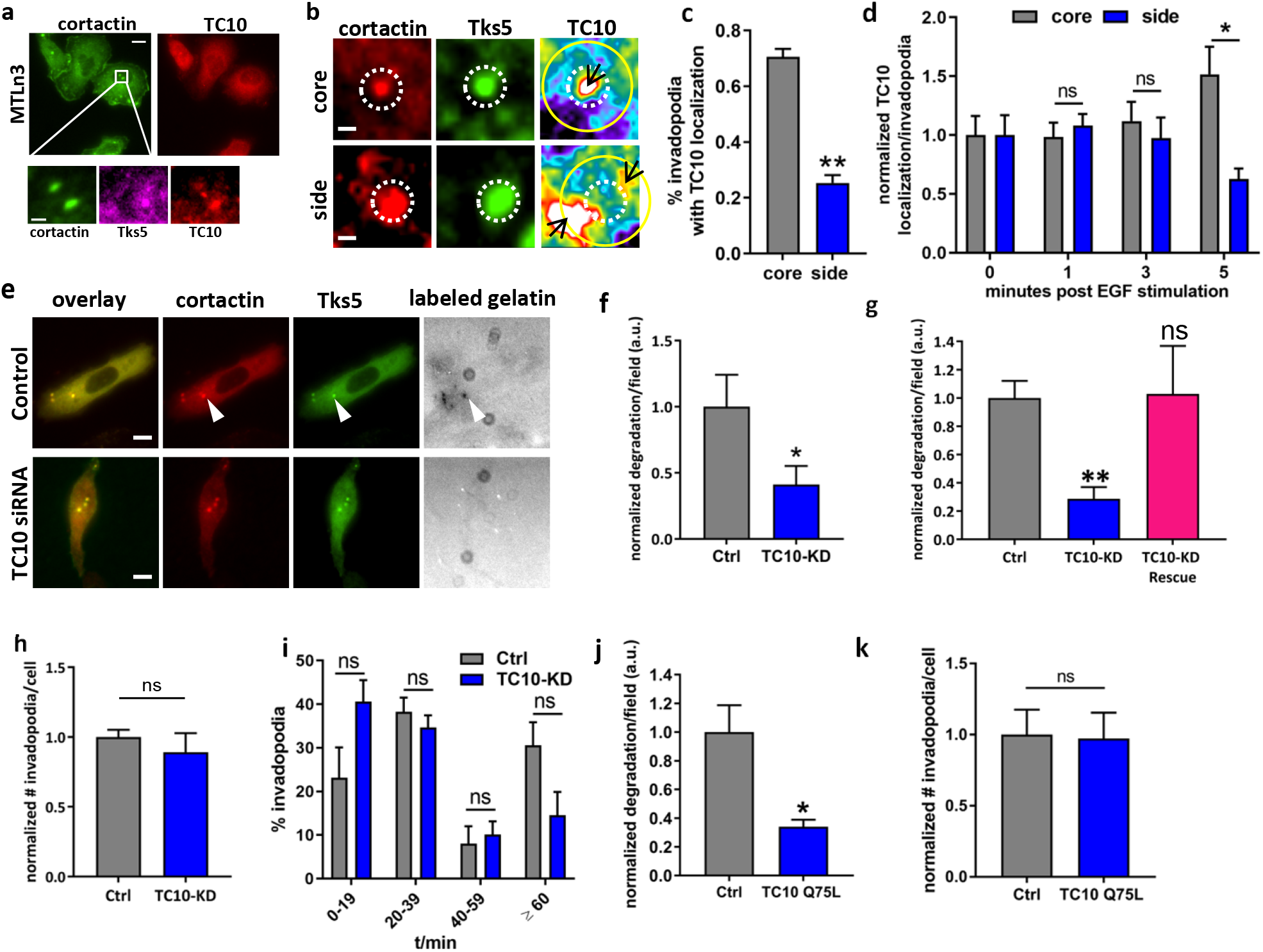
TC10 is localized at invadopodia and is required for matrix degradation. **a.** Representative localization of endogenous TC10 in rat mammary adenocarcinoma MTLn3 cells. Cortactin is shown to denote invadopodia structures. White box area is enlarged to show colocalizations (bottom), showing cortactin, Tks5 and TC10 localizations. White bar = 10-μm (top); 2-μm (bottom). **b.** Representative, enlarged view of the immunostaining of cortactin and Tks5 with TC10-WT-mCherry expression. White bar = 1-μm. The “side” localizations (yellow circle) were identified within a dilated circular region of approximately 30 pixels from the invadopodia core (dashed white circle). Black arrows point to TC10 localizations. **c.** Quantification of TC10-WT-mScarlet localization at invadopodia structures. Student’s t-test, two-tail analysis: ** p=0.001112; n=5 experiments; shown with SEM **d.** Quantification of TC10-WT-mCherry localization in MTLn3 cells stimulated with 5 nM EGF for the indicated times. Results were normalized to t = 0-min values for both side and core localized fractions. Student’s t-test, two-tail analysis: ns p=0.5602 for 1-min side versus core; p=0.5688 for 3-min side versus core; * p=0.01090 for 5-min side versus core; n=4 experiments; shown with SEM. **e.** Representative images from siRNA-mediated TC10 depletion in MDA–MB-231 cells impacting gelatin matrix degradation, visualized using a 405-nm fluorescent gelatin matrix. Invadopodia are denoted by cortactin and Tks5 colocalization with spots of matrix degradation (arrow). White bar = 10-μm. **f.** Quantification of the MDA-MB-231 matrix degradation when TC10 is depleted, shown in (**e**), normalized to the Ctrl. Student’s t-test, one-tail analysis: * p=0.01528; n=3 experiments; shown with SEM. **g.** siRNA-mediated TC10 depletion in MTLn3 cells and rescue mediated by the overexpression of wild-type TC10. Results are normalized against Ctrl. Student’s t-test, one-tail analysis: ** p=0.004146; two-tail analysis: ns p=0.9669; n=3 experiments; shown with SEM. **h.** Total number of steady-state invadopodia/cell in TC10-depleted MTLn3 cells normalized to Ctrl MTLn3 cells. Student’s t-test, two-tail analysis: ** p=0.7994; n=3 experiments; shown with SEM. **i.** Invadopodia lifetimes in MTLn3 cells, shown as a histogram with bins corresponding to 20-min intervals. Student’s t-test, two-tail analysis: ns, p=0.1106 for 0-19 min; p=0.4472 for 20-40 min; p=0.6922 for 41-60 min; p=0.09957 for >60 min; n=3 experiments, shown with SEM. **j.** Overexpression of the GTP hydrolysis-deficient TC10 Q75L mutant in MTLn3 cells plated on a 405-nm fluorescent gelatin matrix, normalized to Ctrl overexpressing wild-type TC10. Student’s t-test, one-tail analysis: * P=0.01349; n=3 experiments; shown with SEM. paired one-tail **k.** Total number of steady-state invadopodia in cells overexpressing wild-type TC10 or the Q75L mutant, normalized to the Ctrl overexpressing the wild-type TC10. Student’s t-test, two-tail analysis: ns P=0.4824; n=3 experiments; shown with SEM. siRNA depletion characterizations are shown in Supplementary Figures S2 and S3.

We next used small interfering RNA (siRNA) to deplete TC10, which resulted in reduced ECM degradation in MDA–MB-231 cells (Fig. 1e and f). A similar phenotype was observed in MTLn3 cells, which could be rescued by the overexpression of a WT TC10 construct that is resistant to siRNA (Fig. 1g). The steady-state number of invadopodia was not affected by TC10 depletion (Fig. 1h), indicating a limited role for TC10 in the structural aspects of invadopodia regulation. Corroborating this observation, the lifetimes (turnover rates) of invadopodia were not significantly impacted by TC10 depletion (Fig. 1i). To explore the functional role of TC10 on its effects on matrix degradation, we overexpressed a TC10-Q75L mutant^23^, which lacks the catalytic ability to hydrolyze GTP. The overexpression of TC10-Q75L resulted in an ECM degradation defect similar to that observed under TC10 depletion conditions (Fig. 1k), and no effect on the total number of invadopodia was observed (Fig. 1l). Together, these observations indicated that the ability of TC10 to hydrolyze GTP and its GTPase cycling activity are necessary to regulate the ECM degradation function of invadopodia. These results also suggested that TC10 plays a functional role, rather than a structural role, during invadopodia dynamics and tumor invasion.

### TC10 regulates MT1-MMP exposure at the plasma membrane of invadopodia

Because TC10 plays a well-known role in vesicular trafficking^23^, in addition to the impacts on ECM degradation at invadopodia observed in the previous experiment, we hypothesized that TC10 regulates the MT1-MMP surface presentation at invadopodia by controlling vesicular fusion at the plasma membrane during exocytosis. To test this hypothesis, we first examined the endogenous localization of MT1-MMP at invadopodia and found two distinct patterns of localization: within the invadopodia core and laterally flanking the invadopodia core (Fig. 2a), with the side localization being more predominant (Fig. 2b). We then overexpressed an MT1-MMP with an enhanced green fluorescent protein (EGFP) tag on the C-terminal, cytoplasmic end. We overexpressed this EGFP-tagged MT1-MMP construct in MTLn3 cells and antibody stained the surface-exposed MT1-MMP without permeabilizing the plasma membrane. When TC10 was depleted in these cells, we observed a significant reduction in the proportion of surface-exposed MT1-MMP staining relative to the total MT1-MMP level, as measured by tracking the EGFP fluorescence intensity (Fig. 2c). No difference in total MT1-MMP levels at invadopodia was observed between the control and TC10-depleted cells (Fig. 2c). These observations suggested that TC10 plays an important role in the surface exposure of MT1-MMP at the plasma membrane, without affecting the trafficking of MT1-MMP-containing vesicles or the loading of MT1-MMP cargo onto vesicles.

**Figure 2:**
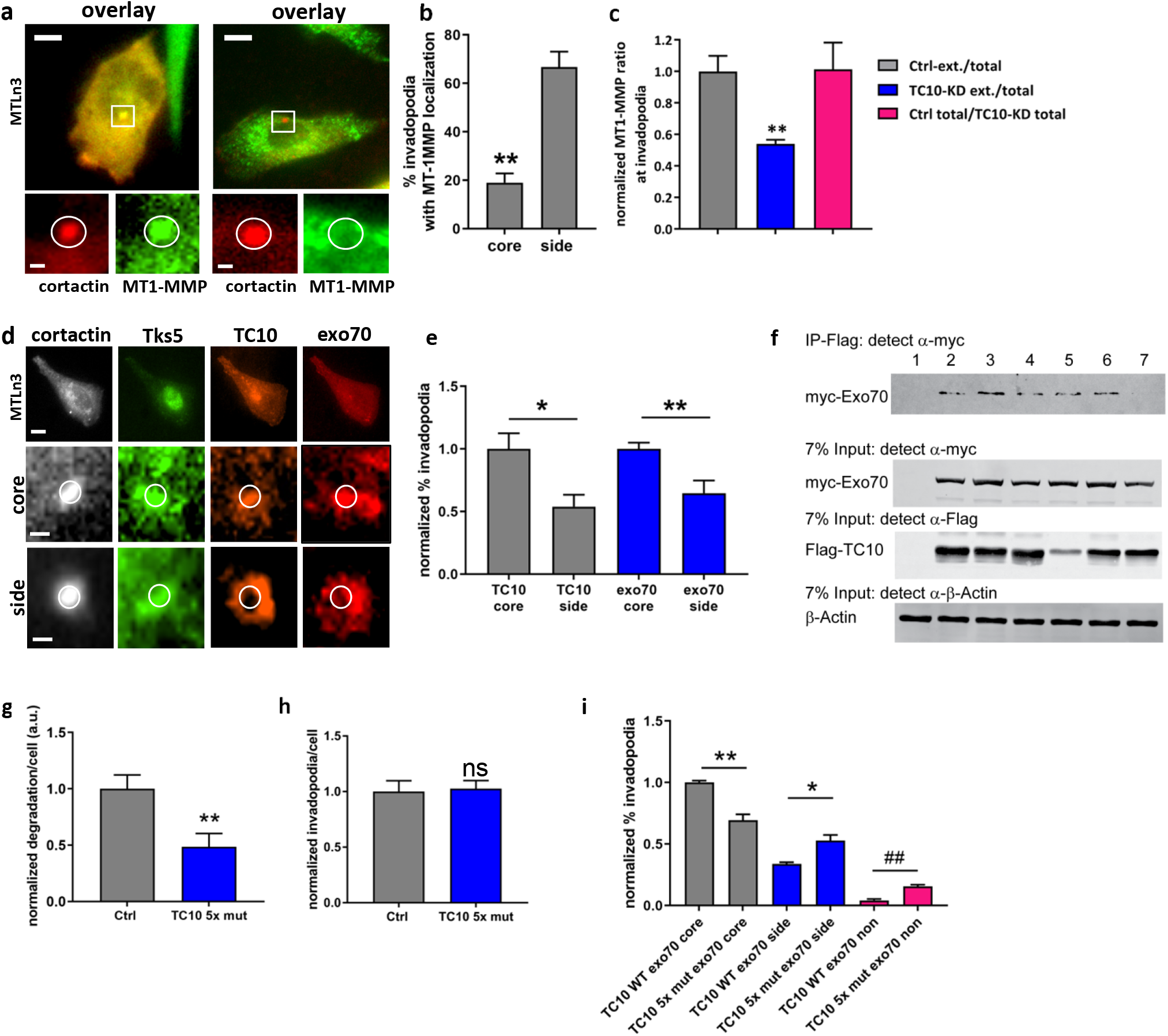
TC10 regulates MT1-MMP cell surface exposure at the plasma membrane of invadopodia. **a.** Representative, localization of endogenous MT1-MMP at invadopodia in MTLn3 cells. Invadopodia are denoted by cortactin staining. White bar in top left = 5-μm; top right = 10-μm; zoomed views = 1-μm. **b.** Quantification of MT1-MMP-WT-GFP localization in MDA-MB-231, either within the invadopodia core or on the lateral sides of the core. Invadopodia cores are denoted by the colocalization of cortactin and Tks5 signals. Student’s t-test, two-tail analysis: ** p=0.003055; n=3 experiments; shown with SEM. Data from MTLn3 cells shown similar trend is shown in Supplementary Figure S4a. **c.** The extracellular surface presentation of MT1-MMP at invadopodia requires TC10 in MTLn3 cells. The ratio between numbers of invadopodia with cytoplasmic (total) MT1-MMP and those featuring surface (ext.) MT1-MMP in cells treated with Ctrl (gray) or TC10 siRNA (blue). The ratio of total MT1-MMP-positive invadopodia counts in cells treated with Ctrl and TC10 siRNA (magenta), indicating that only the surface presentation of MT1-MMP is impacted by TC10 depletion. TC10-KD data are normalized to the Ctrl-ext. over total ratio. Student’s t-test, one-tail analysis: ** p=0.005750; n=3 experiments; shown with SEM. **d.** Representative, immunostaining of TC10 and Exo70 at the invadopodia site in MTLn3 cells, showing the side and the core localizations. White bar in top = 10-μm; zoomed views = 1-μm. **e.** The quantification of TC10 and exo70 localization at invadopodia in MTLn3 cells, as shown in (**d**), normalized to the core % for TC10 and Exo70, respectively. Student’s t-test, two-tail analysis: ** p=0.007508; * p=0.01124; n=8 experiments; shown with SEM. **f.** Immunoprecipitation of wild-type (WT) Exo70 and TC10 mutants, overexpressed in HEK293T cells. Lanes: 1, untransfected; 2, WT TC10; 3, F42L TC10; 4, Q75L TC10; 5, T31N TC10; 6, P43L/E45V/Y46H (3× mut) TC10; and 7, P43L/E45V/Y46H/T49A/Y54C (5× mut) TC10. Full-sized western blots are shown in Supplementary Figure S13. **g.** Matrix degradation per cell, comparing WT TC10 and 5× mut TC10, overexpressed in MTLn3 cells, as plated on a 405-nm fluorescent gelatin matrix. Results are normalized to the Ctrl. Student’s t-test, paired one-tail analysis: ** p=0.005604; n=5 experiments; shown with SEM. **h.** Total number of steady-state invadopodia, comparing WT TC10 and 5× mut TC10, overexpressed in MTLn3 cells. Results are normalized to the Ctrl. Student’s t-test, paired two-tail analysis: ns p=0.7283; n=5 experiments; shown with SEM. **i.** Localization of exo70 at invadopodia (core/side/no localization) in MTLn3 cells, comparing the overexpression of WT TC10 and 5× mut TC10. Student’s t-test, paired two-tail analysis: ** p=0.002105; * p=0.01138; ## p=0.00003960 n=5 experiments; shown with SEM.

Previously, the exocyst complex was observed to dock onto the lateral aspect of invadopodia, which was shown to be important for ECM degradation by invadopodia in breast cancer invasion ^32^. Exo70 is a component of the octameric exocyst complex, which plays a critical role in vesicular docking at the cell membrane and is essential for the exocytic secretion of MMP at invadopodia^33^. Because TC10 has been shown to interact with Exo70^34^, we examined the role played by the TC10–Exo70 interaction on MT1-MMP surface exposure and ECM degradation. We found that Exo70 localization strongly overlapped with TC10, both within the invadopodia core and at the lateral aspects of invadopodia (Fig. 2d). Unlike the MT1-MMP localization, however, Exo70 was predominantly localized with TC10 in the invadopodia core (Fig. 2e). P29L/E31V/Y32H mutations in the switch I/II regions of Cdc42, which are important for effector interactions, have been shown previously to disrupt Cdc42–Exo70 co-immunoprecipitation^35,36^. When analogous mutations (P43L/E45V/Y46H) were introduced into TC10, we observed only a modest reduction in the co-immunoprecipitation of Exo70 with TC10 (Lane 6: Fig. 2f). When two additional effector binding mutations were introduced (T49A and Y54C, which are analogous to mutations in Cdc42 that impact activity status and effector interactions)^37^, a further reduction in co-immunoprecipitation was observed (Lane 7: Fig. 2f), suggesting that these additional mutations further interfered with complex formation. Although the constitutively active, GTP-hydrolysis-deficient version of TC10 (Q75L) only modestly immunoprecipitated with Exo70, similar to the dominant-negative T31N TC10 mutant (Lanes 4 and 5: Fig. 2f), the F42L mutation, which is analogous to a mutation that renders other RhoGTPases into fast-cycling GTPases, resulted in the strong immunoprecipitation of Exo70 (Lane 3: Fig. 2f). These results suggested that GTP hydrolysis and nucleotide cycling activity is important for efficient TC10–Exo70 complex interactions and that mutating the residues P43L/E45V/Y46H/T49A/Y54C (“5×-mutation”) within the Switch I/II region of TC10 was able to impact this interaction. We then used this 5×-mutated TC10, co-expressed with Exo70 in MTLn3 cells, and observed a significant impact on the ability of these cells to degrade the ECM (Fig. 2g), without affecting the total number of steady-state invadopodia (Fig. 2h). The expression of the 5×-mutated version of TC10 altered the localization pattern of Exo70 at invadopodia (Fig. 2i) but did not alter the TC10 localization pattern in invadopodia (Supplementary Fig. S4b). These observations indicated that the TC10-Exo70 interaction is important for the appropriate targeting of Exo70 within invadopodia and that this interaction impacts ECM degradation. Furthermore, these results indicated a potential mechanism through which MT1-MMP might be deposited into the plasma membrane, as MT1-MMP-loaded vesicles containing TC10 approach the lateral/side aspect of invadopodia. GTP hydrolysis by vesicular-bound TC10 may begin to occur in this region, where TC10 might first encounter a cognate GAP that resides within the invadopodium core, which facilitates GTP hydrolysis by TC10 to promote the plasma membrane fusion of the vesicles.

### TC10 activity at invadopodia is spatially regulated

Because GTP hydrolysis is necessary for TC10-mediated vesicular fusion at the plasma membrane^23^, we evaluated the activation dynamics of TC10 at and surrounding the invadopodia. For this purpose, we designed a new FRET-based TC10 biosensor (Fig. 3a). The biosensor design is based on a monomeric, single-chain, genetically encoded approach that is TC10-specific, similar to the design of our previous Rac and Cdc42 sensors^38,39^. The biosensor consists of a monomeric Cerulean 1 and monomeric circularly permuted (cp229) Venus fluorescent protein FRET pair with an optimized Rac/Cdc42-binder motif from our Rac/Cdc42 biosensors^38,40–42^, and full-length TC10 (Fig. 3a). The spectrofluorometric characterization of the TC10 FRET biosensor revealed an approximately 80% difference in FRET/donor emission ratio between the constitutively activated (Q75L) and off states (the dominant-negative T31N or other effector binding mutants) of the TC10 biosensor (Fig. 3b). The WT TC10 version of the biosensor showed high FRET, similar to two different constitutively active TC10 biosensor mutants (G26V and Q75L, Fig. 3c), corroborating previous reports that WT TC10 represents an activated GTPase due to a low Mg^2+^-binding affinity^23^. The co-expression of Rho-targeting p50RhoGAP and p190RhoGAP but not Rap-targeting Rap1GAP1 resulted in the attenuation of FRET (Fig. 3d). The co-expression of caveolin1, a putative GDI for TC10^29^, resulted in attenuation of FRET (Supplementary Fig. S5a). To confirm that the expression of the TC10 biosensor did not result in aberrant overexpression artifacts in downstream signaling, we performed a competitive pull-down assay using purified, exogenous binding domain. The activated TC10 biosensor only interacted with an exogenous effector when both biosensor binding domains within the biosensor were mutated (2XPBD: H83/86D), preventing an interaction between activated TC10 and the GTPase binder motif within the biosensor backbone (Supplementary Fig. S5b). When the TC10 biosensor was overexpressed in MTLn3 breast cancer cells, we observed an approximately 30% difference in whole-cell average TC10 activities between the constitutively active and dominant-negative versions of the TC10 biosensor (Supplementary Fig. S5c). The biosensor also responded to stimulation with serum and EGF following serum starvation (Supplementary Fig. S5d). We then applied a synonymous codon modification^43^, which prevents homologous recombination during transfection and transduction into tumor cells. The TC10 biosensor was stably transduced and integrated into tet-OFF tTA-MTLn3 cells^44^, under the control of a tet-inducible promoter, to achieve tight expression control.

**Figure 3:**
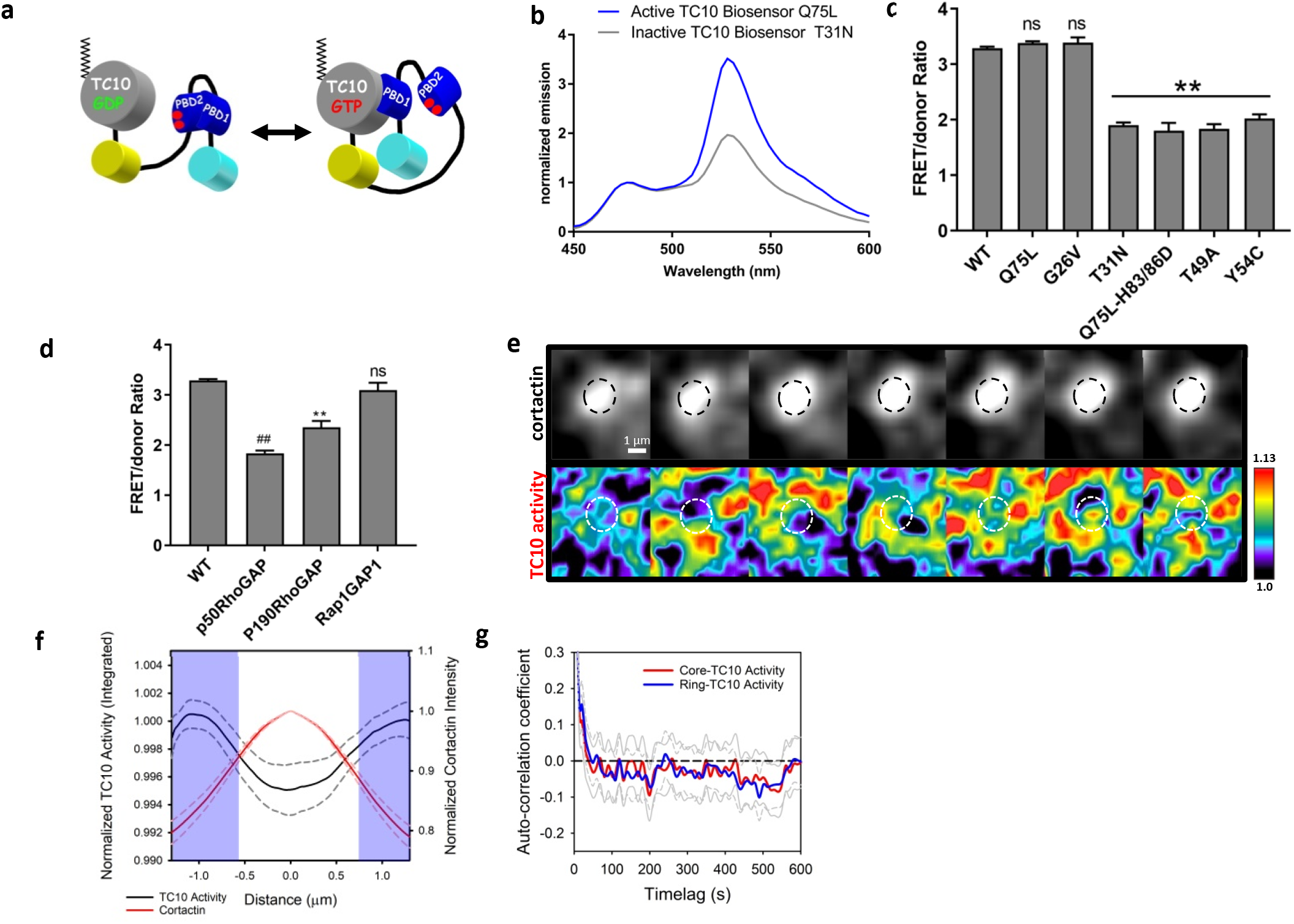
TC10 activity at invadopodia is spatially regulated. **a.** A schematic cartoon of the single-chain, genetically encoded FRET biosensor for TC10 GTPase, based on previous biosensor designs used to evaluate Rac/Cdc42-type GTPases ^38,40–42^. The FRET donor (cyan) and acceptor (yellow) were mCerulean1 and circularly permutated mVenus, respectively. We also produced a near-infrared version of the TC10 FRET biosensor, which behaved similarly to the cyan-yellow version based on a previous design (Supplementary Fig. S6)^85^. **b.** Representative, normalized fluorescence emission spectra of the constitutively activated (CA: Q75L) versus the dominant-negative (DN: T31N) versions of the TC10 biosensor upon excitation at 433 nm when overexpressed in HEK293T cells and measured in cell suspensions. Spectra were normalized to the peaks of the donor emission at 474 nm. **c.** Fluorometric emission ratio of the TC10 biosensor overexpressed in HEK293T cells. WT biosensor expression and the Q75L and G26V CA mutant biosensors showed high emission ratios. The DN biosensor, CA biosensors with GTPase binding-deficient mutations in both PBD domains (Q75L-H83/86D), and effector binding mutants (T49A, Y54C) showed low emission ratios. Student’s t-test, two-tail analysis: ns p=0.06587 for Q75L, p=0.3810 for G26V; ** p=1.544×10^−6^ for T31N, p=0.001531 for Q75L-H83/86D, p=0.0001294 for T49A, and p=0.0001279 for Y54C, all compared to the WT (first bar); n=7 experiments for WT, 4 experiments for all other conditions, all shown with SEM. **d.** The co-expression between p50RhoGAP, p190RhoGAP, and the non-targeting Rap1GAP1 and the WT TC10 biosensor. Student’s t-test, two-tail analysis: ## p=9.899×10^−5^, ** p=0.003775, ns p=0.3204, all compared to the WT TC10 biosensor expression (first bar) without GAP co-expression. N=7 experiments for WT, n=3 experiments for p50RhoGAP and Rap1GAP and n=4 experiments for p190RhoGAP co-expressions, shown with SEM. **e.** Representative, dynamic localization patterns of TC10 activity at and surrounding the invadopodium core (denoted by cortactin fluorescence). Time-lapse sequence intervals are 10 seconds. **f.** The line scan analysis of the intensity distributions across invadopodia showing normalized TC10 activity integrated over time, plotted against the matching cortactin intensity distributions. The blue-shaded regions indicate significant (p < 0.05; Student’s t-test, one-tail; n = 33 invadopodia from 19 cells over 7 experiments; for p-value distributions, see Appendix 1) differences in TC10 activity intensity compared with the invadopodia core center at 0.0 μm. Line scans were normalized to the local maxima of TC10 activity at the ring-like region surrounding the invadopodia core, which was denoted by the cortactin spot. The cortactin intensity was normalized at the center position, taken as the maximal intensity location along the line scans. **g.** Autocorrelation functions showing fluctuations in TC10 activity in the invadopodia core (red) versus the ring-like region (blue) around the invadopodia core. The gray lines (solid: core; dashed: ring) indicate the 95% confidence intervals around the mean. N = 29 invadopodia core and ring measurements, from 19 different cells, in 8 experiments.

Using the new TC10 FRET biosensor, we attempted to determine the dynamics of TC10 activity at and surrounding the invadopodia. We co-transduced cortactin-miRFP703 to serve as a marker of the invadopodia core and observed TC10 activity. The biosensor activities at invadopodia appeared to be highly dynamic and fluctuated markedly during live-cell imaging (Fig. 3e), indicating stochastic behavior over time. We first reduced the complexity of the data temporally by integrating the TC10 activities over time in steady-state invadopodia, similar to a previous analysis performed for a different class of Rho GTPase^39^. We measured and averaged the line scans across invadopodia and determined that the center of the invadopodium core showed a significant reduction in time-integrated TC10 activity compared with the regions surrounding the invadopodium core (Fig. 3f), which suggested the apparent stochastic fluctuation of TC10 activity at invadopodia, underlying spatially ordered distribution of TC10 activity. A similar observation was made in MDA-MB 231 cells (Supplementary Fig. S7). To further quantify the TC10 activity fluctuation, we defined two regions by generating a binary mask, with one region based on the cortactin core (“core”) and the second region defined by dilating the core mask by 30 pixels and subtracting the core to form an annulus (“ring”). The live-cell biosensor measurements were analyzed by autocorrelation to extract the characteristic periodicity^38,45,46^ at these two regions. We observed no periodic fluctuations in TC10 activity for either region, as characterized by the lack of repeated, oscillatory crossings of the zero axis in the autocorrelation functions (Fig. 3g), which indicated that the TC10 activity dynamics in WT, steady-state invadopodia are stochastic in nature. Importantly, the time-integrated activity of TC10 is attenuated significantly in the core, indicating a potential mechanism through which a core-localized GAP may regulate TC10 GTP-hydrolysis activity at the invadopodia core (Fig. 3f).

### p190RhoGAP impacts invadopodia function by targeting TC10

To identify the regulator of TC10 function at invadopodia, we focused on p190RhoGAP (*Arhgap35*), a well-known, integrin-adhesion-associated regulator of RhoGTPases, which binds cortactin within the invadopodia core during invadopodia precursor formation and is present within the core of invadopodia at steady state^7^. Traditionally, p190RhoGAP targets Rac and Rho GTPase isoforms and contains: an N-terminal GTPase-binding domain; four FF domains involved in binding transcription factors; a protrusion localization domain that binds cellular cortactin or Rnd3 GTPase; a p120RasGAP binding site; a polybasic region; and a C-terminal consensus GAP domain that can switch specificity between Rac-GTP and Rho-GTP^47,48^. p190RhoGAP has been associated with the regulation of TC10 activity in a number of systems, including the leading edge of HeLa cells and neurite extensions^23,49^. In melanoma, tyrosine phosphorylation and the activation of p190RhoGAP at invadopodia in response to laminin peptide depends on the activation of β1 integrins^50^; in breast cancer, invadopodia precursor β1 integrins are activated within 3 to 5 min after EGF stimulation in a Rac3 GTPase-dependent manner^30,38^. β1 integrin recruits the non-receptor tyrosine kinase Arg (Ableson-related gene, also known as Abl2) and stimulates the Arg-dependent phosphorylation of p190RhoGAP at the leading edge of fibroblasts; however, whether this occurs in breast cancer cell invadopodia has not yet been elucidated.

Steady-state and EGF-stimulated invadopodia precursor assays demonstrated that p190RhoGAP is a resident protein of the invadopodia core (Fig. 4a and b), and p190RhoGAP depletion phenocopied the ECM degradation deficiency observed with TC10 depletion (Fig. 4c and Supplementary Fig. S9a and b). The overexpression of a dominant-negative version of p190RhoGAP, which lacks the ability to activate GTP hydrolysis by Rho GTPases, also strongly inhibited ECM degradation (Fig. 4c). Moreover, invadopodia lifetimes were not impacted by p190RhoGAP depletion (Fig. 4d), which suggested that p190RhoGAP is associated with the functional aspects of invadopodia, rather than the structural aspects, similar to TC10. p190RhoGAP depletion was associated with a reduction in the TC10 proportion observed in the invadopodia core, accompanied by an increase in the proportion of TC10 observed in the lateral aspects of invadopodia (Fig. 4e), which suggested that the flux of TC10 through the ring-like region and into the invadopodia core was significantly attenuated by p190RhoGAP depletion. In line with our hypothesis that TC10 activity might be affected by perturbations in p190RhoGAP activity, p190RhoGAP depletion also impacted the surface presentation of MT1-MMP in a manner similar to that observed for TC10 depletion (Fig. 4f). These findings indicated that p190RhoGAP plays an important role in the regulation of invasive functions associated with invadopodia.

**Figure 4:**
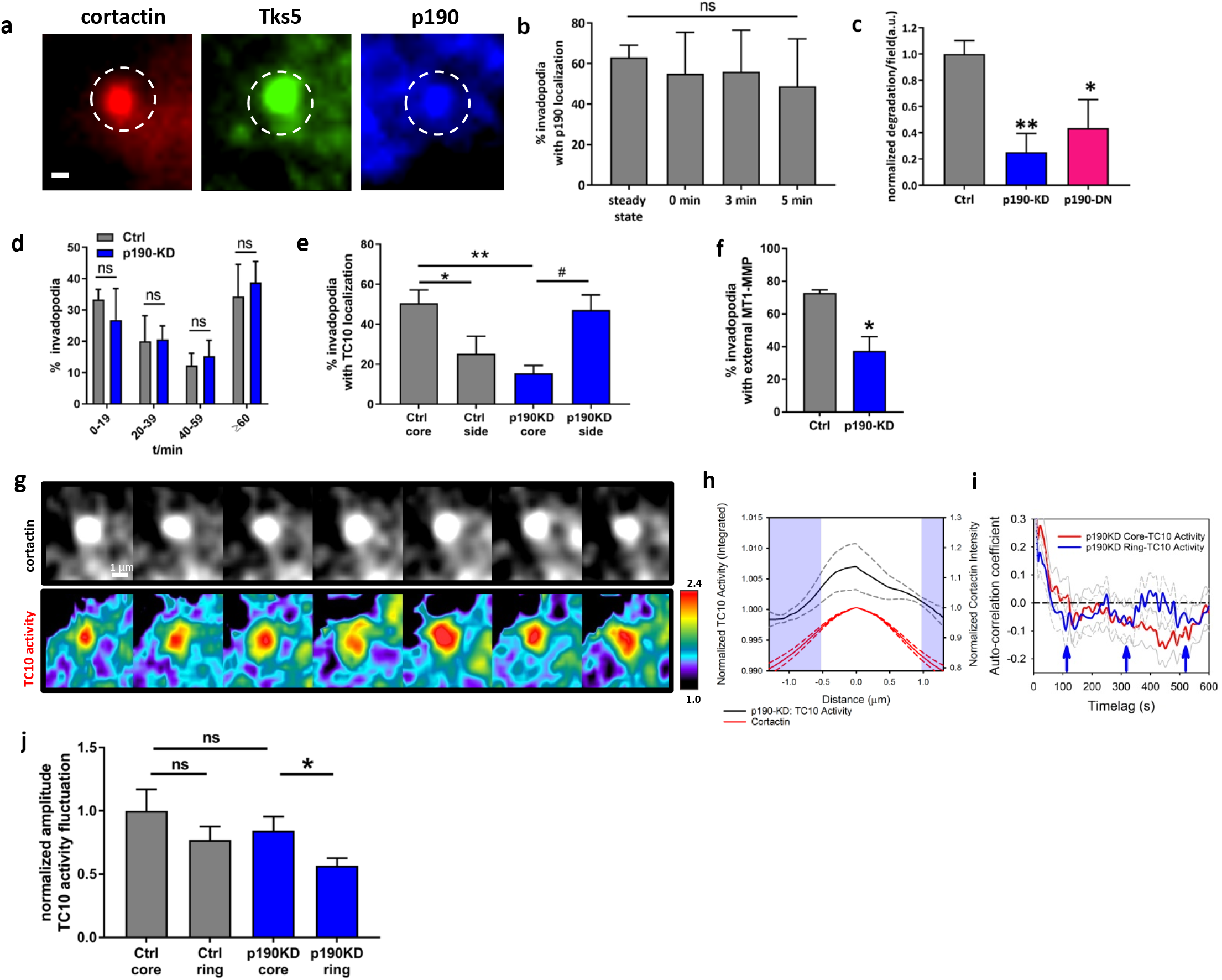
p190RhoGAP impacts invadopodia function by targeting TC10. **a.** Representative, immunostaining of endogenous p190RhoGAP at the invadopodia core, in MTLn3 cells. White bar = 1-μm. **b.** The percentage of invadopodia with p190RhoGAP localization in MTLn3 cells, following starvation and EGF stimulation (5 nM) for the indicated times. The steady state percentage of invadopodia with p190RhoGAP localization in MTLn3 cells in serum is also shown. Student’s t-test, two-tail analysis: ns p=0.7220, steady state versus 0 min; p=0.9730, 0 min versus 3 min; p=0.8520, 0min versus 5 min; n=3 experiments; shown with SEM. **c.** Matrix degradation from MTLn3 cells transfected with control siRNA (Ctrl, gray) or siRNA against p190RhoGAP (KD, blue), and the overexpression of a catalytically dead p190RhoGAP dominant-negative mutant (DN, pink). Results are normalized to the Ctrl. p190RhoGAP depletion characterization and efficiency evaluations are shown in Supplementary Figure S8. Student’s t-test, two-tail analysis: ** p=0.006336; n=3 experiments; shown with SEM; * p= 0.04141; n=4 experiments; shown with SEM. **d.** Invadopodia lifetime assay in MTLn3 cells transfected with Ctrl versus p190RhoGAP siRNA. Student’s t-test, two-tail analysis: ns, p= 0.5646 for 0-19 min; p=0.9495 for 20-39 min; p=0.6673 for 40–59 min; p=0.7356 for >60 min; n=3 experiments, shown with SEM. **e.** The localization of TC10 in MTLn3 cells transfected with Ctrl or p190RhoGAP siRNA. Student’s t-test, one-tail analysis: ** p=0.005126; n=4 experiments; shown with SEM; Student’s t-test, paired one-tail analysis: * p= 0.033138; # p=0.03791; n=4 experiments; shown with SEM. **f.** The percentage of invadopodia with extracellular, endogenous MT1-MMP localization from among 231 cells, transfected with either Ctrl or p190RhoGAP siRNA. Student’s t-test, paired one-tail analysis: * p=0.02492; n=3 experiments; shown with SEM. **g.** Representative example images of TC10 biosensor activity at an invadopodium in MTLn3 cells transfected with p190RhoGAP siRNA. The invadopodium is denoted by the cortactin fluorescence signal. h. The line scan analysis of the intensity distributions across invadopodia for TC10 activity integrated over time, showing the p190RhoGAP-depleted condition, together with the normalized cortactin trace. The blue-shaded regions indicate significant (p < 0.05; Student’s t-test, one-tailed; n = 25 invadopodia from 12 cells over 3 experiments; For p-value distributions, see Appendix 1) differences from the TC10 activity intensity at the center of the invadopodia core at 0.0 μm. TC10 activity line scans were normalized to the position at the ring-like region surrounding the invadopodia core, as determined and shown in Figure 3g. The cortactin intensity was normalized at the center position, taken as the maximal intensity location along the line scans. **i.** Autocorrelation functions for the fluctuation of TC10 activity when p190RhoGAP is depleted in the invadopodium core (red) versus the ring-like region (blue) around the invadopodium core. The gray lines (solid: core; dashed: ring) indicate the 95% confidence intervals around the mean. The autocorrelation function in the core of invadopodia does not appear to inflect after the first zero-crossing (no periodicity). The autocorrelation function in the ring-like region has repeating inflection patterns that cross zero several times at a measurable periodicity (inflection points are indicated with blue arrows) of approximately 229 ± 28 seconds. N = 18 invadopodia core and ring measurements from 10 different cells in 3 experiments. **j.** The absolute values of the amplitude of fluctuation in the TC10 biosensor activity in the core versus the ring-like region around invadopodium core. The data are normalized to the core fluctuation amplitudes in the WT condition (first bar). Student’s t-test, two-tail analysis: ns p= 0.2546 (WT core versus WT ring), ns p= 0.4415 (WT core versus p190KD core), * p= 0.03771 (p190KD core versus p190KD ring); n = 29 invadopodia core and ring measurements for WT, from 19 different cells, in 8 experiments, and n = 18 invadopodia core and ring measurements for p190KD from 10 different cells in 3 experiments; shown with SEM.

Next, we attempted to determine the functional impacts of p190RhoGAP on TC10 activity at invadopodia. We used our FRET biosensor to monitor changes in TC10 activity following p190RhoGAP depletion at invadopodia. Live-cell imaging of TC10 activity in p190RhoGAP-depleted cells revealed strong fluctuations in the activity patterns at and surrounding the invadopodia (Fig. 4g). We integrated the TC10 activity over time at invadopodia and performed a line scan analysis, which showed that the time-integrated activity of TC10 was significantly elevated within the invadopodia core in cells with p190RhoGAP depletion (Fig. 4h). To characterize the temporal fluctuations of TC10 activity at invadopodia under p190RhoGAP depletion conditions, we used the autocorrelation analysis. We found that the core-associated TC10 activity dynamics were stochastic and lacked periodicity, similar to WT conditions (Figs. 4i and 3g). Interestingly, we found that p190RhoGAP depletion produced a small periodic oscillation in TC10 activity in the ring-like region surrounding the core (Fig. 4i). The characteristic periodicity observed for the TC10 activity fluctuation in the ring-like region of invadopodia was approximately 5 min, which is within a similar order of magnitude as previously determined invadopodium core protein fluctuation rates, including those for cortactin and neural Wiskott-Aldrich syndrome protein (N-WASP)^36^. These observations suggest the transient, bulk flux of TC10 activity through the ring-like region surrounding the invadopodia core in the absence of GTP hydrolysis under p190RhoGAP depleted conditions. This may reduce the degree of freedom of activity modulation and could produce a periodic function characterizing only the bulk flux of active TC10 through the ring-like region in transit into the core (Supplementary Fig. S9c). The absolute value of the fluctuation amplitude was significantly reduced in the ring-like region compared with that in the core when p190RhoGAP was depleted (Fig. 4j), supporting the hypothesis that p190RhoGAP depletion resulted in reduced degrees of freedom. This observation is consistent with the hypothesis that p190RhoGAP regulates the ability of TC10 to hydrolyze GTP as it transits from the ring-like region into the invadopodia core, where p190RhoGAP primarily resides.

In addition to targeting TC10, RhoC GTPase activity is also directly impacted by p190RhoGAP ^35,39,51^. Previously, the inactivation of RhoC was shown to increase ECM degradation at invadopodia via a mechanism associated with changes to the invadopodia structural cohesion through the RhoC-Rho kinase 1 (ROCK)-LIM kinase (LimK)-cofilin-phosphorylation pathway^39^. p190RhoGAP depletion would, therefore, be expected to cause the overactivation of RhoC^51^. We generated a fast-cycling, constitutively activated RhoC (F30L) that contained a set of analogous GAP-binding deficiency mutations (E93H and N94H)^52^. The overexpression of this mutant RhoC would allow the effects of p190RhoGAP depletion to be mimicked for RhoC without affecting the ability of native p190RhoGAP to target TC10. The overexpression of this RhoC mutant impacted ECM degradation but had no significant effects on the total number of steady-state invadopodia or the relative MT1-MMP localization at invadopodia (Supplementary Fig. S10). However, as expected, based on the structural effects of RhoC on invadopodia, we observed a shift in the invadopodia lifetimes, favoring structures with faster turnover rates and reducing those with longer lifetimes (Supplementary Fig. S10). An increase in the population of invadopodia that turnover rapidly is associated with structural instability, which impacts invadopodia maturation and reduces ECM degradation. Thus, RhoC activation affects ECM degradation, likely through structural effects rather than vesicular targeting or fusion defects. These observations indicate the divergent roles of TC10- and RhoC-driven pathways at invadopodia that are simultaneously regulated by a single upstream regulator p190RhoGAP.

### Tyrosine phosphorylation of p190RhoGAP is required for ECM degradation

The phosphorylation of p190RhoGAP by the non-receptor tyrosine kinase Arg promotes the binding of p190RhoGAP to p120RasGAP and initiates the recruitment of the p190:120 complex to the cell periphery, where the GAP activity of p190RhoGAP for RhoGTPases is potentiated^53,54^. Arg is activated by β1 integrin binding during invadopodia maturation^30^. Arg phosphorylates p190RhoGAP at Y1105, in the RasGAP-binding region, and Y1087, which stabilizes the interaction between p190RhoGAP and p120RasGAP^53^. Therefore, we determined the phosphorylation status at Y1105 of p190RhoGAP at invadopodia and examined how phosphorylation activity affected the p190RhoGAP-mediated regulation of invadopodia functions. Approximately 90% of steady-state invadopodia contained Y1105-phosphorylated p190RhoGAP, which was found both in the core compartment and occasionally on the lateral sides of invadopodia (Fig. 5a and b), which agrees with a previous study that showed that only the active, phosphorylated form of p190RhoGAP was recruited to the plasma membrane to act on RhoGTPases^53,54^. The phosphorylation of p190RhoGAP at Y1105 is time-dependent following EGF stimulation to induce the synchronous formation of invadopodia precursors (Fig. 5c), mirroring the previously described Arg-mediated phosphorylation events at the invadopodium core following EGF stimulation^55^. In line with the tyrosine-phosphorylated status of p190RhoGAP, we observed a strong colocalization between p120RasGAP, p190RhoGAP, and TC10 at invadopodia, either on the side of the invadopodia or in the core, overlapping with the cortactin signal (Fig. 5d and e). The colocalization of p190RhoGAP at the invadopodia core was significantly altered by the expression of a competitive inhibitor of p190:120 binding^53,54^, shifting to a lateral localization pattern (Fig. 5f). The presence of this competitive inhibitor also reduced ECM degradation (Fig. 5g) without impacting the number of steady-state invadopodia (Fig. 5h). Phosphorylation-deficient p190RhoGAP point mutations, in which the two phosphorylated tyrosines were replaced with phenylalanines (Y1105F and Y1087F), strongly impacted ECM degradation, similar to the effects observed in response to TC10 and p190RhoGAP depletion and the overexpression of the p190:120 competitive binding inhibitor (Fig. 5i). However, these p190RhoGAP point mutations only affected the functional aspects of invadopodia without affecting the number of steady-state invadopodia (Fig. 5j). These observations indicated that p190RhoGAP is targeted to the invadopodia core through tyrosine phosphorylation, which promoted p120RasGAP binding.

**Figure 5:**
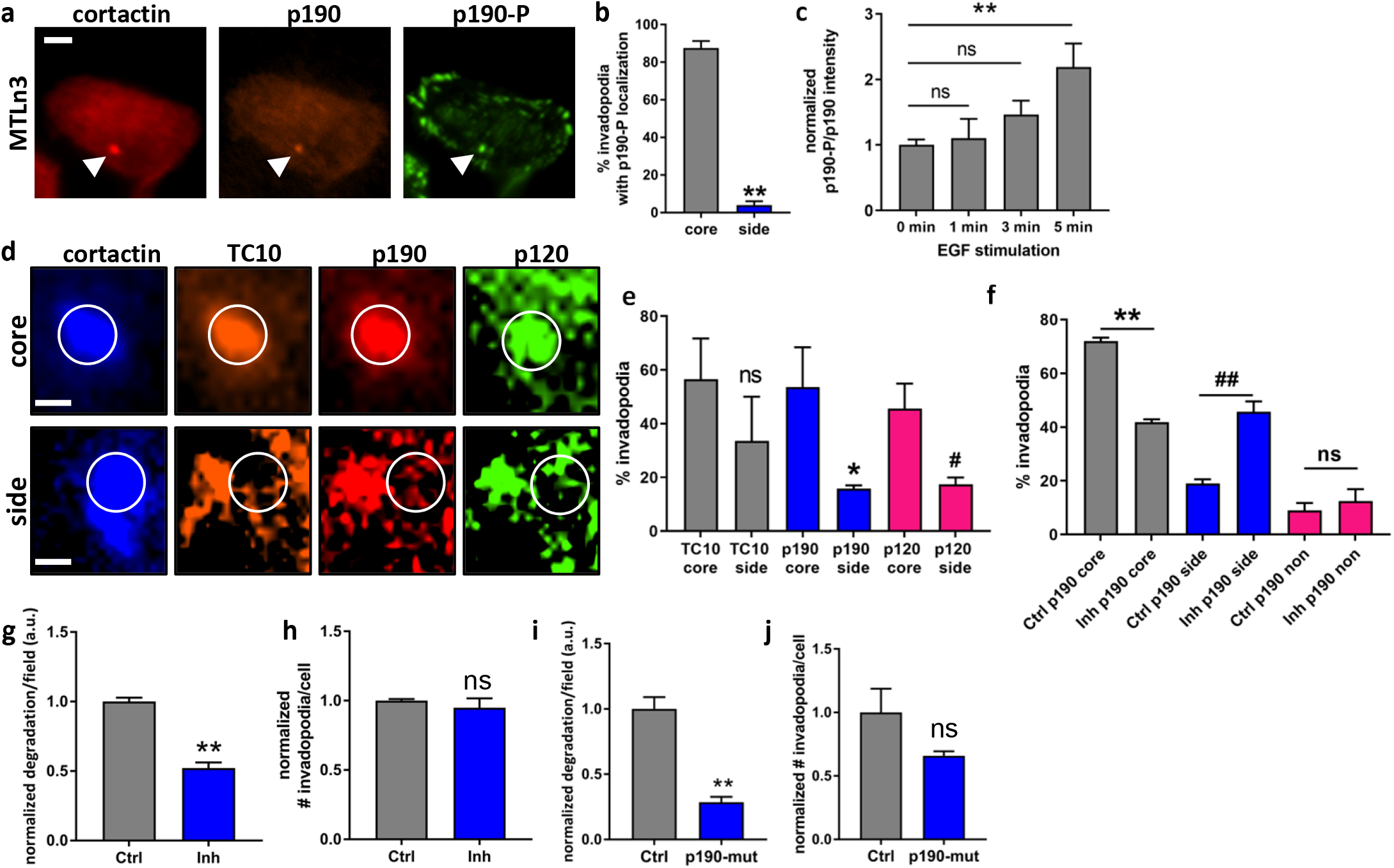
Tyrosine phosphorylation of p190RhoGAP is required for matrix degradation. **a.** Representative, immunostaining for endogenous p190RhoGAP and Y1105-phosphorylated p190RhoGAP, co-expressed with mtagRFP-T-cortactin as an invadopodia marker. White bar = 5-μm. **b.** Quantification of the percentage of invadopodia that were positive for phosphorylated p190RhoGAP colocalization. Student’s t-test, two-tail analysis: ** p=0.0001518; n=3 experiments, shown with SEM. **c.** The ratio of phosphorylated p190RhoGAP to total p190RhoGAP at invadopodia in MTLn3 cells during invadopodia precursor formation, induced by 5 nM EGF treatment following starvation for the indicated times. Results are normalized to the ratio at 0-min. Student’s t-test, two-tail analysis: “ns” p=0.742460426 for 0 min – 1 min; p=0.120533868 for 0 min – 3 min; and ** p=0.007470 for 0 min – 5 min; n=3 experiments, shown with SEM. **d**. Representative, immunostaining of endogenous p190RhoGAP and p120RasGAP at invadopodia, shown together with the co-expression of fluorescent protein-tagged WT TC10 and cortactin. White bar = 1-μm. **e.** Quantification of endogenous p190RhoGAP and p120RasGAP localization at either the core or the side of invadopodia. Student’s t-test, two-tail analysis: “ns” p= 0.3618 for TC10 core vs. side; * p=0.01146 for p190 core vs. side; # p=0.04271 for p120 core vs. side; n=3 experiments, shown with SEM. **f.** Quantification of the change in p190RhoGAP localization upon overexpression of the p190:p120 competitive binding inhibitor. Student’s t-test, two-tail analysis, ** p=0.00006365; ## p=0.002991; “ns” p=0.5417; n=3 experiments, shown with SEM. **g.** Quantification of matrix degradation by MTLn3 cells when the p190:p120 competitive binding inhibitor was overexpressed. Results are normalized to the Ctrl where only the fluorescent protein was overexpressed. Student’s t-test, two-tail analysis: ** p= 0.0005644; n=3 experiments, shown with SEM. **h.** The number of steady-state invadopodia in MTLn3 cells when the p190:p120 competitive binding inhibitor was overexpressed. Results are normalized to the Ctrl where only the fluorescent protein was overexpressed. Student’s t-test, two-tail analysis: “ns” p=0.5016; n=3 experiments, shown with SEM. **i.** Quantification of the matrix degradation by MTLn3 cells when a Y1105/1087 phosphorylation-deficient mutant version of p190RhoGAP was overexpressed. Results are normalized to the Ctrl where only the fluorescent protein was overexpressed. Student’s t-test, two-tail analysis: ** p= 0.002018; n=3 experiments, shown with SEM. **j.** The number of steady-state invadopodia in MTLn3 cells when a Y1105/1087 phosphorylation-deficient mutant version of p190RhoGAP was overexpressed. Results are normalized to the Ctrl where only the fluorescent protein was overexpressed. Student’s t-test, two-tail analysis: ** p=0.1475; n=3 experiments, shown with SEM.

### TC10 is required for cancer cell metastasis *in vivo*

We attempted to determine the functional relevance of TC10 signaling for the process of breast cancer cell invasion and metastasis. We first investigated the ability of tumor cells to invade through the ECM using an *in vitro* invasion assay, in which cultured tumor cells respond to serum stimulation by migrating through a Matrigel-coated filter ^28^. Compared with the control siRNA-treated condition, TC10 depletion significantly impacted the ability of MTLn3 cells to invade through Matrigel-coated filters in an *in vitro* invasion assay (Fig. 6a), which was expected due to the reduced ECM degradation capacity and the reduced MT1-MMP presentation at invadopodia. Moreover, p190RhoGAP depletion in MTLn3 cells also significantly attenuated the ability of these cells to invade, phenocopying TC10 depletion (Fig. 6b). To examine whether TC10 is required for breast tumor metastasis in a mouse model, we generated a clustered regularly interspaced short palindromic repeat (CRISPR)/CRISPR-associated 9 (Cas9)-driven TC10 knockout cell line in MTLn3 cells that stably express EGFP. We chose the CRISPR/Cas9 knockout cell population expressing single-guide RNA (sgRNA) #4, which showed the strongest TC10 knockout efficiency in a stable cell population (Fig. 6c). TC10 knockout cells showed significant ECM degradation defects (Fig. 6d) but no changes in the total number of steady-state invadopodia, similar to the effects observed for siRNA-mediated TC10 depletion (Fig. 6e). ECM degradation deficiencies in the TC10 knockout cells could be fully rescued by the overexpression of WT TC10 (Fig. 6f).

**Figure 6:**
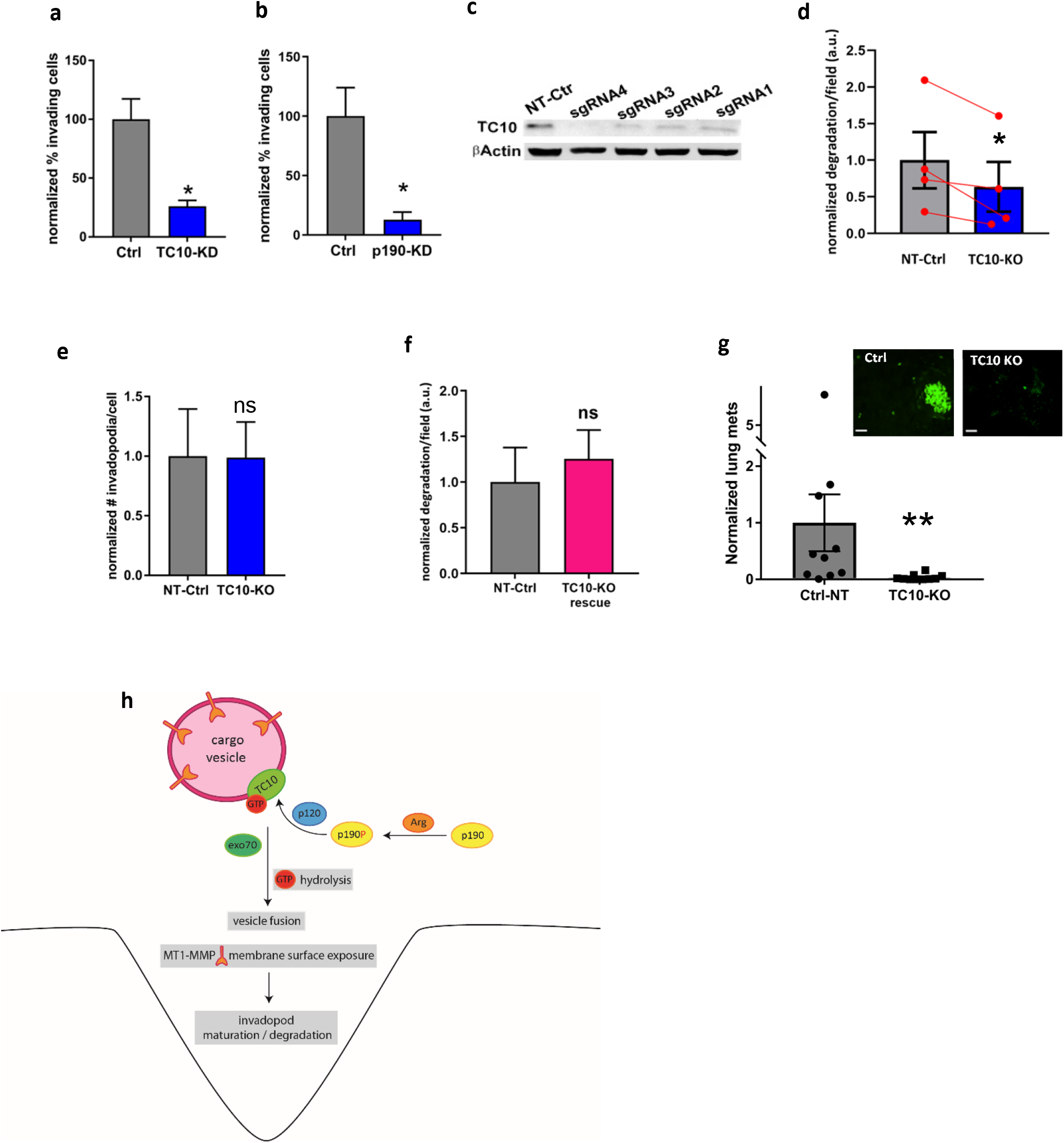
TC10 is required for cancer cell metastasis *in vivo*. **a.** The percentage of invading cells in an *in vitro* invasion assay for MTLn3 cells transfected with Ctrl or TC10 siRNA. Student’s t-test, one-tail pair-wise analysis, * p=0.03928, n=6 experiments. Error bars represent the SEM. **b.** The percentage of invading cells in the *in vitro* invasion assay using MTLn3 cells transfected with Ctrl or p190RhoGAP siRNA. Student’s t-test, two-tail analysis: * p=0.02495; n=3 experiments; shown with SEM. **c.** CRISPR/cas9 knockout populations of TC10 using 4 different sgRNA designs. Full-sized western blots are shown in Supplementary Figure S13. **d.** The matrix degradation by a TC10 knockout MTLn3 cell population using sgRNA4. Results are normalized to the NT Ctrl degradation. Student’s t-test, one-tail pair-wise analysis, * p=0.03370, n=4 experiments. Error bars represent the SEM. **e.** Functional rescue of matrix degradation by the overexpression of WT TC10 in TC10-KO MTLn3 cells. Results are normalized to the NT Ctrl degradation. Student’s t-test, one-tail pair-wise analysis, “ns” p=0.1737, n=3 experiments. Error bars represent the SEM. **f.** The number of steady-state invadopodia per cell in MTLn3 cells, with or without TC10 CRISPR/cas9 knockout. Results are normalized to the NT Ctrl. Student’s t-test, two-tail pair-wise analysis, “ns” p=0.9438, n=4 experiments. Error bars represent the SEM. **g.** The lung surface metastases of the Ctrl non-targeting or sgRNA4-TC10 knockout MTLn3 cells in the spontaneous metastasis assay, as measured by the stable co-transduction of EGFP (insets show representative fields of views). Results are normalized to the NT Ctrl. White bars, 100 μm. N = 12 mice for Ctr and n = 10 for TC10 KO. **p =0.001380 (Mann-Whitney U test, two-tail analysis). Error bars represent the SEM. **h.** A schematic model showing the pathways regulated through TC10 modulation that might impact breast cancer invasion and metastasis.

We orthotopically injected TC10 knockout cells into the mammary fat pads of 6-8-week-old female severe combined immunodeficient (SCID) mice and examined lung metastasis after the primary tumor reached 1 cm in diameter. Lung metastasis was significantly impacted in mice bearing TC10-knockout MTLn3 tumors compared with mice bearing non-targeting control tumors (Fig. 6g). Together, these results suggest that TC10 functionally impacts breast tumor dissemination and metastasis.

## Discussion

In this study, we demonstrated a new role for TC10 GTPase, a close paralog of Cdc42, in breast cancer invasion and metastasis at breast tumor invadopodia. We designed these studies to test our hypothesis that an important, previously unidentified GTPase, might regulate the invadopodia surface presentation of MT1-MMP enzymes, which are necessary for ECM degradation during tumor invasion. Our results suggested a model in which TC10 regulates MT1-MMP-containing vesicular fusion at the invadopodia membrane, which is influenced by the regulation of activity of TC10 by the upstream regulator p190RhoGAP (*Arhgap35*) at invadopodia (Fig. 6h). Our findings indicated that TC10, an important member of the Cdc42-class of Rho GTPases, plays an important role during breast cancer invasion and metastasis through the control of ECM degradative functions at invadopodia structures.

TC10 depletion resulted in significant impacts on the ability of tumor cells to degrade the ECM, associated with a decrease in the surface exposure of MT1-MMP at invadopodia. Because the overall number of steady-state invadopodia and invadopodia lifetimes were not significantly impacted by TC10 perturbations, TC10 appears to primarily play a functional role at invadopodia, without being involved in the structural aspects of invadopodia maintenance. This finding is in stark contrast with the role played by the canonical GTPase, Cdc42, at invadopodia, as the activation of Cdc42 by an upstream GEF, Vav1, has been shown to be critical for the initial formation of invadopodia precursors^56,57^. We also observed the transient but persistent activation of Cdc42 within the nascent core of invadopodia precursor structures during the assembly of the cortactin core (Supplementary Fig. S11). Our observations indicated that these close paralog GTPases play divergent roles at invadopodia structures during invadopodia assembly and function.

p190RhoGAP has been documented to primarily target Rac and Rho GTPases^58^. We previously localized p190RhoGAP at both the leading edge and the core of invadopodia in breast cancer cells^39,51^, likely due to binding with cortactin via its protrusion localization domain^43^. In these previous studies, we showed that p190RhoGAP targeted another class of RhoGTPase, RhoC, to impact actin polymerization both at the leading edge and within the core of invadopodia^39,51^. In our present work, we showed that p190RhoGAP also targets TC10 at invadopodia to regulate TC10 activity, which ultimately affects the surface presentation of MT1-MMP and ECM degradation. Moreover, we previously showed that Rac1 activity was attenuated within the invadopodia core and that the subsequent activation of Rac1 was critical for the regulation of invadopodia structural turnover^41^. Although we did not identify a specific GAP involved in Rac1 regulation in that study, the over-activation of Rac1 reduced the total number of invadopodia, whereas the depletion of p190RhoGAP in the present study did not change the total number of steady-state invadopodia or affect invadopodia lifetimes compared with control conditions. These observations suggest that a different set of signaling pathways are likely responsible for regulating Rac1 activity at invadopodia, separate from the p190RhoGAP-TC10 pathway. RhoGAPs have been shown to be relatively promiscuous, interacting with many GTPases^59^; therefore, the multi-specificity of p190RhoGAP at invadopodia is likely to coordinate the signaling regulation of a number of RhoGTPases, including RhoC and TC10 but not Rac1.

Interestingly, the depletion of p190RhoGAP led to an accumulation of TC10 at the lateral side of invadopodia, while the fraction of TC10 occupying the core of invadopodia was significantly reduced. This observation suggests that the flux of TC10-containing vesicles into the invadopodia core region could be impacted when p190RhoGAP is depleted. Corroborating this observation, a complete shutdown of vesicular flux has been previous observed when another GTPase important for the regulation of vesicular trafficking, (Arf6) was perturbed ^38,60^. Here, the GTP-hydrolysis by TC10 was perturbed through the depletion of p190RhoGAP. This perturbation of p190-TC10 signaling node likely prevented the exocytic fusion of the vesicles at the plasma membrane of invadopodia core and significantly attenuated the flux of the vesicles containing TC10 into the invadopodia core compartment. Importantly, those TC10 that were still able to transport into the invadopodia core compartment when p190RhoGAP was depleted, showed significantly elevated activity, pointing to the lack of p190RhoGAP-action on that population of TC10.

During the p190RhoGAP activation cascade^61^, the non-receptor tyrosine kinase Arg phosphorylates p190RhoGAP tyrosines 1105 and 1087^53^. The phosphorylation of these two sites is dependent on β1 integrin activation and is important for the formation of a complex between p190RhoGAP and p120RasGAP, which promotes the appropriate localization and GAP activity of p190RhoGAP toward RhoGTPases during cell adhesion^53,62,63^. In the present study, we showed that the phosphorylation of Y1105 is time-dependent following EGF stimulation and mirrors the previously reported Arg-mediated phosphorylation dynamics at the invadopodia core following EGF stimulation^64^. Furthermore, the expression of a phosphorylation-deficient p190RhoGAP mutant strongly impacted ECM degradation. The expression of a competitive inhibitor, based on the SH2-SH3-SH2 motif sequence in p120RasGAP, which is important for binding to phospho-tyrosine 1105 in p190RhoGAP^53,54^, significantly impacted both ECM degradation and the localization of p190RhoGAP at the invadopodia core. In line with the important role played by p120RasGAP binding on the control of p190RhoGAP localization and function, p120RasGAP depletion also resulted in reduced ECM degradation by invadopodia (Supplementary Fig. S12). However, we also noted a small but significant reduction in the total number of invadopodia when p120RasGAP was depleted (Supplementary Fig. S12). This observation may indicate that p120RasGAP might also play a role in invadopodia assembly or the maintenance of structural components, possibly associated with its documented role during β1/2 integrin recycling^65^. Changes in integrin recycling mechanisms could alter the availability of functional integrins at the cell surface, which could potentially impact invadopodia stability and turnover. Our findings indicated the importance of the localization and functional mechanisms of the p190:p120 GAP-signaling complex at the invadopodia core, which target GTPases, including TC10, to regulate invadopodia functions.

The functional consequences of TC10 activity regulation and the associated effects on ECM degradation, cell invasion, and metastasis were underscored by our findings from the *in vitro* invasion and *in vivo* metastasis assays. Although the initial growth rates of primary tumors seeded using CRISPR/Cas9 TC10-knockout MTLn3 cells were similar to primary tumors seeded using non-targeting control cells (data not shown), metastasis was significantly reduced. Given the reduced invasion caused by TC10 loss, we speculate that TC10 depletion resulted in strong impacts during vascular-crossing or on secondary metastatic outgrowths. A full analysis of the *in vivo* effects of TC10 loss on metastatic capability is beyond the scope of this work, but our studies suggest TC10 plays a critical role in facilitating the efficient metastatic spread of breast tumor cells within the metastatic cascade via targeting of MT1-MMP surface exposure at invadopodia.

## Acknowledgment

This work was supported by an American Cancer Society Lee National Denim Day Postdoctoral Fellowship [PF-15-135-01-CSM (S.D.)]; NIH grants [CA100324 (J.E.S.), T32GM007288 (S.P.H.M.), and R35GM136226 (L.H.)]. J.E.S. is the Betty and Sheldon Feinberg Senior Faculty Scholar in Cancer Research. L.H. is a Hirschl Career Scientist. We thank members of the Condeelis, Segall, and Cox laboratories at Albert Einstein College of Medicine for their helpful discussions.

## Author contributions

M.H., S.K.D., and L.H. conceived the project. M.H. and L.H. designed experiments. M.H., S.K.D., and L.H. performed experiments. M.H. and L.H. analyzed the results. P.V.V. and L.H. designed the biosensors and characterized the biosensors. S.P.H.M., J.E.S., and L.H. performed the metastasis assay. L.H. directed the project. M.H. and L.H. wrote and revised the manuscript. All authors reviewed the manuscript and provided feedback.

## Methods

### Cell Culture

MTLn3 cells (rat adenocarcinoma) ^66^ were cultured in Minimum Essential Medium (MEM, Corning, Corning, NY, USA) supplemented with 5% fetal bovine serum (FBS), 1% glutamine, and 100 I.U. penicillin and 100 μg/mL streptomycin (Invitrogen, Carlsbad, CA, USA), as previously described^67^. MDA-MB-231 (HTB-26, ATCC, Manassas, VA, USA) cells were cultured in Dulbecco’s modified Eagle medium (DMEM, Corning) supplemented with 10% FBS, 1% glutamine, and penicillin/streptomycin, as previously described ^38^. All cell lines were tested regularly for mycoplasma using the PCR-based assay (Stratagene, San Diego, CA, USA).

### Transfection

Plasmid transfections were performed in OptiMEM, using Lipofectamine 2000 (Invitrogen). Cells were plated at 1 × 10^5^ cells/well in a 6-well plate and incubated overnight prior to transfection. Following the manufacturer’s protocols, 2 μg of total DNA was transfected into each well of a 6-well plate. Cells were treated with the transfection mixture for 45 min, and the transfection was terminated by exchanging the medium with the normal growth medium.

### ECM Degradation assay

Alexa Fluor 405 NHS Ester (Thermo Fisher Scientific, Waltham, MA, USA) was conjugated with 0.2% porcine gelatin (Sigma-Aldrich, St. Louis, MO, USA), according to the Thermo Fisher bioconjugation protocol. Glass coverslips (25 mm, circular #1.5, Warner Instruments, Hamden, CT, USA) were coated with 0.01% poly-L-lysine for 20 min at room temperature (RT), followed by a 15 min treatment with 0.2% glutaraldehyde in phosphate-buffered saline (PBS). The Alexa Fluor 405-labeled gelatin aliquot was centrifuged at 22,000 rcf for 10 min at RT to pellet any precipitates, the supernatant wasdiluted 1:4 with unlabeled 0.2% gelatin, and maintained at 37°C. The glutaraldehyde-treated coverslips were coated with the Alexa 405-gelatin mixture for 10 min at RT, followed by a 5 min treatment with 0.2% glutaraldehyde. Then, the coverslips were incubated in 5 mg/mL NaBH4 solution for 15 min at RT and washed 3 × with PBS. The coverslips were placed in normal culture media at 37°C and 5% CO_2_ for at least 20 mins prior to cell plating. Cells were plated at a density of 1.5 × 10^5^ cells/coverslip in wells of a 6-well plate for 16 h before fixation with 1% paraformaldehyde (PFA) for 15 mins at RT. ECM degradation was measured by quantifying the mean area of non-fluorescent pixels per field, using a manual threshold in MetaMorph software (ver. 7.10.3; Molecular Devices, San Jose, CA, USA). For experiments in which a transgene was expressed in cells, only the degraded areas under the transfected cells, as identified by fluorescent protein expression, were considered.

### EGF stimulation

EGF stimulation was performed as previously described^39,41^. In brief, MTLn3 cells were starved for 4 h in L15 media containing 0.003% bovine serum albumin (BSA) at 37°C, without CO_2_, and then stimulated with 5 nM EGF (Invitrogen) for the indicated times at 37°C before fixation for 15 min at RT using 1% PFA.

### Western Blotting

Cells were lysed on ice in a buffer containing 1% NP-40, 50 mM Tris pH 7.4, 150 mM NaCl, 10 mM ethylenediaminetetraacetic acid (EDTA), 1 mM phenylmethylsulfonyl fluoride (PMSF), and 1× protease inhibitor cocktail (Sigma). The lysate was clarified by centrifugation at 22,000 rcf for 10 min at 4°C. Lysates were resolved by 8%–12% sodium dodecyl sulfate-polyacrylamide gel electrophoresis (SDS-PAGE). Proteins were transferred to polyvinylidene fluoride membranes. After blocking for at least 1 hour in 5% BSA in Tris-buffered saline containing Tween-20 (TBS-T), membranes were incubated with primary antibodies at 1:1000 dilution overnight at 4°C. Membranes were incubated with secondary fluorescently labeled antibodies (LI-COR Biosciences, Lincoln, NE, USA) at 1:10,000 dilution for 1 h at RT. Immunoblots were visualized using the Odyssey Imager (LI-COR Biosciences).

### Antibodies

TC10 (Novus, Littleton, CO, USA; 07-2151; rabbit polyclonal used at 1:500 for western blots), p190 (BD Transduction Laboratories; 610149; Clone 30/p190; mouse monoclonal), p120 (Abcam, Cambridge, UK; ab2922; Clone B4F8; mouse monoclonal), Exo70 (Santa Cruz Biotechnology; sc-365825; Clone D-6; mouse monoclonal), Vamp7 (Abcam; ab36195; Clone 158.2; mouse monoclonal), MT1-MMP-Hinge region (Millipore, Burlington, MA, USA; AB6004; rabbit polyclonal), MT1-MMP (Millipore; MAB3328; Clone LEM-2/15.8; mouse monoclonal), MT1-MMP (Abcam; ab38971; rabbit polyclonal), Cortactin (Abcam; ab3333; Clone 0.T.21; mouse monoclonal, used at 1:600), Cortactin (Abcam; ab81208; Clone EP1922Y; rabbit monoclonal), Cortactin (Santa Cruz Biotechnology; sc-30771; G-18; goat polyclonal), MYC (Cell Signaling Technology, Danvers, MA, USA; mab2278; Clone 71D10; rabbit monoclonal), FLAG (Sigma; F1804; Clone M2; mouse monoclonal), and EGFP (Roche; 11814460001; Clones 7.1 and 13.1; mixture mouse monoclonal). Unless otherwise stated, all primary antibodies were used at 1:200 dilution for immunofluorescence and 1:1,000 for western blotting.

### *In vitro* invasion assay

*In vitro* invasion assays were performed as previously described ^39^. In brief, 1.5 × 10^5^ cells were plated in the top wells of Growth Factor Reduced Matrigel-coated invasion chambers (8 μm pore size, BD Bio Coat). Media containing 5% was added to the lower chamber, and cells were allowed to invade along the serum gradient for 18 h at 37°C. The assay was fixed with 3.7% PFA for 20 min and stained with NucBlue (Invitrogen) to visualize the nuclei. When siRNA-transfected cells were used, siGLO-Red (Dharmacon, Lafayette, CO, USA) was co-transfected in the cells to identify siRNA-treated cells. The membrane was detached from the chamber and mounted on a coverslip, and 10 random fields of view were imaged across the membrane at 20× magnification on an IX81-ZDC microscope (Olympus, Tokyo, Japan). The number of invading cells was counted manually with ImageJ software by thresholding onto the nucleus, and data are reported as the means of 3 experiments for each condition.

### Invadopodia lifetime assay

MTLn3 cells were transfected with cortactin-miRFP703 and EGFP-Tks5^68^ before plating on gelatin-coated coverslips for 16 h. The cells were imaged every 2 min for 4 h on an IX81-ZDC inverted epifluorescence microscope at 60× magnification (Olympus). Invadopodia lifetimes were quantified manually for at least 30 invadopodia from at least 10 cells per condition in at least 3 experiments. Control and siRNA conditions were imaged on the same day for each experiment. Cells expressing siRNA and scrRNA were identified by co-transfection with siGLO-Red (Dharmacon).

### Immunoprecipitation and pull-down experiments

HEK293T cells were plated overnight at a density of 1 × 10^6^ cells on poly-L-lysine-coated six-well plates. The FLAG-tagged TC10 mutants and the MYC-tagged WT Exo70 expression constructs were mixed at a 1:1 ratio, and the cells were transfected using the polyethyleneimine (PEI) reagent at the optimized 2 μg DNA to 8 μL PEI ratio for each well, according to published protocols^69^. After 48 h, cells were lysed in a buffer containing 1% NP-40, 20 mM Tris HCl, pH 7.4, 137 mM NaCl, 10 mM MgCl2, 1 mM PMSF, and 1× protease inhibitor cocktail (Sigma-Aldrich). Lysates were clarified by centrifugation at 22,000 rcf for 10 min at 4°C. After removing an “input fraction, lysates were mixed with protein A/G agarose beads (Pierce, Waltham, MA, USA) conjugated to antibodies against FLAG-tag (Sigma-Aldrich) or Exo70 (Santa Cruz, Dallas, TX, USA), at a concentration of 2 μg antibody per sample, and incubated overnight at 4°C with gentle rocking. Samples were washed 3× in lysis buffer, mixed with 5× gel loading buffer, and boiled for 5 min at 99°C prior to loading separation by SDS-PAGE for western blotting analysis.

Biosensor pull-downs were performed using purified PAK1-PBD-agarose beads, as previously described^41^. To prepare the glutathione (GSH)-agarose beads, 72 mg of GSH-agarose (Sigma-Aldrich) was resuspended in 10 ml sterile water and incubated at 4°C for 1 h. The suspension was briefly centrifuged, and the pellet was washed three times with sterile water, followed by washing two times in a resuspension buffer (50 mM Tris, pH 8.0, 40 mM EDTA, and 25% sucrose). The washed GSH-agarose slurry was resuspended in 1 ml of resuspension buffer. To generate GST-PAK1-PBD, pGEX-PBD (a gift from G. Bokoch^70^) was transformed into BL21(DE3)-competent bacteria (Agilent Technologies, Santa Clara, CA, USA) and grown in a shaker flask at 225 rpm and 37°C until an optical density of 1.0 at 600 nm was achieved. Protein synthesis was induced by the addition of 0.2 mM Isopropyl β-d-1-thiogalactopyranoside (IPTG), and the flask was immediately chilled to RT and incubated at 225 rpm and 24°C overnight. The next day, bacteria were pelleted and resuspended in 20 ml resuspension buffer containing 1 mM PMSF, 1× protease inhibitor cocktail (Sigma-Aldrich), and 2 mM β–mercaptoethanol and rotated on a Nutator for 20 min at 4°C. After incubation, 8 ml detergent buffer (50 mM Tris, pH 8.0, 100 mM MgCl_2_, and 0.2% [wt/vol] Triton X-100) was added, and the mixture was incubated at 4°C for 10 min on a Nutator. After incubation, the mixture was ultrasonicated (4× cycles of 30-s ultrasonication followed by 1 min rest on ice) and centrifuged at 22,000 rcf for 45 min at 4°C. The supernatant was transferred to a 50-ml tube, and 1 ml previously prepared GSH-agarose beads were added and incubated at 4°C for 1 h on a Nutator. The beads were then pelleted by a brief centrifugation step and washed four times with wash buffer (50 mM Tris, pH 7.6, 50 mM NaCl, and 5 mM MgCl2) followed by resuspension in 500 μl of 50:50 glycerol/wash buffer. Aliquots of this mixture at 50 μl aliquots were stored at −80°C until use. For pull-down experiments, HEK293T cells were transfected and lysed as described above. Lysates were clarified by centrifugation at 22,000 rcf for 10 min at 4°C. After removing an “input” fraction, lysates were incubated with PAK1–PBD-conjugated agarose beads for 1 h at 4°C, washed 3× in lysis buffer, resuspended in final sample buffer, and analyzed by western blotting. Incubation with Ponceau S solution (Sigma-Aldrich) was used to visualize GST-PAK1-PBD to control for equal loading. Anti-GFP (mouse; 11814460001; clones 7.1 and 13.1 mix; Roche, Basel, Switzerland) antibody was used to detect the TC10 biosensor or fluorescently tagged TC10 protein.

### Generation of a TC10-knockout cell line using CRISPR-Cas9

Four different 20-nt guide sequences for TC10 were selected using the online CRISPR Design Tool (http://tools.genome-engineering.org) against rat TC10 GTPase. Sequences for the primer pairs are as follows: sgRNA 1: 5’-CACCGCGTAGTGGTCGAAGACAGT-3’ and 5’-AAACACTGTCTTCGACCACTACGC-3’; sgRNA 2: 5’-CACCGTGCGTAGTGGTCGAAGACAG-3’ and 5’-AAACCTGTCTTCGACCACTACGCAC-3’; sgRNA 3: 5’-CACCGAGGTACTGCTTGCCCCCCA-3’ and 5’-AAACTGGGGGGCAAGCAGTACCTC-3’; and sgRNA 4: 5’-CACCGGGGGGCAAGCAGTACCTCT-3’ and 5’-AAACAGAGGTACTGCTTGCCCCCC-3’. A negative control NT1 with the sequence 5’-GCGAGGTATTCGGCTCCGCG-3’ was also used, which was based on a negative control sequence from the GeCKOv2 Mouse Library Pool A^71^. sgRNAs were cloned into the pLentiCRISPR v2 plasmid ^71,72^ by digestion with *Bsm*BI (New England Biololabs, Ipswich, MA, USA). pLentiCRISPR v2 was a gift from F. Zhang (Massachusetts Institute of Technology, Cambridge, MA, USA; 52961; Addgene #52961, Watertown, MA, USA). The GP2-293 cell line (Takara Bio Inc., Shiga, Japan) was used to produce the lentivirus by co-transfection with pVSVg, gag-pol, rev, and tat vectors (Takara Bio Inc.). MTLn3 cells were infected with the lentivirus containing the four TC10-targeting sgRNAs or the NT1 control sgRNA and were cultured as described in the Cell Culture section. Transduced cells were selected for the stable incorporation of the CRISPR/Cas9 vector by puromycin treatment (2 μg/ml). CRISPR knockout efficiency was assessed by western blotting against TC10 (Fig. 6B). An efficient knockout population was achieved with sgRNA4, which was used for subsequent experiments.

### Expression cDNA constructs

Cortactin–mtagRFP-T^64^ and EGFP-Tks5^68^ have been previously described. To generate cortactin-miRFP703, mtagRFP-T was replaced with miRFP703^73^. MT1-MMP-GFP, as previously described^74^.

Full-length human p120RasGAP1 was a gift from D. Esposito (Addgene #70511). To construct the competitive inhibitor of p190:p120 interaction, the sequence for amino acids 180 to 474 of the human p120RasGAP1 was PCR amplified, based on the sequence homology to Rat p120RasGAP1, as published previously^54^. The following primer pair was used: 5’-GGAATGTTAAGCAATGGATCCTGGTATCACGGAAAACTTGACAGAAC-3’ and 5’-CGAGTACAAGTAATTCATCTCGAGCTAAATGTTTTTATAAAAGGCATCCTTTG-3’.

The PCR amplified fragment was digested with *Bam*HI and *XhoI* and ligated into the pTriEX-4 backbone at *BamHI/XhoI* sites. A codon-optimized mScarlet ^75^ fluorescent protein was synthesized (Genewiz, South Plainfield, NJ, USA) with an upstream *NcoI* site and a downstream 10 amino acid linker: GSGSGSGSGG (5’-GGCAGCGGCTCCGGGAGCGGGTCCGGAGGC-3’), followed by a *Bam*HI site, and inserted into the pTriEX-4 vector containing the 2-3-2 fragment at the *NcoI/BamHI* sites. To produce the pTriEX-mtagBFP2 version of the p190:p120 competitive inhibitor construct, the mScarlet fluorescent protein was restriction digested with *NcoI/BamHI*, and mtagBFP2^76^ was 2-step PCR-amplified using the following primer pairs: 5’-GCAATATAATGAATACCATGGTGTCTAAGGGCGAAGAGCTGAT-3’ and 5’-ACCCGCTCCCGGAGCCGCTGCCATTAAGCTTGTGCCCCAGTTTGCTA-3’, followed by 5’-GCAATATAATGAATACCATGGTGTCTAAGGGCGAAGAGCTGAT-3’ and 5’-GGTAATAAGTATATCGGATCCGCCTCCGGACCCGCTCCCGGAGCCGCTGCCATT-3’, to encode the 10 amino acid linker GSGSGSGSGG (5’-GGCAGCGGCTCCGGGAGCGGGTCCGGAGGC-3’) followed by a *Bam*HI site. The 2-step PCR-amplified fragment was digested with *Nco*I and *Bam*HI and ligated into the pTriEX backbone containing the competitive inhibitor fragment.

Full-length p190RhoGAP-A (mouse) was previously published^39^. P190RhoGAP-A mutants were produced through PCR-based site-directed mutagenesis using the Quikchange kit (Stratagene, San Diego, CA). For the Y1087F mutation, the primer pair: 5’-GGATGGATTTGATCCTTCTGACTTCGCAGAGCCCAT-3’ and 5’-ATGGGCTCTGCGAAGTCAGAAGGATCAAATCCATCC-3’ was used. For the Y1105F mutation, the primer pair: 5’-CAAGGAATGAGGAAGAAAACATATTCTCAGTGCCCCAC-3’ and 5’-GTGGGGCACTGAGAATATGTTTTCTTCCTCATTCCTTG-3’ was used. For the R1283A (catalytically-dead/dominant-negative) mutation, the primer pair: 5’-GCACTGAAGGCATCTACGCGGTCAGTGGAAACAAGT-3’ and 5’-ACTTGTTTCCACTGACCGCGTAGATGCCTTCAGTGC-3’ was used. To produce a fluorescent protein-tagged p190RhoGAP-A, the following PCR primers were used:

5’-GCATATATTAAGCAATCAAGAATTCATGGCAAGAAAGCAAGATGTCCGAA-3’ and 5’-GGTTTAAATATAGCATATACTCGAGCTACAGCGTGTGTTCGGCTTGGAGC-3’. The PCR fragment was digested with *Eco*RI and *XhoI* and ligated into the pTriEX backbone at corresponding *EcoRI/XhoI* sites, which contained the appropriate fluorescent protein at the N-terminal end of the multiple cloning site.

Full-length human WT TC10 GTPase cDNA was purchased from www.cDNA.org. TC10 mutants were produced through PCR-based site-directed mutagenesis using the Quikchange kit (Stratagene). For the Q75L mutation, the primer pair: 5’-GGTCATAGTCTTCCAGTCCGGCCGTGTCA-3’ and 5’-TGACACGGCCGGACTGGAAGACTATGACC-3’ was used. For the T31N mutation, the primer pair: 5’-CATGAGTAGGCAATTCTTGCCCACCGCCCCGTC-3’ and 5’-GACGGGGCGGTGGGCAAGAATTGCCTACTCATG-3’ was used. For the T49A mutation, the primer pair: 5’-TGGTCGAAGACGGCGGGCACGTACTCC-3’ and 5’-GGAGTACGTGCCCGCCGTCTTCGACCA-3’ was used. For the Y54C mutation, the primer pair: 5’-ACGCTGACTGCGCAGTGGTCGAAGACGG-3’ and 5’-CCGTCTTCGACCACTGCGCAGTCAGCGT-3’ was used. For the G26V mutation, the primer pair: 5’-TGCCCACCGCCACGTCGCCGACC-3’ and 5’-GGTCGGCGACGTGGCGGTGGGCA-3’ was used. For the F42L mutation, the primer pair: 5’-GCTATGCCAACGACGCCTTACCGGAGGAGT-3’ and 5’-ACTCCTCCGGTAAGGCGTCGTTGGCATAGC-3’ was used. For the P43L/E45V/Y46H mutations, the primer pair: 5’-ACGACGCCTTCCTGGAGGTGCACGTGCCCACCG-3’ and 5’-CGGTGGGCACGTGCACCTCCAGGAAGGCGTCGT-3’ was used. For the P43L/E45V/Y46H/T49A/Y54C mutations, the primer pair: 5’-CGACGCCTTCCTGGAGGTGCACGTGCCCGCC-3’ and 5’-GGCGGGCACGTGCACCTCCAGGAAGGCGTCG-3’ was used. To generate fusion constructs containing TC10 and fluorescent proteins or a FLAG-tag, the following primer pair was used to PCR amplify the TC10 fragment: 5’-GAGATTATTAGATGATATAGAATTCATGCCCGGAGCCGGCCGCAGCAGCAT-3’ and 5’-GCTATGCATATAATATAATCCTCGAGTCACGTAATTAAACAACAGTTTATACATC-3’. The PCR-amplified fragment was digested with *Eco*RI/*Xho*I and ligated into the pTriEX backbone, which contained the appropriate fusion tags at *Eco*RI/*Xho*I sites.

### siRNA

siRNA Smart pools for TC10, p190RhoGAP, p120RasGAP were purchased from Dharmacon/GE Healthcare (siGenome). Transfections were performed with Oligofectamine 2000 (Invitrogen) for MTLn3 cells and via electroporation, using Amaxa cell line nucleofector kit V (VACA 1003, Lonza, Basel, Switzerland), for MDA-MB-231 cells. To monitor the transfection efficiency, siGLO-Red (Dharmacon) was co-transfected, according to the manufacturer’s protocols. Knockdown was assessed, and subsequent assays were performed at 48 h (MTLn3) or 72 h (MDA-MB-231) after transfection.

### TC10 biosensor

A FRET biosensor for TC10 was constructed based on the previously published Rac-type, single-chain, genetically encoded biosensor backbone system^41^. Briefly, WT and mutant human TC10 GTPase sequences were PCR-amplified using the primer pair: 5’-GAGATTATTAGATGATATAGAATTCATGCCCGGAGCCGGCCGCAGCAGCAT-3’ and 5’-GCTATGCATATAATATAATCCTCGAGTCACGTAATTAAACAACAGTTTATACATC-3’ and restriction digested with *Eco*RI and *Xho*I. The digested fragments were ligated into the pTriEX-4 vector containing the Rac1 FRET biosensor backbone^41^ at the *EcoR*I/*Xho*I sites to exchange the Rac1 GTPase sequence for the TC10 GTPase fragments. This sensor backbone was previously codon-optimized with synonymous modifications^77^ to improve the stability and expression fidelity of the biosensor in target cells. To generate the retroviral vector containing the biosensor in the tet-inducible system, the pRetro-X vector system (Clontech, Mountainview, CA, USA) was used. Briefly, pRetro-X-puro (Clontech) was modified by inserting a Gateway destination (-DEST) cloning cassette (Invitrogen) into the multiple cloning site. The pTriEX-TC10 biosensor was restriction digested using *Nco*I and *Xho*I to extract the TC10 biosensor as a full-length cassette, which was then ligated into the pENTR-4 vector (Invitrogen) at *Nco*I/*Xho*I sites. The pENTR-TC10 biosensor was then processed for Gateway cloning, together with the pRetro-X-Puro-DEST vector, using LR Clonase II (Invitrogen), following the manufacturer’s protocols. The resulting pRetro-X-Puro-TC10 biosensor was used to produce the retrovirus used to infect cells to produce stable/inducible tet-OFF biosensor cell lines, as previously described^44^.

### Microscopy imaging

MTLn3 or MDA-MB-231 cells were plated at a cell density of 1.5 × 10^5^ on gelatin-coated glass coverslips. For fixed-cell imaging, cells were fixed for 15 min with 1% PFA in PBS and processed for immunofluorescence 16 hours after plating. A widefield imaging modality was used to obtain immunofluorescence images. For colocalization analyses, z-stacks were imaged using 0.2-μm z-steps for 26 steps, centered on the in-focus plane, and the resultant z-stacks were deconvolved (Microvolution, Cupertino, CA, USA) to remove out of focus light. For live-cell imaging, the imaging medium was prepared by using Ham’s F12K medium, without phenol red (Crystalgen, Commack, NY, USA), and supplemented with 1× glutamine, and sparged with Argon gas for 1 min to reduce the dissolved oxygen concentration. The medium was supplemented with 5% FBS, Oxyfluor Reagent (1:100 dilution, Oxyrase Inc., Mansfield, OH, USA), and 10 mM dl-lactate (Sigma-Aldrich)^78^. Cells were imaged at 37°C in a closed chamber^44^ mounted on an inverted microscope stage. Images were acquired through a 60× magnification objective lens (UIS 60× 1.45 NA; Olympus) using a custom microscope^79^ capable of the simultaneous acquisition of FRET and mCerulean emissions through two Coolsnap ES2 cameras (Photometrics, Tucson, AZ, USA) that are mounted via an optical beam splitter and containing a T505LPXR mirror, ET480/40M for mCerulean emission, and ET535/30M for mVenus-FRET emission (Chroma Technology Corp, Bellows Falls, VT, USA). The relative intensities between the two channels were balanced by the inclusion of a neutral density filter (ND0.2 in mCerulean channel) to ensure that the range of brightness in both mCerulean and FRET channels were similar to maximize the signal to noise ratio. Cells were illuminated with a 100W Hg arc lamp through a neutral density filter to attenuate light as needed and then through an ET436/20X bandpass filter for mCerulean excitation. The main fluorescence turret of the microscope contained a 20/80 mirror (Chroma Technology) that allowed 20% of the excitation illumination to reach the specimen and 80% of the emitted light to pass through to detection. The IX81ZDC microscope was fitted with a T555LPXR longpass mirror within the internal port-switching prism holder to direct the biosensor emission channels to the left-hand side port of the microscope and direct the longer wavelengths, including the cortactin and differential interference contrast (DIC) channels, to the bottom port of the microscope. The bottom port of the microscope was fitted with a single Coolsnap HQ2 camera (Photometrics) via either FF585/29 emission filter for mtagRFP-T to detect cortactin fluorescence or an aligned linear polarizer to detect the DIC illumination. MetaMorph software (Molecular Devices) was used to control the microscope, motion control devices, and image acquisition. MetaMorph and MatLab software (ver 2011a; Mathworks, Natick, MA, USA) were used to perform image processing and data analyses, as previously described^38,41,44^. Image processing included camera noise subtraction, flatfield correction, background subtraction, image registration, ratio calculations, and correction for photobleaching^80^. In brief, camera noise images were acquired at the same exposure times as the foreground image sets but without the field illumination. This represented the camera read noise and the dark current noise and was subtracted from all subsequent foreground images. Flatfield correction involved the acquisition of cell-free fields of view with the same exposure and field illumination conditions as the foreground image sets, followed by camera noise subtracted to obtain the shading images. The camera noise–subtracted foreground images were then divided by the shading images to obtain flatfield-corrected images. A small region of interest in the background (cell-free) area was selected in the flatfield-corrected foreground image sets, and the mean gray value from such a region was subtracted from the whole field of view, calculated, and processed at each time point to obtain the background-subtracted image sets. The background-subtracted image sets were then subjected to an affine transformation based, on a priori calibration, to account for misalignments between the three cameras used for the simultaneous imaging of the FRET and mCerulean channels, plus the cortactin and DIC channels in the longer wavelengths. After the transformation, a linear X–Y registration was performed on the resulting image sets, before ratio calculations, in which the FRET image set was divided by the mCerulean channel image set. For photobleaching corrections of the ratio image set, whole-cell mean gray values were calculated at each time point and fitted to a biexponential decay model. The inverse function of the regressed model was then multiplied into the ratio image set to approximate the effect of photobleaching. For fixed-cell biosensor imaging, a single Coolsnap HQ2 camera (Photometrics) attached to the bottom port of the microscope was used, together with a 60× magnification objective lens. In this case, excitation and emission filter wheels switched appropriate filter sets, in addition to the appropriate neutral density filters, to acquire mCerulean and FRET emissions plus any other additional wavelengths, as required. For the imaging of biosensors, we adjusted the camera acquisition time duration by targeting to fill approximately 80% of the total digitization range of the charge-coupled device circuitry, to maximize the dynamic range, using excitation light intensities of 0.4–1.0 mW at the specimen plane.

### Fluorometric characterization and validation of the biosensor

The characterization of the biosensor response was performed in HEK293T cells by transiently overexpressing WT or mutant versions of the biosensor with or without the appropriate upstream regulators, as described previously^81,82^. In brief, HEK293T cells were plated overnight at 1 × 10^6^ cells/well in six-well plates coated with poly-L-lysine (Sigma-Aldrich) and transfected the following day using PEI reagent according to the published optimized procedures^69^. After 48 h transfection, cells were washed once with PBS, briefly trypsinized, and resuspended in 500 μL of cold PBS per well. Cell suspensions were stored on ice until assay. Fluorescence emission spectra were measured with a spectrofluorometer (Horiba-Jobin-Yvon Fluorolog-3MF2; HORIBA, Kyoto, Japan). The fluorescence emission spectra were obtained by exciting the cell suspension in a 500 μL quartz cuvette (Starna Cells, Atascadero, CA, USA) at 433 nm, and emission fluorescence was scanned between 450–600 nm. The background fluorescence reading of cells containing an empty vector (pCDNA3.1) was used to measure light scatter and autofluorescence and was subtracted from the data. The resulting spectra were normalized to the peak of the donor mCerulean emission intensity at 474 nm to generate the final ratiometric spectra. To validate the biosensor in cancer cells using exogenous stimulation, MTLn3 cells transiently expressing the biosensor were serum-starved for 4 h and stimulated using medium containing 5% serum or 5 nM EGF. Cells were fixed and imaged at 0, 1, 2, and 3 min after stimulation and analyzed for changes in the FRET/donor ratio.

### Biosensor activity analysis at invadopodia

A time-lapse series of the region of interest containing an invadopodium was analyzed first by producing time projections. The cortactin core location image was calculated by obtaining the median projection, over time, of the cortactin channel. TC10 activity localizations were calculated by taking the summation of intensities over time at the invadopodium region of interest in a time-lapse stack, as previously described for a different class of Rho GTPase activity measurements at invadopodia^39^. Line scans were measured and averaged over 4 perpendicular lines that were centered on the core of the cortactin spot, with each line rotationally 45 degrees apart. Line scans were normalized to the local maxima of TC10 activity at the ring-like region surrounding the invadopodia core, which was denoted by the cortactin spot. The cortactin intensity was normalized at the center position, taken as the maximal intensity location along the line scans.

For experiments measuring the frequency ratio of high biosensor activity at invadopodia during transient invadopodia formation, we identified regions of cells featuring the formation of nascent invadopodium in a time-lapse experiment under steady-state conditions. The cortactin image stack was used to identify and select an elliptical region of interest in which the invadopodium core was transiently developing. A random region was also chosen away from all cortactin spots to serve as the background, and both regions were tracked for average intensity values over the entire time course of an experiment. The regions of interest were transferred to the respective biosensor ratio data stack, and the average biosensor intensity values were also measured as a function of time. The average foreground intensities over time and the standard deviation (SD) were calculated from the data, as follows: for data corresponding to the regions with cortactin spot formation, the average and SD were calculated up to the time point at which a nascent invadopodia formation became visible; for the random background control region, the average and SD were measured for the entire duration of the time-lapse experiment. The data were then thresholded at +1.0 SD away from the mean, and any activity values above this threshold were considered to be positive biosensor activity events. The total number of positive biosensor activity events were divided by the total number of time points in the corresponding time domains (“before” or “during” invadopodia formation, as determined from the cortactin data stack), and the resulting positive activity event per time data “during” invadopodia formation were normalized against the values from “before” invadopodia formation.

### Autocorrelation analysis for periodicity

For the fluctuation analysis, a binary mask was created in MetaMorph using cortactin fluorescence intensity as a reference to designate the core of the invadopodium. Subsequently, this mask was dilated 30 pixels, and the original core was subtracted to generate a binary mask to designate the invadopodia ring-like region surrounding the core. These binary masks were used to measure the intensity in each compartment of the invadopodium. The area of the ring was based on a spatial distance of 1.74 μm radius outside of the core, which is similar to the binary mask used in a previous work^39^. To quantitatively determine the periodicity of biosensor activity fluctuations within the core of an invadopodium versus the ring surrounding the invadopodium, a time series of the ratio of intensities was measured within binary masks that were generated to target either the invadopodium core or the ring surrounding the invadopodium core. These ratio time series were analyzed using the autocorrelation function *xcov* in MatLab. The individual autocorrelation function distribution was smooth-spline fitted, pooled between all invadopodia analyzed in all cells, and the mean autocorrelation function and 95% confidence intervals were calculated by a nonparametric bootstrap method^83^. The measured temporal width to the peaks of the first side lobes after the zero-crossing was taken as the period of oscillation^45^.

### Analysis of tumor intravasation and metastasis *in vivo*

MTLn3 cells that stably expressed EGFP and featured the CRISPR/Cas9-mediated TC10 deletion were injected into the mammary glands of female SCID mice (6–8-wk-old; Jackson ImmunoResearch Laboratories, Inc., West Grove, PA, USA ^84^). A total of 1.0 × 10^6^ cells were trypsinized and resuspended in 100 μl PBS for injection into each mouse (CRISPR/Cas9 non-targeting control, n = 20 mice; TC10-knockout, n = 20 mice). Mice were sacrificed 3–4 wk after injection when the primary tumor reached 1 cm in diameter. Lung metastases were confirmed and counted at necropsy using a fluorescent microscope to image EGFP fluorescence in freshly excised and isolated lungs mounted on a microscope coverslip. Twenty randomly selected fields of view at 10× magnification per mouse lung (10 fields of view per lung lobe) were analyzed to determine the ratio between total EGFP fluorescence and background fluorescence. To quantify the circulating tumor cell counts, 1 ml of mouse blood, obtained through cardiopuncture at the time of euthanasia, was lysed in red blood cell lysis buffer (04-4300-54; Thermo Fisher Scientific/eBioscience, San Diego, CA, USA), according to the manufacturer’s protocols. The remaining cells were plated into MTLn3 growth media and cultured for one additional week. The numbers of EGFP-positive MTLn3 cells were quantified in 1/4 of the area of a 10-cm tissue culture dish for each animal. All animal experiments were performed in accordance with a protocol approved by the Office of the Institutional Animal Care and Use Committee of the Albert Einstein College of Medicine (protocol 20170507). For data analysis, mice with primary tumors that showed indications of ulceration or intraperitoneal growths were omitted from the final tally.

### Statistical analysis

All statistical significance based on p-values were calculated using a Student’s *t*-test, unless stated otherwise in the figure legend. No vertebrate animals were involved. No statistical methods were used to pre-determine the sample size. No randomizations were used. The investigators were not blinded to allocation during experiments and outcome assessment. Statistical tests used are stated on every figure legend with p-values as appropriate. Data distribution should meet the normal distribution requirements. No estimate of variation. No pre-established criteria were used to determine data inclusion or exclusion.

## Data availability

The data that support the findings of this study are available from the corresponding author on request.

## Code availability

All Matlab codes and Metamorph scripts used were previously published elsewhere ^45,46,80^, but are also available from the corresponding author on request.

