## Supplemental Figures for "TC10 regulates breast cancer invasion and metastasis by controlling membrane type-1 matrix metalloproteinase at invadopodia"

### Slide 1
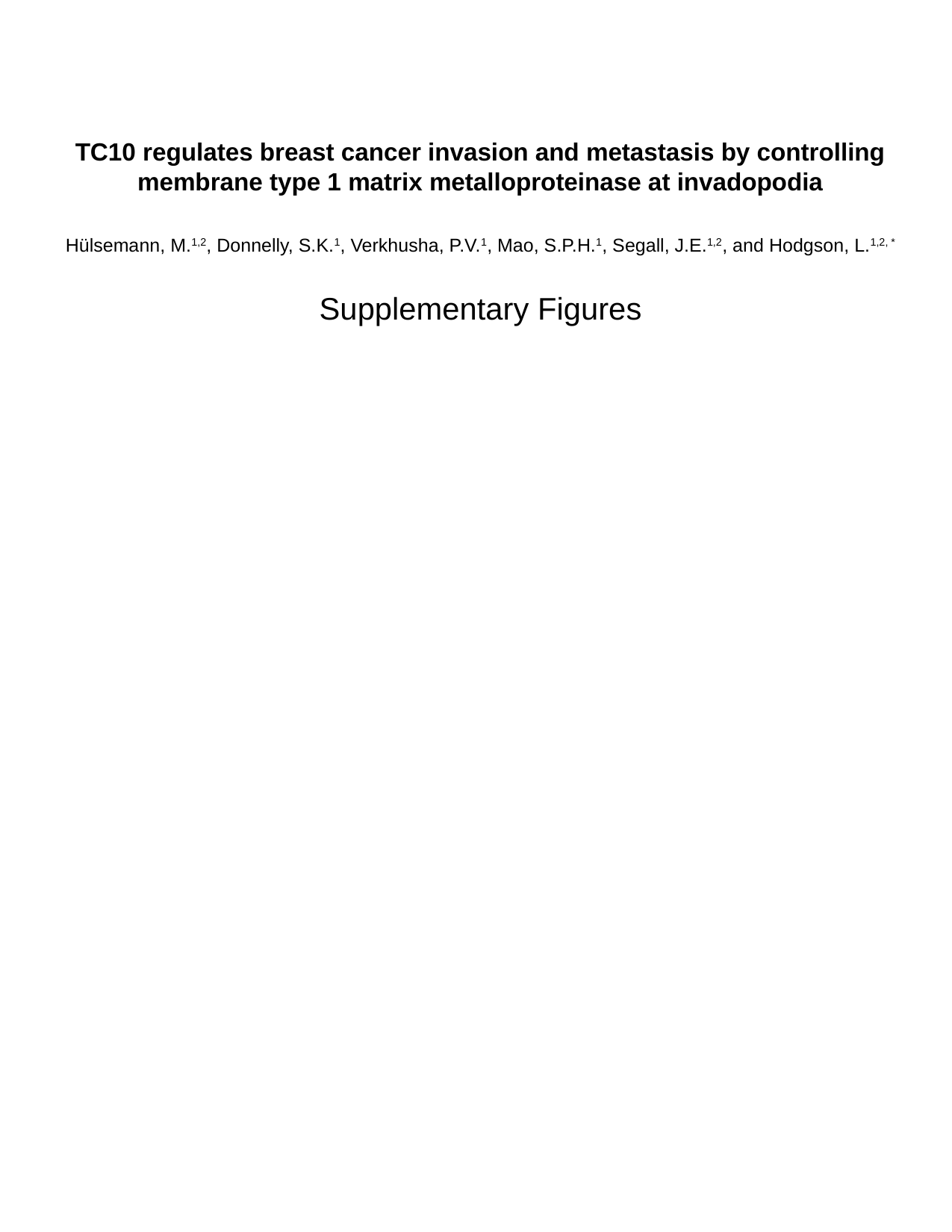

TC10 regulates breast cancer invasion and metastasis by controlling membrane type 1 matrix metalloproteinase at invadopodia
Hülsemann, M.^1,2^, Donnelly, S.K.^1^, Verkhusha, P.V.^1^, Mao, S.P.H.^1^, Segall, J.E.^1,2^, and Hodgson, L.^1,2, *^
Supplementary Figures

### Slide 2
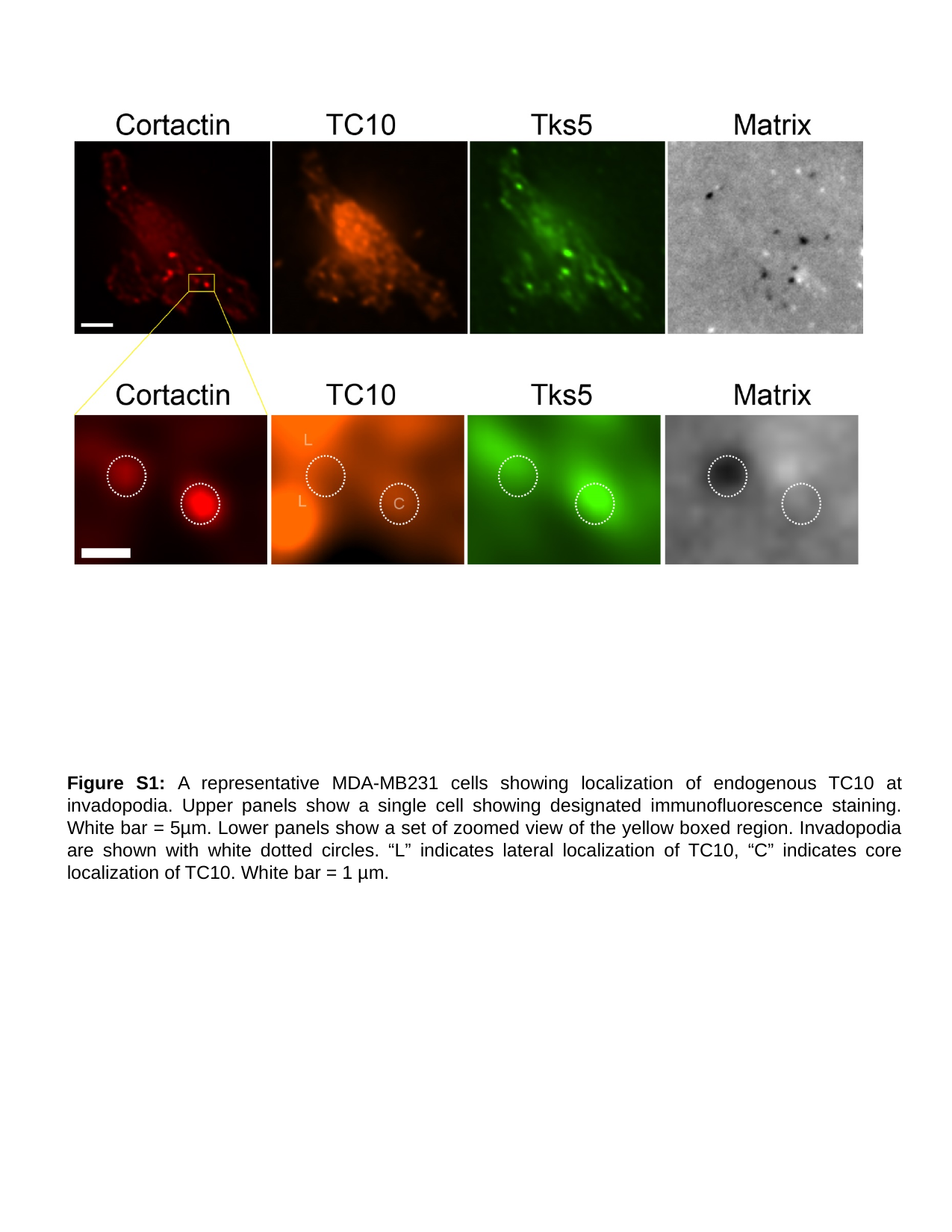

Figure S1: A representative MDA-MB231 cells showing localization of endogenous TC10 at invadopodia. Upper panels show a single cell showing designated immunofluorescence staining. White bar = 5µm. Lower panels show a set of zoomed view of the yellow boxed region. Invadopodia are shown with white dotted circles. “L” indicates lateral localization of TC10, “C” indicates core localization of TC10. White bar = 1 µm.

### Slide 3
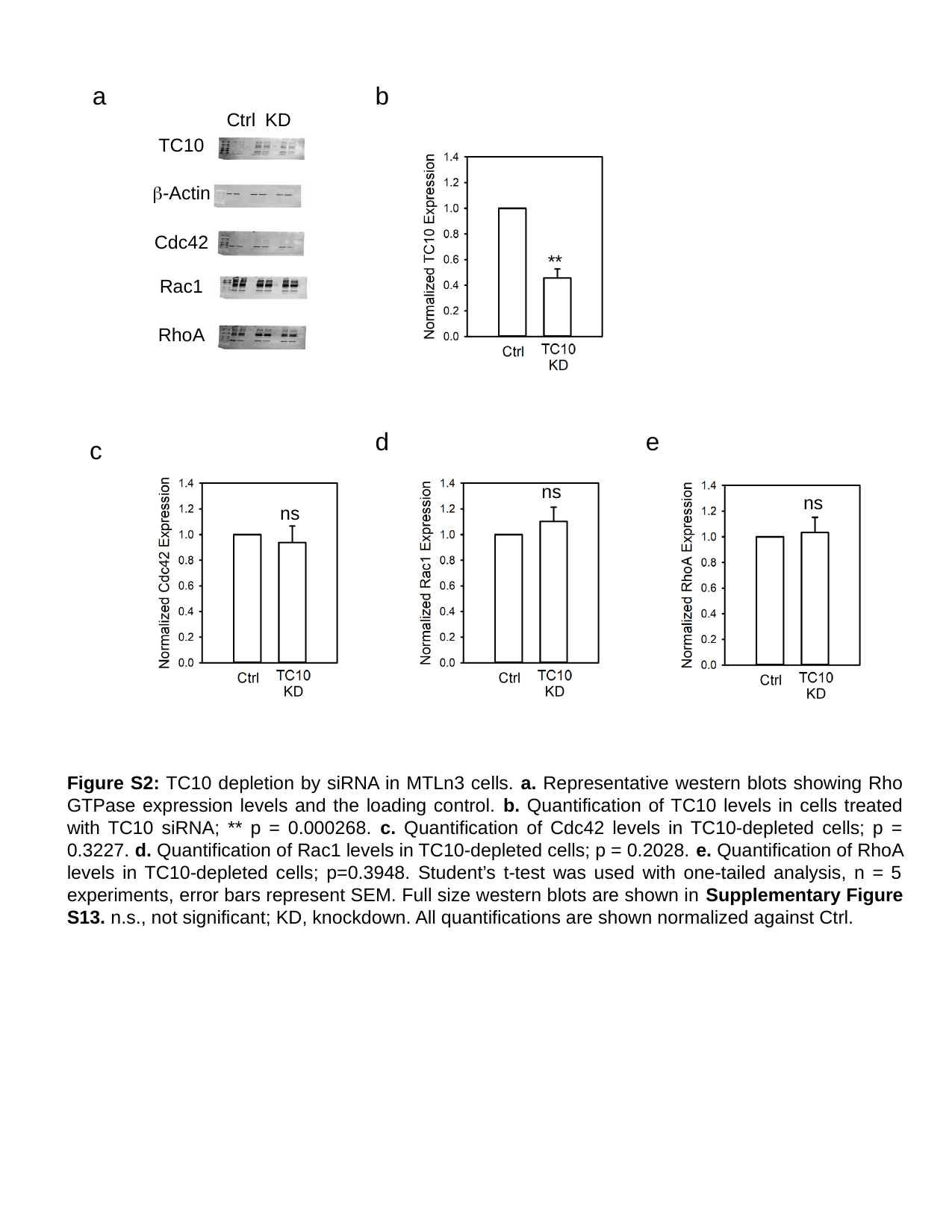

a
b
Ctrl
KD
TC10
β-Actin
Cdc42
Rac1
RhoA
**
d
e
c
ns
ns
ns
Figure S2: TC10 depletion by siRNA in MTLn3 cells. a. Representative western blots showing Rho GTPase expression levels and the loading control. b. Quantification of TC10 levels in cells treated with TC10 siRNA; ** p = 0.000268. c. Quantification of Cdc42 levels in TC10-depleted cells; p = 0.3227. d. Quantification of Rac1 levels in TC10-depleted cells; p = 0.2028. e. Quantification of RhoA levels in TC10-depleted cells; p=0.3948. Student’s t-test was used with one-tailed analysis, n = 5 experiments, error bars represent SEM. Full size western blots are shown in Supplementary Figure S13. n.s., not significant; KD, knockdown. All quantifications are shown normalized against Ctrl.

### Slide 4
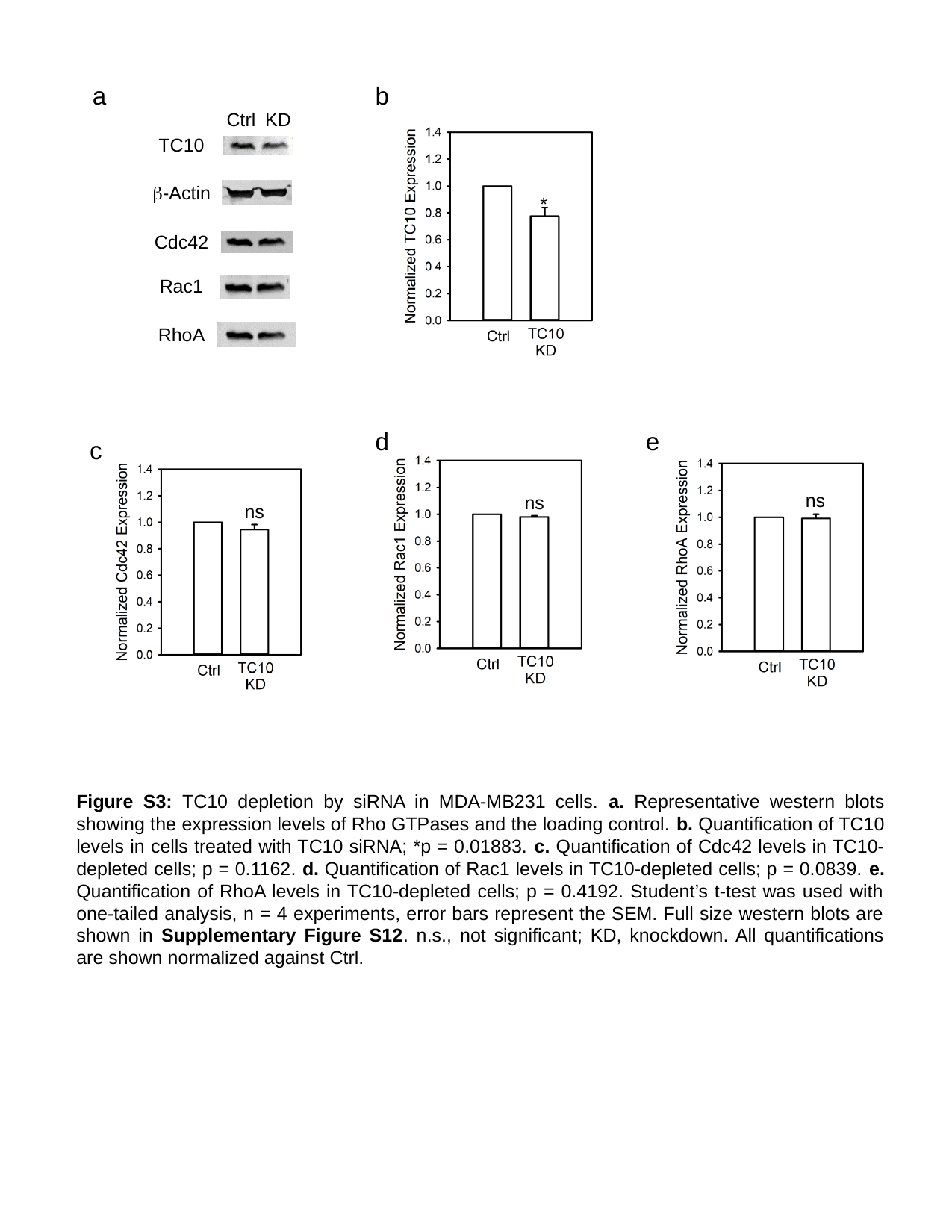

a
b
Ctrl
KD
*
TC10
β-Actin
Cdc42
Rac1
RhoA
d
e
c
ns
ns
ns
Figure S3: TC10 depletion by siRNA in MDA-MB231 cells. a. Representative western blots showing the expression levels of Rho GTPases and the loading control. b. Quantification of TC10 levels in cells treated with TC10 siRNA; *p = 0.01883. c. Quantification of Cdc42 levels in TC10-depleted cells; p = 0.1162. d. Quantification of Rac1 levels in TC10-depleted cells; p = 0.0839. e. Quantification of RhoA levels in TC10-depleted cells; p = 0.4192. Student’s t-test was used with one-tailed analysis, n = 4 experiments, error bars represent the SEM. Full size western blots are shown in Supplementary Figure S12. n.s., not significant; KD, knockdown. All quantifications are shown normalized against Ctrl.

### Slide 5
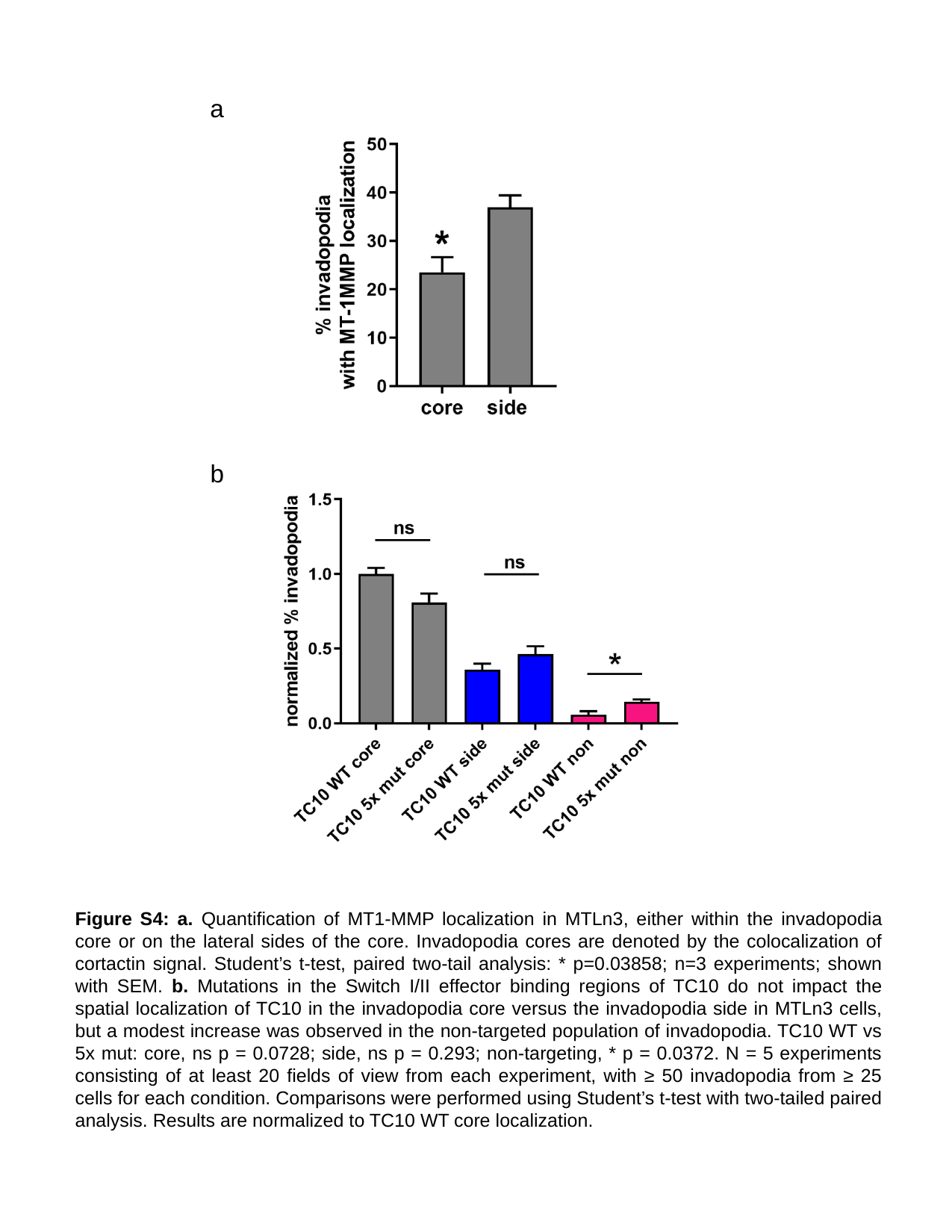

a
b
Figure S4: a. Quantification of MT1-MMP localization in MTLn3, either within the invadopodia core or on the lateral sides of the core. Invadopodia cores are denoted by the colocalization of cortactin signal. Student’s t-test, paired two-tail analysis: * p=0.03858; n=3 experiments; shown with SEM. b. Mutations in the Switch I/II effector binding regions of TC10 do not impact the spatial localization of TC10 in the invadopodia core versus the invadopodia side in MTLn3 cells, but a modest increase was observed in the non-targeted population of invadopodia. TC10 WT vs 5x mut: core, ns p = 0.0728; side, ns p = 0.293; non-targeting, * p = 0.0372. N = 5 experiments consisting of at least 20 fields of view from each experiment, with ≥ 50 invadopodia from ≥ 25 cells for each condition. Comparisons were performed using Student’s t-test with two-tailed paired analysis. Results are normalized to TC10 WT core localization.

### Slide 6
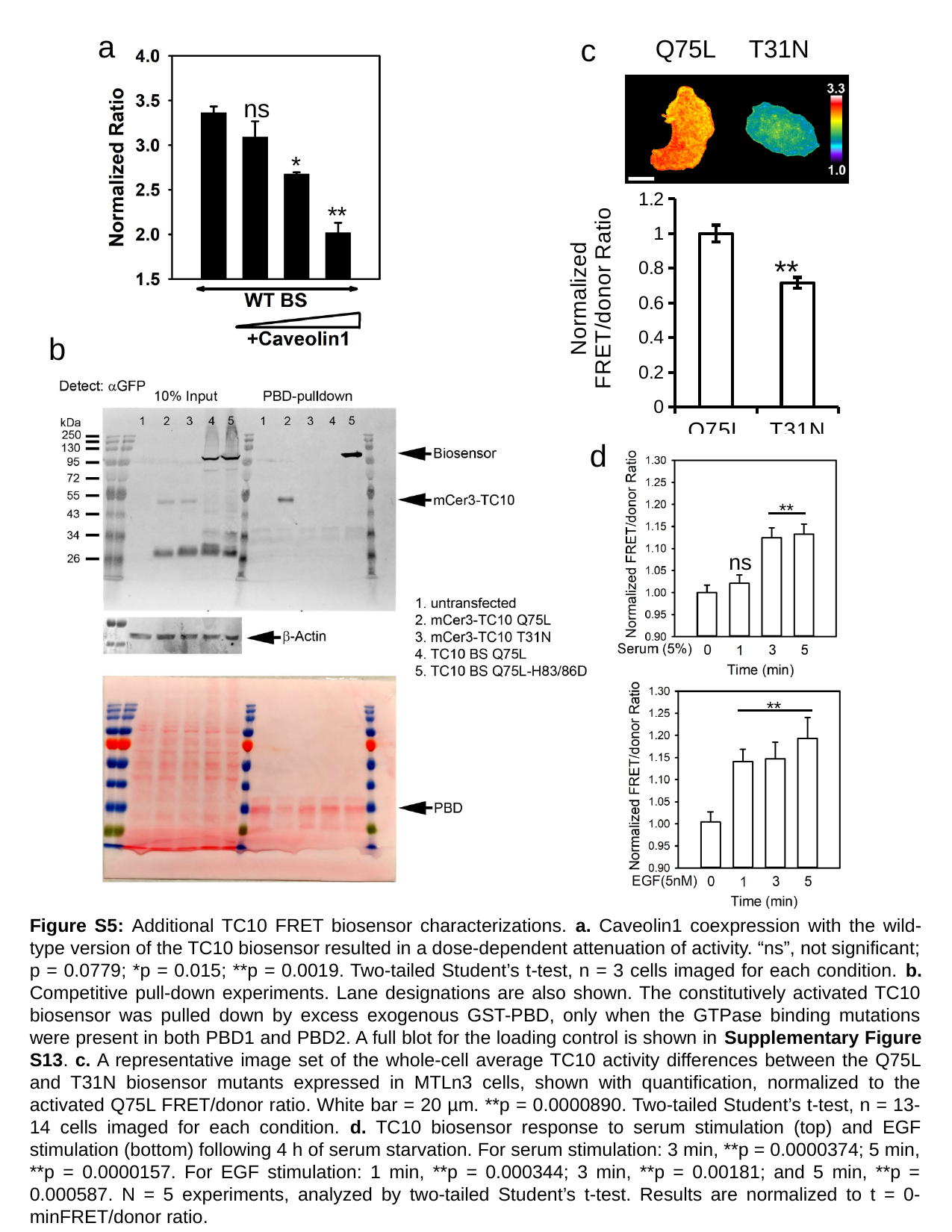

a
c
Q75L
T31N
ns
*
#### Chart
| Category | |
|---|---|
| Q75L | 0.9999999999999999 |
| T31N | 0.716797169464709 |**
**
b
d
**
ns
**
Figure S5: Additional TC10 FRET biosensor characterizations. a. Caveolin1 coexpression with the wild-type version of the TC10 biosensor resulted in a dose-dependent attenuation of activity. “ns”, not significant; p = 0.0779; *p = 0.015; **p = 0.0019. Two-tailed Student’s t-test, n = 3 cells imaged for each condition. b. Competitive pull-down experiments. Lane designations are also shown. The constitutively activated TC10 biosensor was pulled down by excess exogenous GST-PBD, only when the GTPase binding mutations were present in both PBD1 and PBD2. A full blot for the loading control is shown in Supplementary Figure S13. c. A representative image set of the whole-cell average TC10 activity differences between the Q75L and T31N biosensor mutants expressed in MTLn3 cells, shown with quantification, normalized to the activated Q75L FRET/donor ratio. White bar = 20 µm. **p = 0.0000890. Two-tailed Student’s t-test, n = 13-14 cells imaged for each condition. d. TC10 biosensor response to serum stimulation (top) and EGF stimulation (bottom) following 4 h of serum starvation. For serum stimulation: 3 min, **p = 0.0000374; 5 min, **p = 0.0000157. For EGF stimulation: 1 min, **p = 0.000344; 3 min, **p = 0.00181; and 5 min, **p = 0.000587. N = 5 experiments, analyzed by two-tailed Student’s t-test. Results are normalized to t = 0-minFRET/donor ratio.

### Slide 7
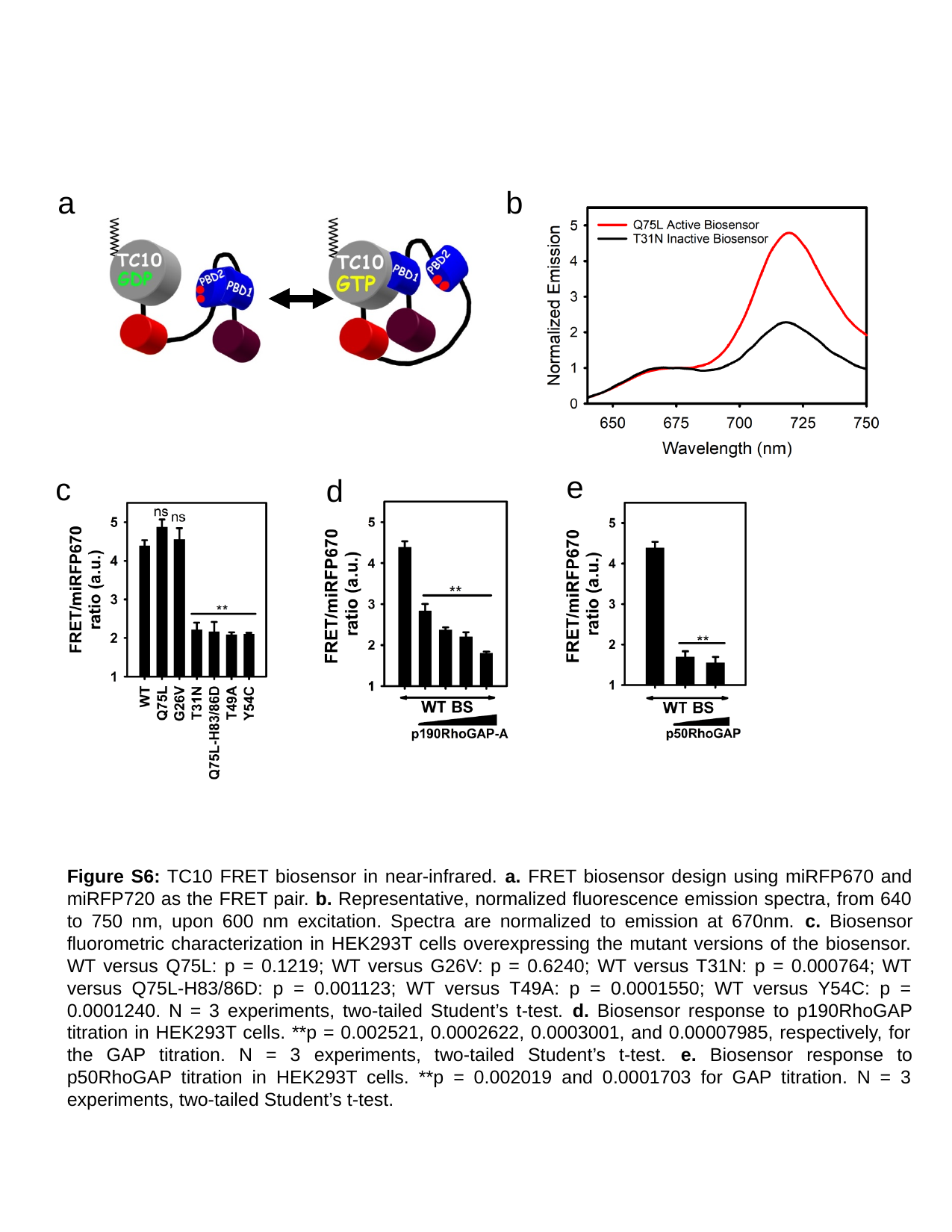

a
b
e
c
d
Figure S6: TC10 FRET biosensor in near-infrared. a. FRET biosensor design using miRFP670 and miRFP720 as the FRET pair. b. Representative, normalized fluorescence emission spectra, from 640 to 750 nm, upon 600 nm excitation. Spectra are normalized to emission at 670nm. c. Biosensor fluorometric characterization in HEK293T cells overexpressing the mutant versions of the biosensor. WT versus Q75L: p = 0.1219; WT versus G26V: p = 0.6240; WT versus T31N: p = 0.000764; WT versus Q75L-H83/86D: p = 0.001123; WT versus T49A: p = 0.0001550; WT versus Y54C: p = 0.0001240. N = 3 experiments, two-tailed Student’s t-test. d. Biosensor response to p190RhoGAP titration in HEK293T cells. **p = 0.002521, 0.0002622, 0.0003001, and 0.00007985, respectively, for the GAP titration. N = 3 experiments, two-tailed Student’s t-test. e. Biosensor response to p50RhoGAP titration in HEK293T cells. **p = 0.002019 and 0.0001703 for GAP titration. N = 3 experiments, two-tailed Student’s t-test.

### Slide 8
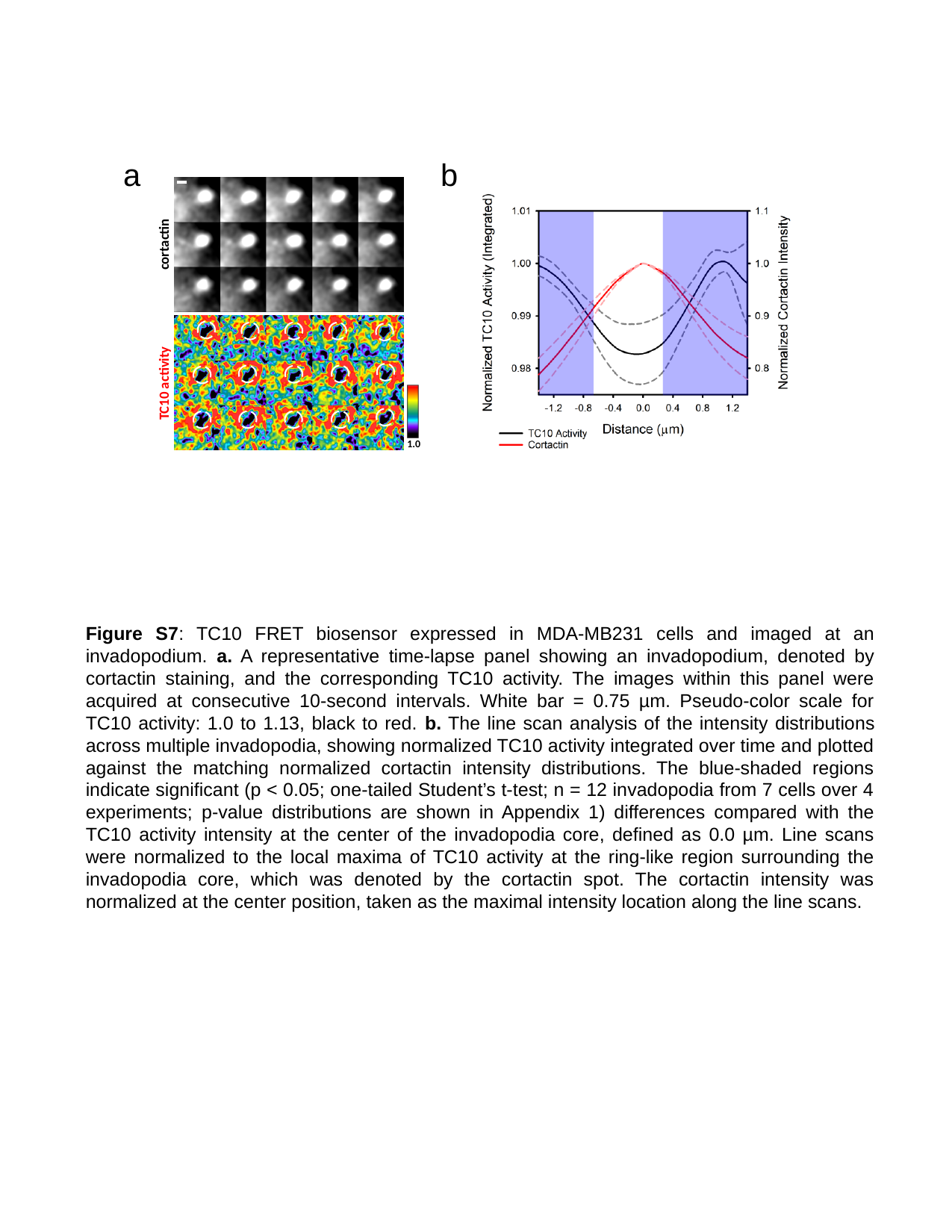

a
b
cortactin
TC10 activity
1.0
Figure S7: TC10 FRET biosensor expressed in MDA-MB231 cells and imaged at an invadopodium. a. A representative time-lapse panel showing an invadopodium, denoted by cortactin staining, and the corresponding TC10 activity. The images within this panel were acquired at consecutive 10-second intervals. White bar = 0.75 µm. Pseudo-color scale for TC10 activity: 1.0 to 1.13, black to red. b. The line scan analysis of the intensity distributions across multiple invadopodia, showing normalized TC10 activity integrated over time and plotted against the matching normalized cortactin intensity distributions. The blue-shaded regions indicate significant (p < 0.05; one-tailed Student’s t-test; n = 12 invadopodia from 7 cells over 4 experiments; p-value distributions are shown in Appendix 1) differences compared with the TC10 activity intensity at the center of the invadopodia core, defined as 0.0 µm. Line scans were normalized to the local maxima of TC10 activity at the ring-like region surrounding the invadopodia core, which was denoted by the cortactin spot. The cortactin intensity was normalized at the center position, taken as the maximal intensity location along the line scans.

### Slide 9
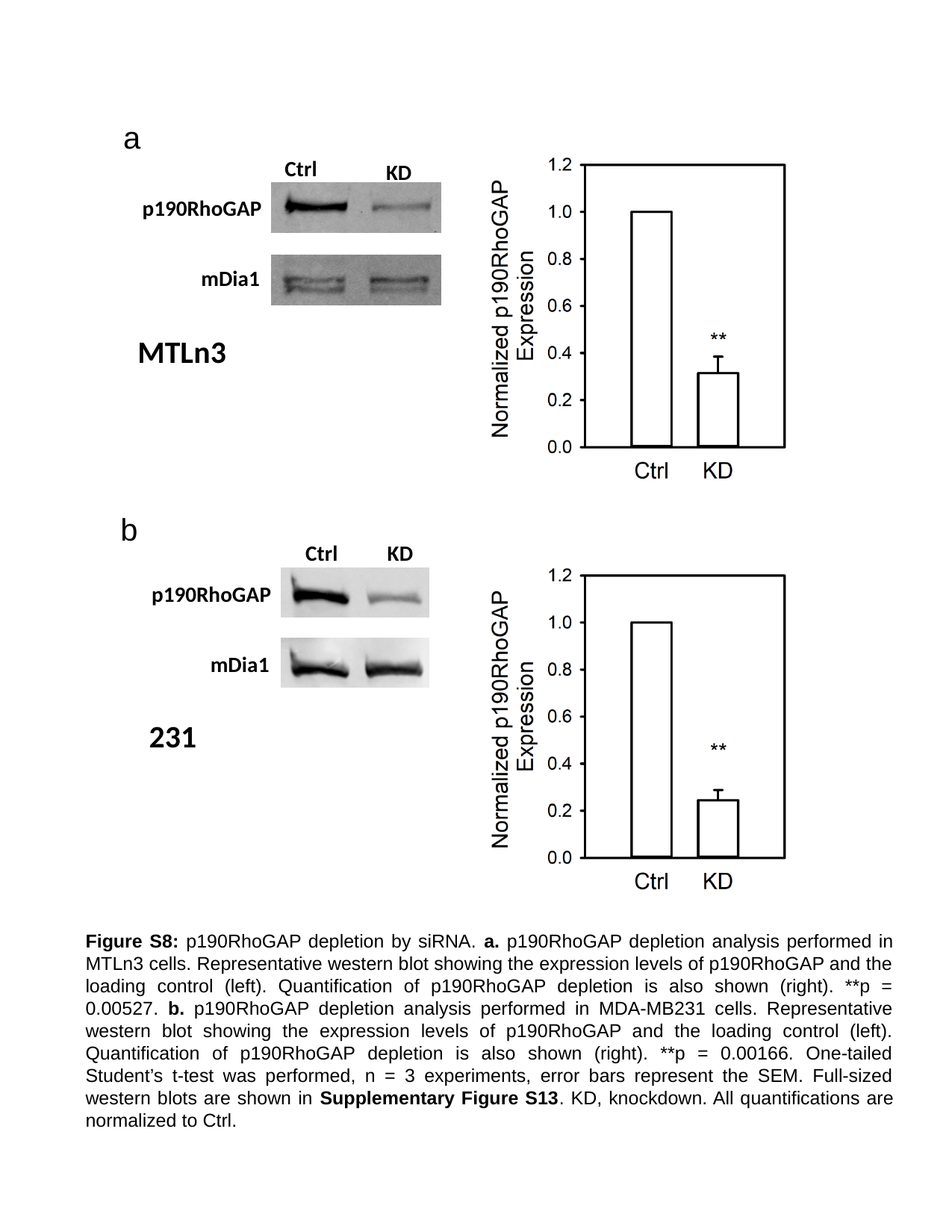

a
Ctrl
KD
p190RhoGAP
mDia1
MTLn3
b
Ctrl
KD
p190RhoGAP
mDia1
231
Figure S8: p190RhoGAP depletion by siRNA. a. p190RhoGAP depletion analysis performed in MTLn3 cells. Representative western blot showing the expression levels of p190RhoGAP and the loading control (left). Quantification of p190RhoGAP depletion is also shown (right). **p = 0.00527. b. p190RhoGAP depletion analysis performed in MDA-MB231 cells. Representative western blot showing the expression levels of p190RhoGAP and the loading control (left). Quantification of p190RhoGAP depletion is also shown (right). **p = 0.00166. One-tailed Student’s t-test was performed, n = 3 experiments, error bars represent the SEM. Full-sized western blots are shown in Supplementary Figure S13. KD, knockdown. All quantifications are normalized to Ctrl.

### Slide 10
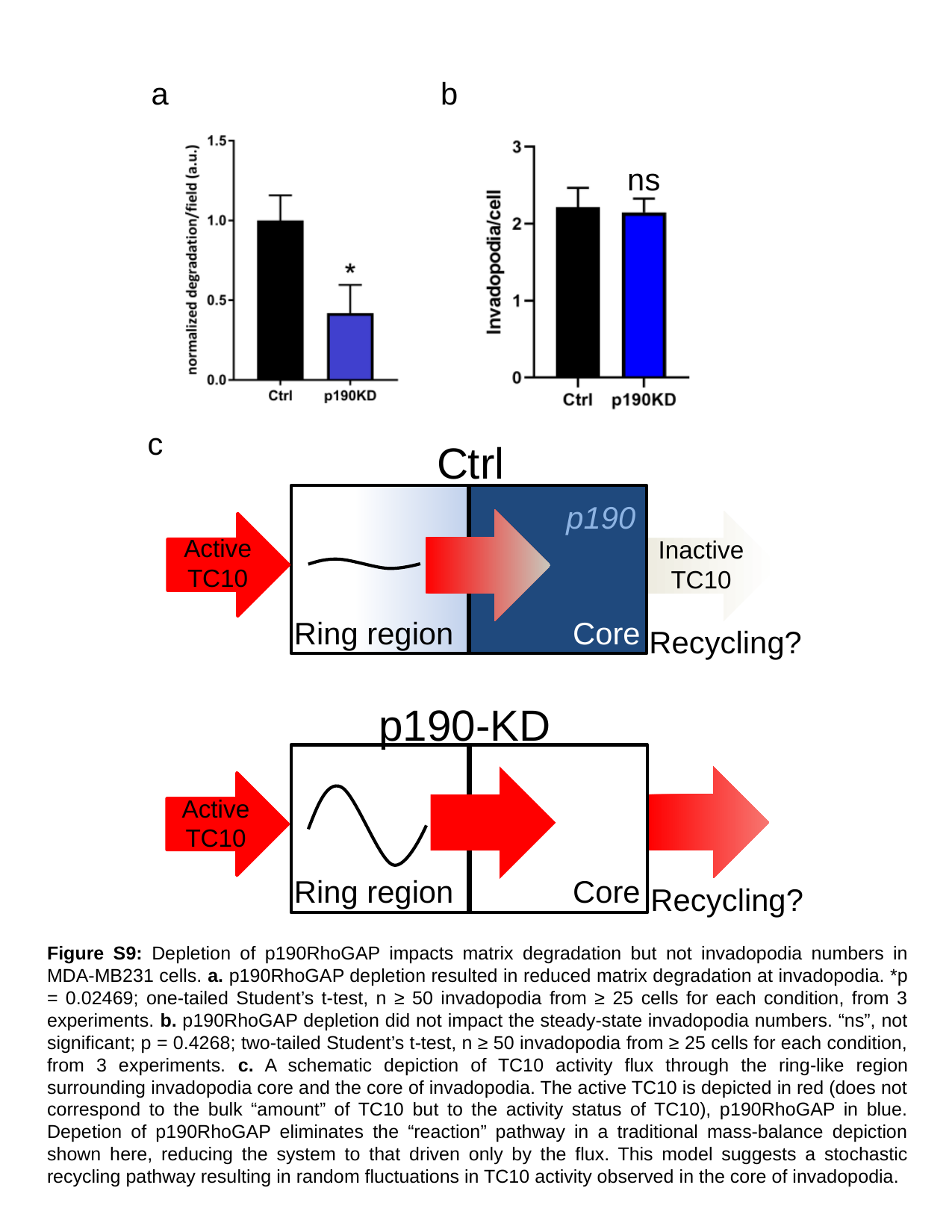

a
b
ns
c
Ctrl
p190
Active TC10
Inactive TC10
Ring region
Core
Recycling?
p190-KD
Active TC10
Ring region
Core
Recycling?
Figure S9: Depletion of p190RhoGAP impacts matrix degradation but not invadopodia numbers in MDA-MB231 cells. a. p190RhoGAP depletion resulted in reduced matrix degradation at invadopodia. *p = 0.02469; one-tailed Student’s t-test, n ≥ 50 invadopodia from ≥ 25 cells for each condition, from 3 experiments. b. p190RhoGAP depletion did not impact the steady-state invadopodia numbers. “ns”, not significant; p = 0.4268; two-tailed Student’s t-test, n ≥ 50 invadopodia from ≥ 25 cells for each condition, from 3 experiments. c. A schematic depiction of TC10 activity flux through the ring-like region surrounding invadopodia core and the core of invadopodia. The active TC10 is depicted in red (does not correspond to the bulk “amount” of TC10 but to the activity status of TC10), p190RhoGAP in blue. Depetion of p190RhoGAP eliminates the “reaction” pathway in a traditional mass-balance depiction shown here, reducing the system to that driven only by the flux. This model suggests a stochastic recycling pathway resulting in random fluctuations in TC10 activity observed in the core of invadopodia.

### Slide 11
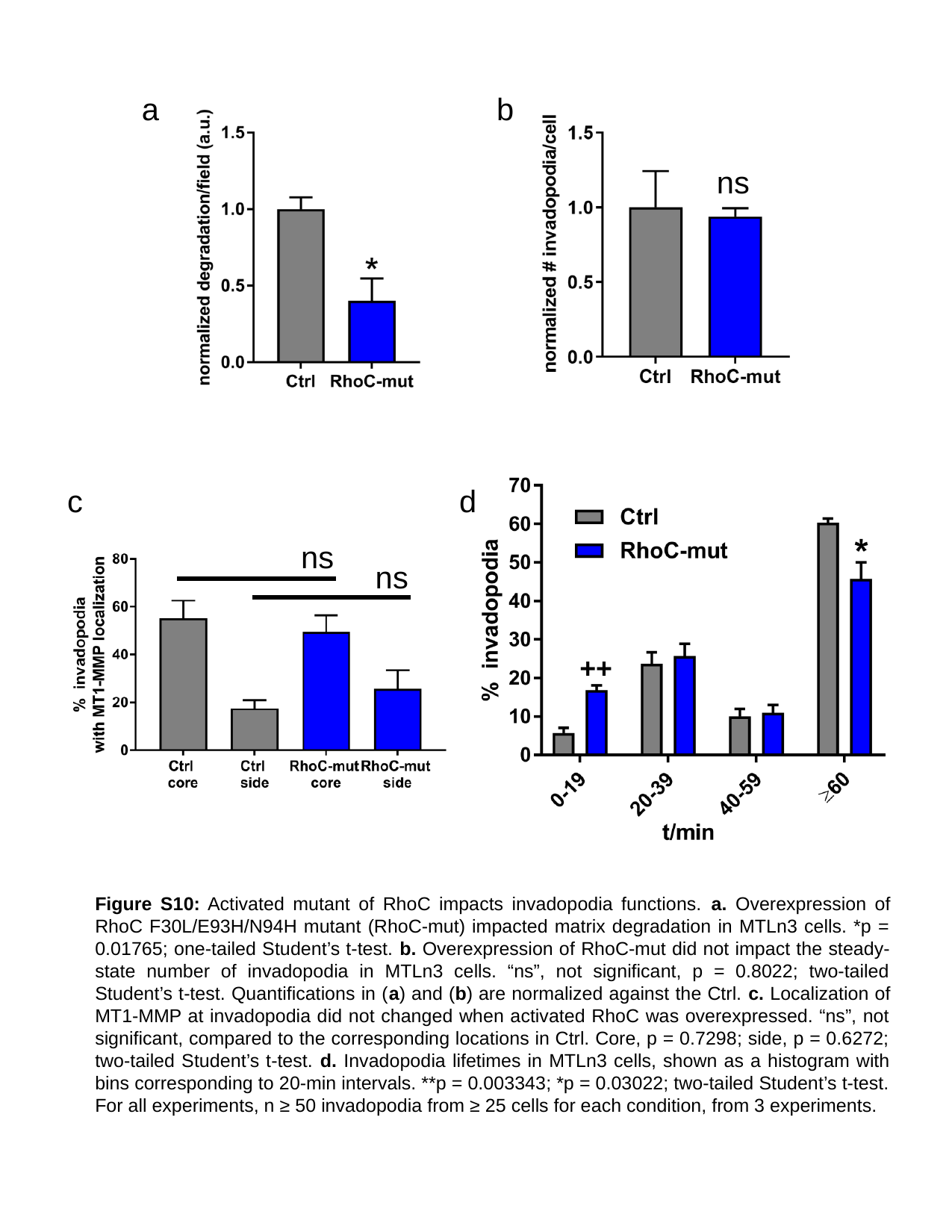

a
b
ns
++
c
d
ns
ns
Figure S10: Activated mutant of RhoC impacts invadopodia functions. a. Overexpression of RhoC F30L/E93H/N94H mutant (RhoC-mut) impacted matrix degradation in MTLn3 cells. *p = 0.01765; one-tailed Student’s t-test. b. Overexpression of RhoC-mut did not impact the steady-state number of invadopodia in MTLn3 cells. “ns”, not significant, p = 0.8022; two-tailed Student’s t-test. Quantifications in (a) and (b) are normalized against the Ctrl. c. Localization of MT1-MMP at invadopodia did not changed when activated RhoC was overexpressed. “ns”, not significant, compared to the corresponding locations in Ctrl. Core, p = 0.7298; side, p = 0.6272; two-tailed Student’s t-test. d. Invadopodia lifetimes in MTLn3 cells, shown as a histogram with bins corresponding to 20-min intervals. **p = 0.003343; *p = 0.03022; two-tailed Student’s t-test. For all experiments, n ≥ 50 invadopodia from ≥ 25 cells for each condition, from 3 experiments.

### Slide 12
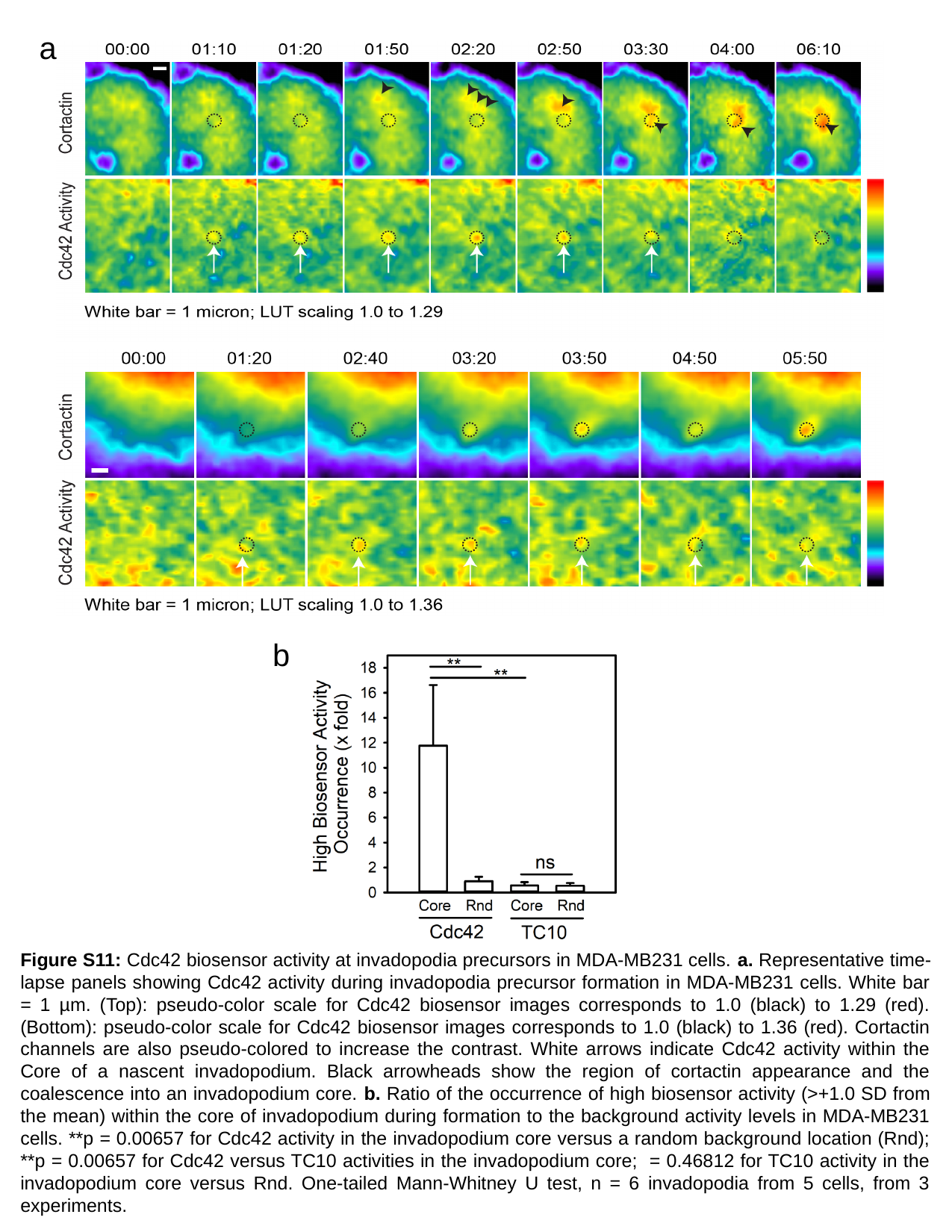

a
b
Figure S11: Cdc42 biosensor activity at invadopodia precursors in MDA-MB231 cells. a. Representative time-lapse panels showing Cdc42 activity during invadopodia precursor formation in MDA-MB231 cells. White bar = 1 µm. (Top): pseudo-color scale for Cdc42 biosensor images corresponds to 1.0 (black) to 1.29 (red). (Bottom): pseudo-color scale for Cdc42 biosensor images corresponds to 1.0 (black) to 1.36 (red). Cortactin channels are also pseudo-colored to increase the contrast. White arrows indicate Cdc42 activity within the Core of a nascent invadopodium. Black arrowheads show the region of cortactin appearance and the coalescence into an invadopodium core. b. Ratio of the occurrence of high biosensor activity (>+1.0 SD from the mean) within the core of invadopodium during formation to the background activity levels in MDA-MB231 cells. **p = 0.00657 for Cdc42 activity in the invadopodium core versus a random background location (Rnd); **p = 0.00657 for Cdc42 versus TC10 activities in the invadopodium core; = 0.46812 for TC10 activity in the invadopodium core versus Rnd. One-tailed Mann-Whitney U test, n = 6 invadopodia from 5 cells, from 3 experiments.

### Slide 13
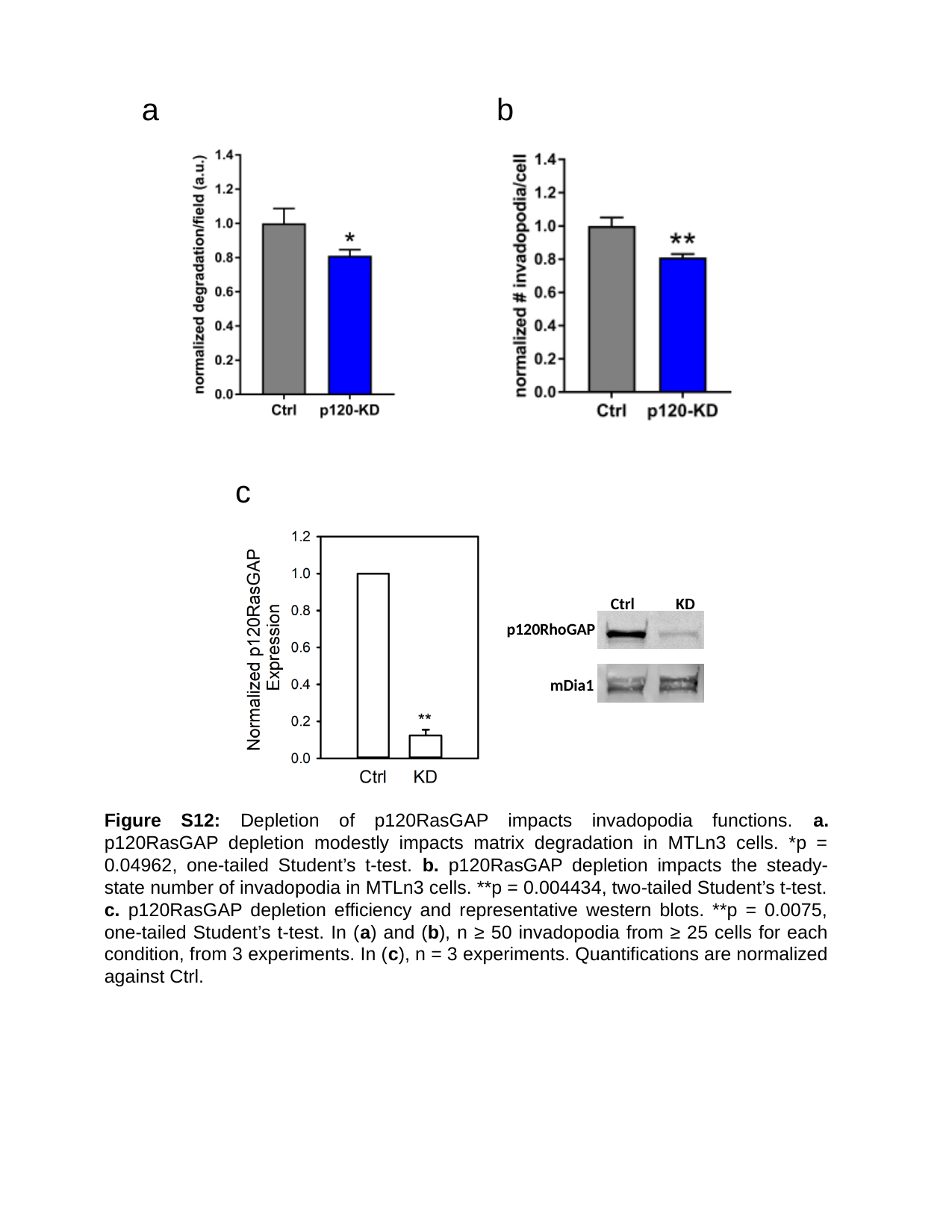

a
b
c
Ctrl
KD
p120RhoGAP
mDia1
Figure S12: Depletion of p120RasGAP impacts invadopodia functions. a. p120RasGAP depletion modestly impacts matrix degradation in MTLn3 cells. *p = 0.04962, one-tailed Student’s t-test. b. p120RasGAP depletion impacts the steady-state number of invadopodia in MTLn3 cells. **p = 0.004434, two-tailed Student’s t-test. c. p120RasGAP depletion efficiency and representative western blots. **p = 0.0075, one-tailed Student’s t-test. In (a) and (b), n ≥ 50 invadopodia from ≥ 25 cells for each condition, from 3 experiments. In (c), n = 3 experiments. Quantifications are normalized against Ctrl.

### Slide 14
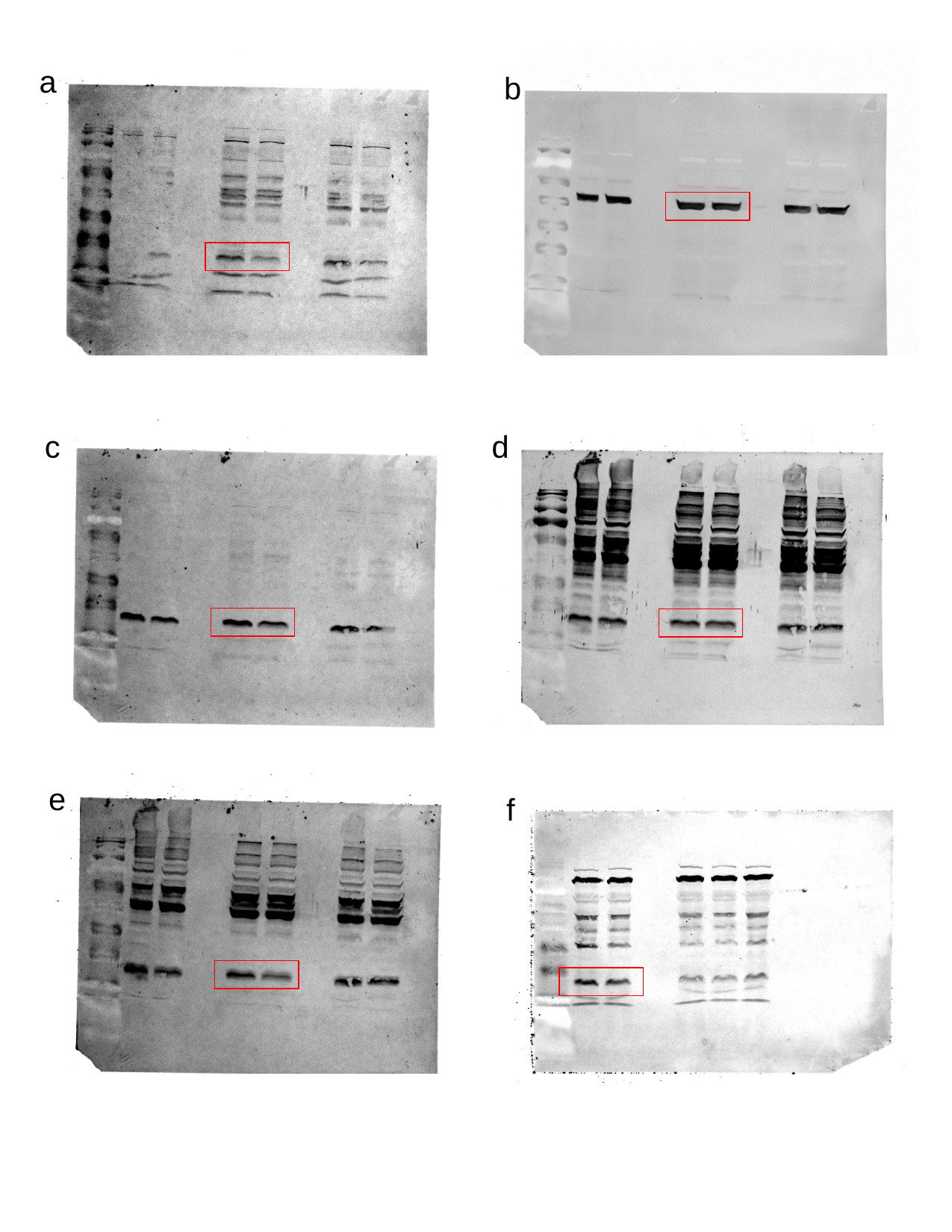

a
b
d
c
e
f

### Slide 15
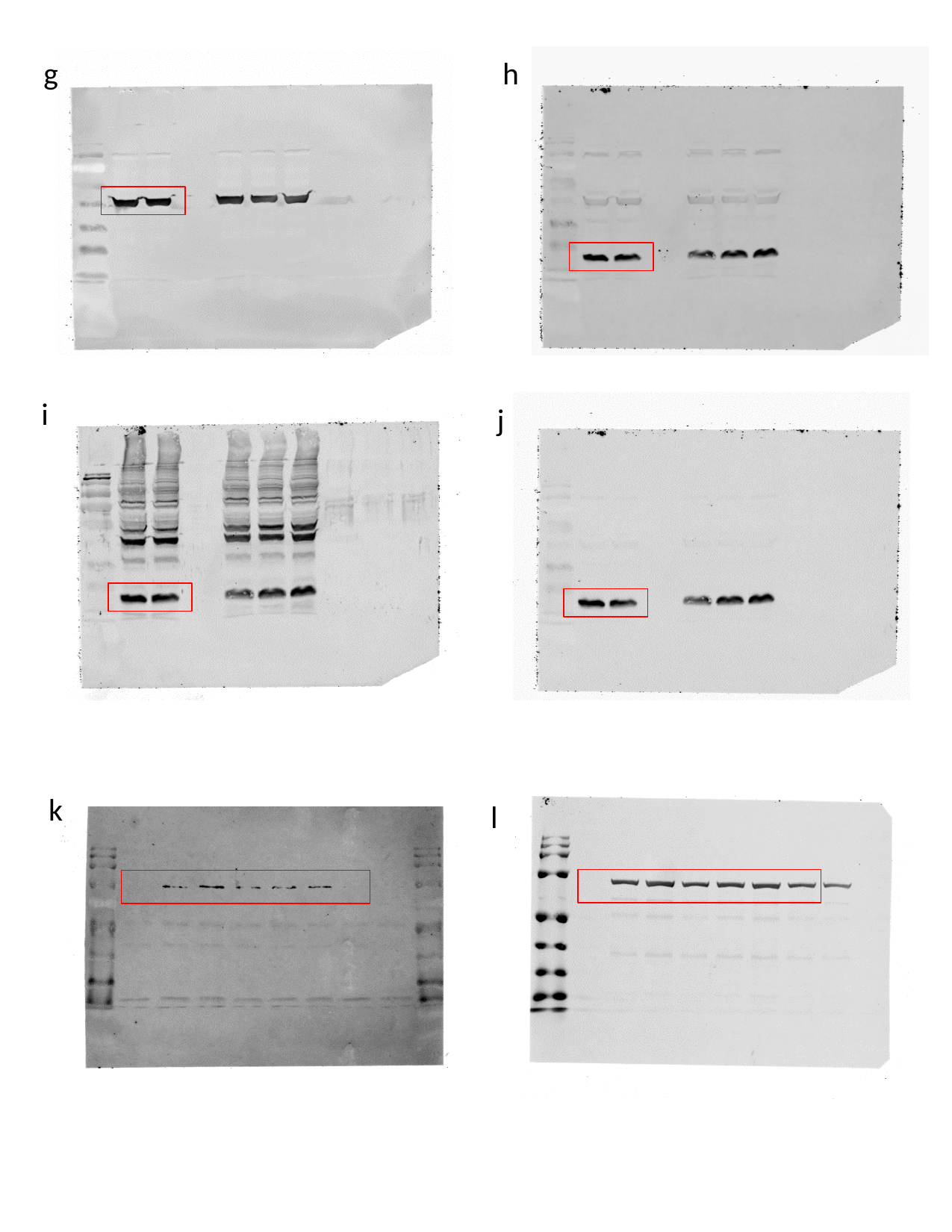

g
h
i
j
k
l

### Slide 16
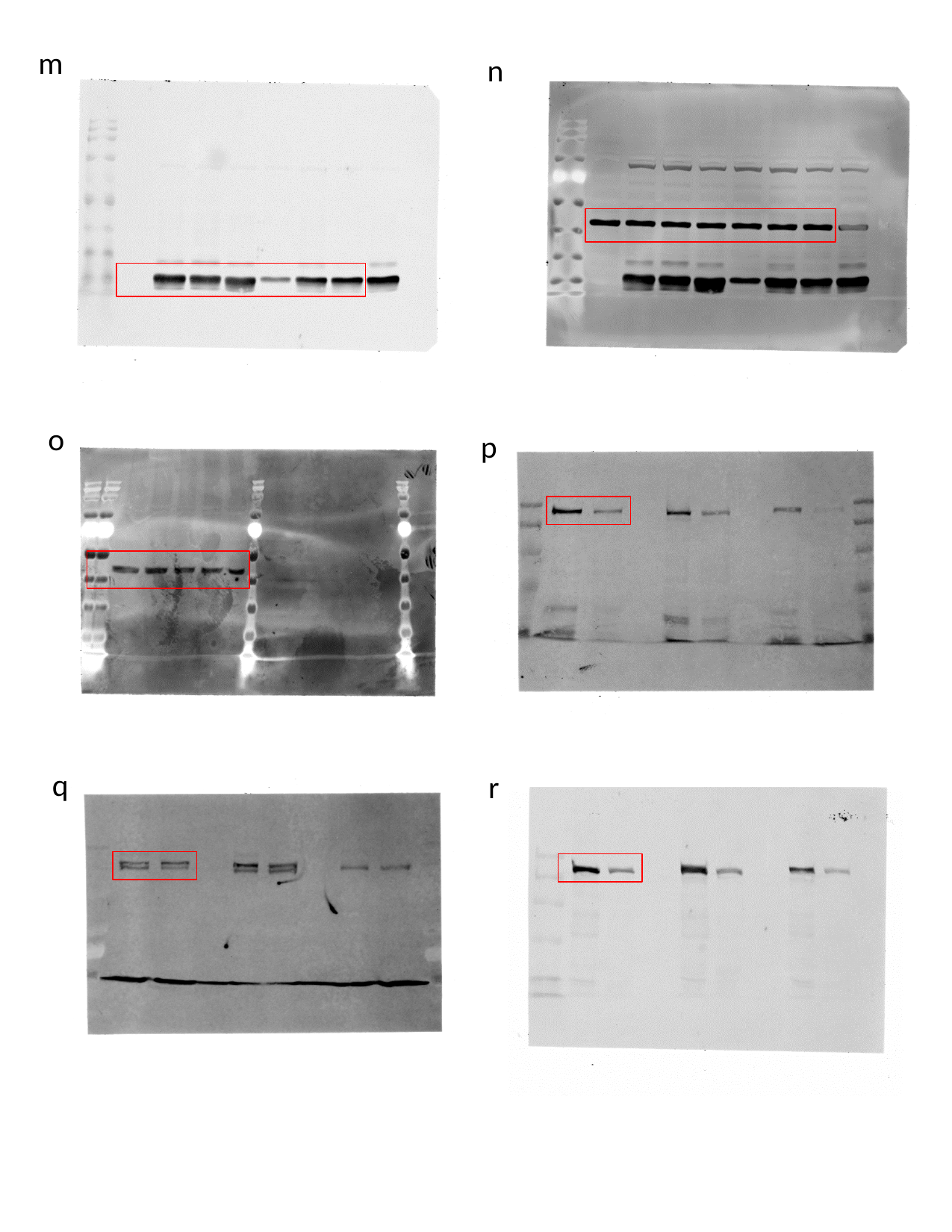

m
n
o
p
q
r

### Slide 17
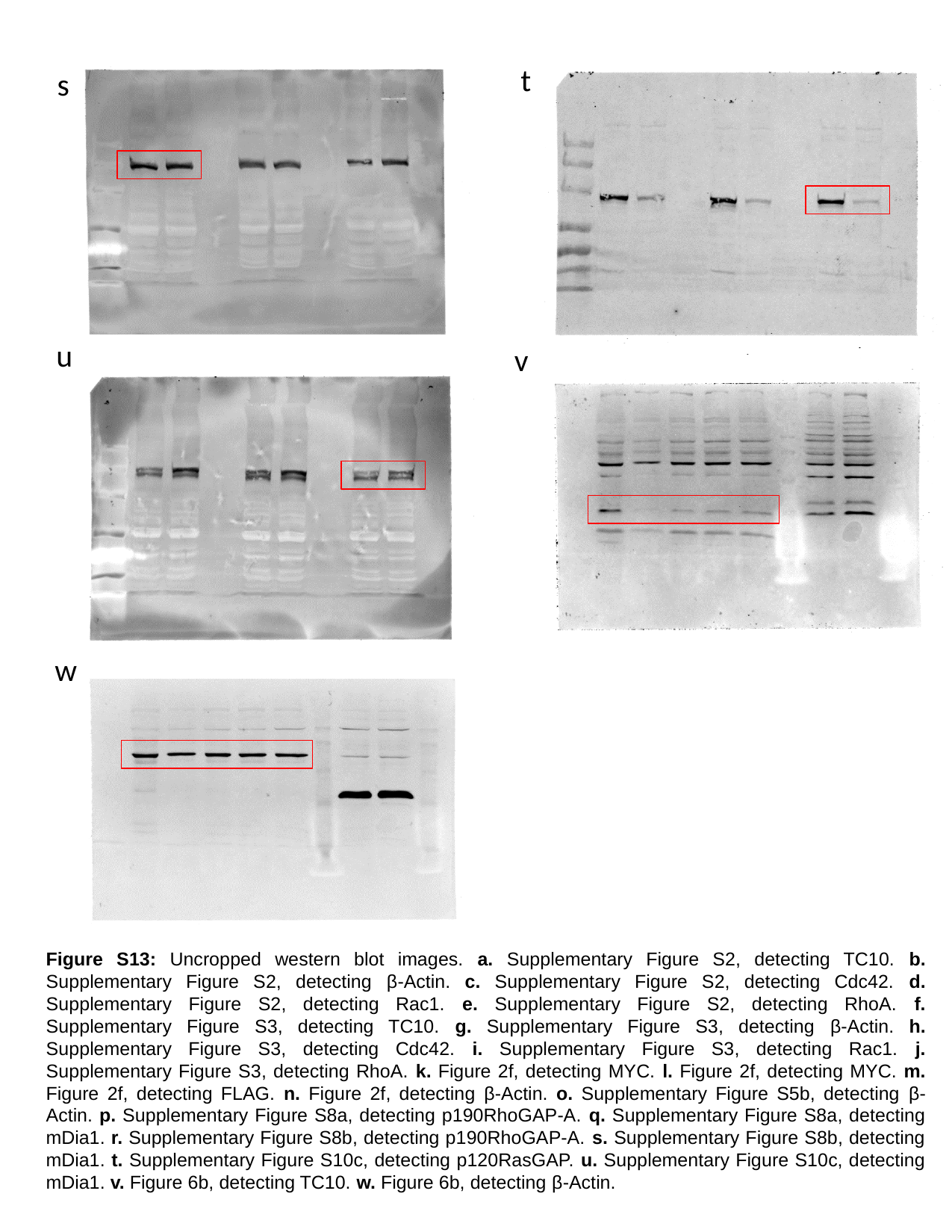

t
s
u
v
w
Figure S13: Uncropped western blot images. a. Supplementary Figure S2, detecting TC10. b. Supplementary Figure S2, detecting β-Actin. c. Supplementary Figure S2, detecting Cdc42. d. Supplementary Figure S2, detecting Rac1. e. Supplementary Figure S2, detecting RhoA. f. Supplementary Figure S3, detecting TC10. g. Supplementary Figure S3, detecting β-Actin. h. Supplementary Figure S3, detecting Cdc42. i. Supplementary Figure S3, detecting Rac1. j. Supplementary Figure S3, detecting RhoA. k. Figure 2f, detecting MYC. l. Figure 2f, detecting MYC. m. Figure 2f, detecting FLAG. n. Figure 2f, detecting β-Actin. o. Supplementary Figure S5b, detecting β-Actin. p. Supplementary Figure S8a, detecting p190RhoGAP-A. q. Supplementary Figure S8a, detecting mDia1. r. Supplementary Figure S8b, detecting p190RhoGAP-A. s. Supplementary Figure S8b, detecting mDia1. t. Supplementary Figure S10c, detecting p120RasGAP. u. Supplementary Figure S10c, detecting mDia1. v. Figure 6b, detecting TC10. w. Figure 6b, detecting β-Actin.
