## Appendix Data for "TC10 regulates breast cancer invasion and metastasis by controlling membrane type-1 matrix metalloproteinase at invadopodia"

### Appendix 1:

P-value limits for Linescan analyses of invadopodia

**Figure 3g: MTLn3 invadopodia**

Red-highlighted cells show $p<0.05$ compared to the center of invadopodia

| X-position (micron) | p-value |
|---|---|
| -2.325 | 0.333324342 |
| -2.2875 | 0.208341653 |
| -2.25 | 0.296269208 |
| -2.2125 | 0.350796313 |
| -2.175 | 0.284425359 |
| -2.1375 | 0.101854135 |
| -2.1 | 0.039761836 |
| -2.0625 | 0.020733754 |
| -2.025 | 0.440452375 |
| -1.9875 | 0.475158242 |
| -1.95 | 0.472519412 |
| -1.9125 | 0.446133117 |
| -1.875 | 0.380729702 |
| -1.8375 | 0.316636021 |
| -1.8 | 0.254786302 |
| -1.7625 | 0.19000394 |
| -1.725 | 0.125409147 |
| -1.6875 | 0.417552361 |
| -1.65 | 0.276192979 |
| -1.6125 | 0.128916081 |
| -1.575 | 0.032170484 |
| -1.5375 | 0.009330844 |
| -1.5 | 0.007353831 |
| -1.4625 | 0.006621893 |
| -1.425 | 0.005689452 |
| -1.3875 | 0.005328448 |
| -1.35 | 0.005049157 |
| -1.3125 | 0.037907498 |
| -1.275 | 0.014168476 |
| -1.2375 | 0.006406293 |
| -1.2 | 0.004126149 |
| -1.1625 | 0.003475415 |
| -1.125 | 0.003334697 |

| | |
|---|---|
| -1.0875 | 0.003172104 |
| -1.05 | 0.002818788 |
| -1.0125 | 0.002572025 |
| -0.975 | 0.002255678 |
| -0.9375 | 0.002085959 |
| -0.9 | 0.002029547 |
| -0.8625 | 0.002112927 |
| -0.825 | 0.002409877 |
| -0.7875 | 0.003175228 |
| -0.75 | 0.004607168 |
| -0.7125 | 0.00695018 |
| -0.675 | 0.010806163 |
| -0.6375 | 0.016747789 |
| -0.6 | 0.026183937 |
| -0.5625 | 0.03979324 |
| -0.525 | 0.058705123 |
| -0.4875 | 0.079897876 |
| -0.45 | 0.108509542 |
| -0.4125 | 0.14119996 |
| -0.375 | 0.187114707 |
| -0.3375 | 0.236470397 |
| -0.3 | 0.288371794 |
| -0.2625 | 0.332177156 |
| -0.225 | 0.375859459 |
| -0.1875 | 0.407927339 |
| -0.15 | 0.416348955 |
| -0.1125 | 0.439032497 |
| -0.075 | 0.499316663 |
| -0.0375 | 0.456031005 |
| 0 | |
| 0.0375 | 0.396179108 |
| 0.075 | 0.342177935 |
| 0.1125 | 0.305097247 |
| 0.15 | 0.32503061 |
| 0.1875 | 0.316373871 |
| 0.225 | 0.307210677 |
| 0.2625 | 0.287808159 |
| 0.3 | 0.266467949 |
| 0.3375 | 0.235618967 |
| 0.375 | 0.215571055 |
| 0.4125 | 0.19078104 |
| 0.45 | 0.158410646 |
| 0.4875 | 0.126241655 |

| | |
|---|---|
| 0.525 | 0.101024423 |
| 0.5625 | 0.082839771 |
| 0.6 | 0.068551121 |
| 0.6375 | 0.0583591 |
| 0.675 | 0.054496712 |
| 0.7125 | 0.050498075 |
| 0.75 | 0.047386757 |
| 0.7875 | 0.043311871 |
| 0.825 | 0.04074987 |
| 0.8625 | 0.038905284 |
| 0.9 | 0.036345277 |
| 0.9375 | 0.03611919 |
| 0.975 | 0.035589072 |
| 1.0125 | 0.033285296 |
| 1.05 | 0.031821307 |
| 1.0875 | 0.030761876 |
| 1.125 | 0.030350513 |
| 1.1625 | 0.030398241 |
| 1.2 | 0.031320882 |
| 1.2375 | 0.032505294 |
| 1.275 | 0.034634523 |
| 1.3125 | 0.037962288 |
| 1.35 | 0.041484272 |
| 1.3875 | 0.045362593 |
| 1.425 | 0.048703985 |
| 1.4625 | 0.05276189 |
| 1.5 | 0.055798638 |
| 1.5375 | 0.058677506 |
| 1.575 | 0.059607995 |
| 1.6125 | 0.059010065 |
| 1.65 | 0.048689868 |
| 1.6875 | 0.047343311 |
| 1.725 | 0.051247473 |
| 1.7625 | 0.048087206 |
| 1.8 | 0.046222397 |
| 1.8375 | 0.043829358 |
| 1.875 | 0.043272947 |
| 1.9125 | 0.041425821 |
| 1.95 | 0.038361493 |
| 1.9875 | 0.047915285 |
| 2.025 | 0.044578962 |
| 2.0625 | 0.06020391 |
| 2.1 | 0.039472482 |

| | |
|---|---|
| 2.1375 | 0.036986344 |
| 2.175 | 0.033465416 |
| 2.2125 | 0.025079735 |
| 2.25 | 0.0176303 |
| 2.2875 | 0.058425873 |
| 2.325 | 0.186300019 |
| 2.3625 | 0.110356492 |
| 2.4 | 0.092503273 |
| 2.4375 | 0.268574024 |
| 2.475 | 0.184732528 |

**Figure 4j: MTLn3 invadopodia p190-KD**

Red-highlighted cells show p<0.05 compared to the center of invadopodia

| X-position (micron) | p-value |
|---|---|
| -2.25 | 0.050782474 |
| -2.2125 | 0.075852293 |
| -2.175 | 0.058310925 |
| -2.1375 | 0.069475634 |
| -2.1 | 0.103179033 |
| -2.0625 | 0.063463629 |
| -2.025 | 0.058986632 |
| -1.9875 | 0.060570371 |
| -1.95 | 0.053611941 |
| -1.9125 | 0.052763864 |
| -1.875 | 0.05209951 |
| -1.8375 | 0.053737485 |
| -1.8 | 0.050279228 |
| -1.7625 | 0.047196872 |
| -1.725 | 0.044118186 |
| -1.6875 | 0.041878844 |
| -1.65 | 0.039594551 |
| -1.6125 | 0.037208705 |
| -1.575 | 0.034750065 |
| -1.5375 | 0.032906844 |
| -1.5 | 0.03180472 |
| -1.4625 | 0.03099728 |
| -1.425 | 0.031654407 |
| -1.3875 | 0.031302972 |
| -1.35 | 0.030267969 |

| | |
|---|---|
| -1.3125 | 0.029111931 |
| -1.275 | 0.027360577 |
| -1.2375 | 0.025886194 |
| -1.2 | 0.024851269 |
| -1.1625 | 0.024331694 |
| -1.125 | 0.02421346 |
| -1.0875 | 0.024118902 |
| -1.05 | 0.024594397 |
| -1.0125 | 0.025593928 |
| -0.975 | 0.026312657 |
| -0.9375 | 0.025785745 |
| -0.9 | 0.024671493 |
| -0.8625 | 0.024796326 |
| -0.825 | 0.025606237 |
| -0.7875 | 0.025868018 |
| -0.75 | 0.026771134 |
| -0.7125 | 0.028048212 |
| -0.675 | 0.029914769 |
| -0.6375 | 0.032141399 |
| -0.6 | 0.036277966 |
| -0.5625 | 0.041224043 |
| -0.525 | 0.050654095 |
| -0.4875 | 0.061725638 |
| -0.45 | 0.079982369 |
| -0.4125 | 0.104205557 |
| -0.375 | 0.135419574 |
| -0.3375 | 0.177505206 |
| -0.3 | 0.212942247 |
| -0.2625 | 0.225119507 |
| -0.225 | 0.235656429 |
| -0.1875 | 0.252797892 |
| -0.15 | 0.29076287 |
| -0.1125 | 0.324339139 |
| -0.075 | 0.348726193 |
| -0.0375 | 0.48153016 |
| **0** | |
| 0.0375 | 0.013424108 |
| 0.075 | 0.016465449 |
| 0.1125 | 0.028289859 |
| 0.15 | 0.039791478 |
| 0.1875 | 0.049857137 |
| 0.225 | 0.055551281 |

| | |
|---|---|
| 0.2625 | 0.057721995 |
| 0.3 | 0.063805393 |
| 0.3375 | 0.063415755 |
| 0.375 | 0.065701646 |
| 0.4125 | 0.06661752 |
| 0.45 | 0.064690615 |
| 0.4875 | 0.064528098 |
| 0.525 | 0.067004869 |
| 0.5625 | 0.070253298 |
| 0.6 | 0.074282161 |
| 0.6375 | 0.075676377 |
| 0.675 | 0.076436752 |
| 0.7125 | 0.074520357 |
| 0.75 | 0.074045691 |
| 0.7875 | 0.072314384 |
| 0.825 | 0.07144668 |
| 0.8625 | 0.070543247 |
| 0.9 | 0.067955492 |
| 0.9375 | 0.064131489 |
| 0.975 | 0.059360173 |
| 1.0125 | 0.055602025 |
| 1.05 | 0.051282496 |
| 1.0875 | 0.047167878 |
| 1.125 | 0.044488198 |
| 1.1625 | 0.041270876 |
| 1.2 | 0.035261731 |
| 1.2375 | 0.034197906 |
| 1.275 | 0.031022597 |
| 1.3125 | 0.028537317 |
| 1.35 | 0.026890378 |
| 1.3875 | 0.025598446 |
| 1.425 | 0.02446709 |
| 1.4625 | 0.023140167 |
| 1.5 | 0.021841825 |
| 1.5375 | 0.020486301 |
| 1.575 | 0.020292167 |
| 1.6125 | 0.020986661 |
| 1.65 | 0.022520364 |
| 1.6875 | 0.025722371 |
| 1.725 | 0.027792642 |
| 1.7625 | 0.029648492 |
| 1.8 | 0.030264515 |
| 1.8375 | 0.032114313 |

| | |
|---|---|
| 1.875 | 0.033724173 |
| 1.9125 | 0.036140348 |
| 1.95 | 0.03765679 |
| 1.9875 | 0.034573558 |
| 2.025 | 0.035654023 |
| 2.0625 | 0.039355849 |
| 2.1 | 0.033188557 |
| 2.1375 | 0.027523077 |
| 2.175 | 0.035337232 |
| 2.2125 | 0.0373626 |
| 2.25 | 0.037970843 |
| 2.2875 | 0.052986975 |
| 2.325 | 0.050337147 |
| 2.3625 | 0.057073567 |
| 2.4 | 0.218757277 |
| 2.4375 | 0.22532759 |
| 2.475 | 0.4880494 |
| 2.5125 | 0.455204192 |
| 2.55 | 0.403096376 |

**Supplementary Figure S7: MDA-MB-231 invadopodia**

Red-highlighted cells show $p<0.05$ compared to the center of invadopodia

| X-position (micron) | p-value |
|---|---|
| -2.4 | 0.36661081 |
| -2.3625 | 0.27616192 |
| -2.325 | 0.39031267 |
| -2.2875 | 0.19272857 |
| -2.25 | 0.10117501 |
| -2.2125 | 0.02930602 |
| -2.175 | 0.33271778 |
| -2.1375 | 0.337406 |
| -2.1 | 0.34443526 |
| -2.0625 | 0.36192108 |
| -2.025 | 0.12687112 |
| -1.9875 | 0.03606141 |
| -1.95 | 0.02423647 |
| -1.9125 | 0.02708208 |
| -1.875 | 0.02623878 |
| -1.8375 | 0.02259117 |
| -1.8 | 0.01958408 |
| -1.7625 | 0.01691168 |

| | |
|---|---|
| -1.725 | 0.01462997 |
| -1.6875 | 0.01265248 |
| -1.65 | 0.0110351 |
| -1.6125 | 0.0099337 |
| -1.575 | 0.00968663 |
| -1.5375 | 0.0097351 |
| -1.5 | 0.01024929 |
| -1.4625 | 0.01114021 |
| -1.425 | 0.01238654 |
| -1.3875 | 0.01409839 |
| -1.35 | 0.01585096 |
| -1.3125 | 0.01771973 |
| -1.275 | 0.01986259 |
| -1.2375 | 0.02191907 |
| -1.2 | 0.02345927 |
| -1.1625 | 0.02493678 |
| -1.125 | 0.02635447 |
| -1.0875 | 0.02766339 |
| -1.05 | 0.02890687 |
| -1.0125 | 0.03019102 |
| -0.975 | 0.03180102 |
| -0.9375 | 0.03334727 |
| -0.9 | 0.03522549 |
| -0.8625 | 0.03740731 |
| -0.825 | 0.03972603 |
| -0.7875 | 0.04221071 |
| -0.75 | 0.04497575 |
| -0.7125 | 0.04913846 |
| -0.675 | 0.05440852 |
| -0.6375 | 0.06245967 |
| -0.6 | 0.07170322 |
| -0.5625 | 0.08608786 |
| -0.525 | 0.10246767 |
| -0.4875 | 0.12522983 |
| -0.45 | 0.15485995 |
| -0.4125 | 0.19483684 |
| -0.375 | 0.23944077 |
| -0.3375 | 0.29365732 |
| -0.3 | 0.34009279 |
| -0.2625 | 0.3861762 |
| -0.225 | 0.43252048 |
| -0.1875 | 0.48018583 |

| | |
|---|---|
| -0.15 | 0.48317933 |
| -0.1125 | 0.41352652 |
| -0.075 | 0.37485747 |
| -0.0375 | 0.31663943 |
| **0** | |
| 0.0375 | 0.11653715 |
| 0.075 | 0.09384527 |
| 0.1125 | 0.07028582 |
| 0.15 | 0.06634571 |
| 0.1875 | 0.05389081 |
| 0.225 | 0.04626272 |
| 0.2625 | 0.04161859 |
| 0.3 | 0.03711621 |
| 0.3375 | 0.03239859 |
| 0.375 | 0.02839538 |
| 0.4125 | 0.02566925 |
| 0.45 | 0.02422039 |
| 0.4875 | 0.02287208 |
| 0.525 | 0.02184326 |
| 0.5625 | 0.02105992 |
| 0.6 | 0.02048602 |
| 0.6375 | 0.02009409 |
| 0.675 | 0.01948284 |
| 0.7125 | 0.01889112 |
| 0.75 | 0.01815053 |
| 0.7875 | 0.0170939 |
| 0.825 | 0.01584866 |
| 0.8625 | 0.01470911 |
| 0.9 | 0.01328461 |
| 0.9375 | 0.01190243 |
| 0.975 | 0.01035734 |
| 1.0125 | 0.00861576 |
| 1.05 | 0.00711624 |
| 1.0875 | 0.00579983 |
| 1.125 | 0.00469465 |
| 1.1625 | 0.00400749 |
| 1.2 | 0.00401325 |
| 1.2375 | 0.00500566 |
| 1.275 | 0.00781752 |
| 1.3125 | 0.01406827 |
| 1.35 | 0.02440962 |
| 1.3875 | 0.03793254 |

| | |
|---|---|
| 1.425 | 0.05425427 |
| 1.4625 | 0.07294114 |
| 1.5 | 0.09541913 |
| 1.5375 | 0.12078098 |
| 1.575 | 0.15174364 |
| 1.6125 | 0.18427059 |
| 1.65 | 0.21690215 |
| 1.6875 | 0.24167521 |
| 1.725 | 0.2627792 |
| 1.7625 | 0.28598333 |
| 1.8 | 0.30033324 |
| 1.8375 | 0.34048641 |
| 1.875 | 0.36163741 |
| 1.9125 | 0.37990438 |
| 1.95 | 0.3949334 |
| 1.9875 | 0.40999731 |
| 2.025 | 0.43786287 |
| 2.0625 | 0.45270585 |
| 2.1 | 0.46271555 |
| 2.1375 | 0.40499116 |
| 2.175 | 0.41188695 |
| 2.2125 | 0.31322599 |
| 2.25 | 0.24384137 |
| 2.2875 | 0.22609376 |
| 2.325 | 0.22832288 |
| 2.3625 | 0.1971288 |
| 2.4 | 0.19515633 |
| 2.4375 | 0.47423152 |
| 2.475 | 0.3559208 |
| 2.5125 | 0.07721943 |

### Appendix 2:

TC10 FRET biosensor sequence:

mCerulean-cpmVenus version:

mCerulean1 (Syn.Mod.):
ATGGTGTCCAAAGGAGAAGAACTGTTTACAGGAGTGGTCCCTATTCTGGTGGAACTGGAT
GGAGATGTGAATGGACATAAATTTTCCGTGAGCGGAGAAGGAGAAGGAGACGCTACATAT
GGAAAACTGACACTGAAGTTTATTTGTACAACAGGAAAACTGCCTGTGCCTTGGCCTACA
CTGGTGACCACACTGACATGGGGAGTCCAGTGTTTTGCTAGGTATCCTGATCATATGAAA
CAGCATGATTTCTTTAAAAGCGCTATGCCTGAGGGATATGTGCAGGAAAGGACAATTTTC
TTTAAAGATGATGGAAATTATAAAACAAGGGCTGAAGTGAAATTTGAAGGAGATACACTG
GTGAATAGGATTGAACTGAAAGGAATTGATTTTAAAGAAGATGGAAATATTCTGGGACAT
AAACTGGAATATAATGCTATTAGCGACAATGTGTACATTACAGCTGATAAACAGAAAAAT
GGAATTAAGGCTAATTTTAAAATTAGGCATAATATTGAAGATGGAAGCGTGCAGCTGGCT
GATCATTATCAGCAGAATACACCTATTGGAGATGGACCTGTGCTGCTGCCTGATAATCAT
TATCTGTCCACACAGAGCAAACTGTCCAAAGATCCTAATGAAAAAAGGGACCATATGGTG
CTGCTGGAATTTGTGACAGCTGCCGGCATTACCCTGGGAATGGATGAACTGTATAAA

Linker:
GGATCC

PBD1:
AAGGAGCGCCCCGAGATTTCCCTCCCTTCCGATTTCGAACACACGATTCACGTCGGGTTC
GATGCTGTCACGGGGGAGTTCACGGGGATGCCTGAGCAGTGGGCCCGCCTGCTCCAGACG
TCCAACATCACGAAGTCCGAGCAGAAGAAAAACCCTCAGGCTGTCCTGGATGTCCTCGAA
TTTTACAACTCCAAGAAAACGTCCAACAGCCAGAAATACATGAGCTTCACGGATAAGTCC

Linker:
GGCAGCGGCGGCAAGCTTCCCCCCGGCAGCGGGGGCTCCGGC

PBD2 with H83/86D:
AAGGAACGGCCTGAAATCAGCCTGCCCAGCGACTTCGAGGACACCATCGATGTGGGCTTC
GACGCCGTGACCGGCGAGTTTACCGGCATGCCCGAACAGTGGGCTCGGCTCCTGCAGACC
AGCAACATCACCAAAAGCGAACAGAAAAAGAACCCCCAGGCCGTGCTGGACGTGCTGGAG
TTCTATAATAGCAAAAAGACCAGCAATTCCCAGAAGTATATGTCCTTCACCGACAAAAGC

Linker:
GCGGCCGCA

mcp229Venus:
ATGGGCAGCACCAGCGGCAGCGGCAAACCGGGCAGCGGCGAAGGCAGCATGGTGAGCAAG
GGCGAGGAGCTGTTCACCGGGGTGGTGCCCATCCTGGTCGAGCTGGACGGCGACGTAAAC
GGCCACAAGTTCAGCGTGTCCGGCGAGGGCGAGGGCGATGCCACCTACGGCAAGCTGACC
CTGAAGCTGATCTGCACCACCGGCAAGCTGCCCGTGCCCTGGCCCACCCTCGTGACCACC
CTGGGCTACGGCCTGATGTGCTTCGCCCGCTACCCCGACCACATGAAGCAGCACGACTTC

TTCAAGTCCGCCATGCCCGAAGGCTACGTCCAGGAGCGCACCATCTTCTTCAAGGACGAC
GGCAACTACAAGACCCGCGCCGAGGTGAAGTTCGAGGGCGACACCCTGGTGAACCGCATC
GAGCTGAAGGGCATCGACTTCAAGGAGGACGGCAACATCCTGGGGCACAAGCTGGAGTAC
AACTACAACAGCCACAACGTCTATATCACCGCCGACAAGCAGAAGAACGGCATCAAGGCC
AACTTCAAGATCCGCCACAACATCGAGGACGGCGGCGTGCAGCTCGCCGACCACTACCAG
CAGAACACCCCCATCGGCGACGGCCCCGTGCTGCTGCCCGACAACCACTACCTGAGCTAC
CAGTCCAAGCTGAGCAAAGACCCCAACGAGAAGCGCGATCACATGGTCCTGCTGGAGTTC
GTGACCGCCGCCGGG

Linker:
GAATTC

TC10:
ATGCCCGGAGCCGGCCGCAGCAGCATGGCTCACGGGCCCGGCGCGCTGATGCTCAAGTGC
GTGGTGGTCGGCGACGGGGCGGTGGGCAAGACGTGCCTACTCATGAGCTATGCCAACGAC
GCCTTCCCGGAGGAGTACGTGCCCACCGTCTTCGACCACTACGCAGTCAGCGTCACCGTG
GGGGGCAAGCAGTACCTCCTAGGACTCTATGACACGGCCGGACAGGAAGACTATGACCGT
CTGAGGCCTTTATCTTACCCAATGACCGATGTCTTCCTTATATGCTTCTCGGTGGTAAAT
CCAGCCTCATTTCAAAATGTGAAAGAGGAGTGGGTACCGGAACTTAAGGAATACGCACCA
AATGTACCCTTTTTATTAATAGGAACTCAGATTGATCTCCGAGATGACCCCAAAACTTTA
GCAAGACTGAATGATATGAAAGAAAAACCTATATGTGTGGAACAAGGACAGAAACTAGCA
AAAGAGATAGGAGCATGCTGCTATGTGGAATGTTCAGCTTTAACCCAGAAGGGATTGAAG
ACTGTTTTTGATGAGGCTATCATAGCCATTTTAACTCCAAAGAAACACACTGTAAAAAAA
AGAATAGGATCAAGATGTATAAACTGTTGTTTAATTACGTGA

Near infrared version:

miRFP720:
ATGGCCGAGGGCAGCGTGGCCCGCCAGCCCGACCTGCTGACCTGCGACGACGAGCCCATC
CACATCCCCGGCGCCATCCAGCCCCACGGCCTGCTGCTGGCCCTGGCCGCCGACATGACC
ATCGTGGCCGGCAGCGACAACCTGCCCGAGCTGACCGGCCTGGCCATCGGCGCCCTGATC
GGCCGCAGCGCCGCCGACGTGTTCGACAGCGAGACCCACAACCGCCTGACCATCGCCCTG
GCCGAGCCCGGCGCCGCCGTGGGCGCCCCCATCACCGTGGGCTTCACCATGCGCAAGGAC
GCCGGCTTCATCGGCAGCTGGCACCGCCACGACCAGCTGATCTTCCTGGAGCTGGAGCCC
CCCCAGCGCGACGTGGCCGAGCCCCAGGCCTTCTTCCGCCGCACCAACAGCGCCATCCGC
CGCCTGCAGGCCGCCGAGACCCTGGAGAGCGCCTGCGCCGCCGCCGCCCAGGAGGTGCGC
AAGATCACCGGCTTCGACCGCGTGATGATCTACCGCTTCGCCAGCGACTTCAGCGGCAGC
GTGATCGCCGAGGACCGCTGCGCCGAGGTGGAGAGCAAGCTGGGCCTGCACTACCCCGCC
AGCTTCATCCCCGCCCAGGCCCGCCGCCTGTACACCATCAACCCCGTGCGCATCATCCCC
GACATCAACTACCGCCCCGTGCCCGTGACCCCCGACCTGAACCCCGTGACCGGCCGCCCC
ATCGACCTGAGCTTCGCCATCCTGCGCAGCGTGAGCCCCAACCACCTGGAGTTCATGCGC
AACATCGGCATGCACGGCACCATGAGCATCAGCATCCTGCGCGGCGAGCGCCTGTGGGGC
CTGATCGTGTGCCACCACCGCACCCCCTACTACGTGGACCTGGACGGCCGCCAGGCCTGC
AAGCGCGTGGCCGAGCGCCTGGCCACCCAGATCGGCGTGATGGAGGAG

Linker:

GGATCC

PBD1:
AAGGAGCGCCCCGAGATTTCCCTCCCTTCCGATTTCGAACACACGATTCACGTCGGGTTC
GATGCTGTCACGGGGGAGTTCACGGGGATGCCTGAGCAGTGGGCCCGCCTGCTCCAGACG
TCCAACATCACGAAGTCCGAGCAGAAGAAAAACCCTCAGGCTGTCCTGGATGTCCTCGAA
TTTTACAACTCCAAGAAAACGTCCAACAGCCAGAAATACATGAGCTTCACGGATAAGTCC

Linker:
GGCAGCGGCGGCAAGCTTCCCCCCGGCAGCGGGGGCTCCGGC

PBD2 with H83/86D:
AAGGAACGGCCTGAAATCAGCCTGCCCAGCGACTTCGAGGACACCATCGATGTGGGCTTC
GACGCCGTGACCGGCGAGTTTACCGGCATGCCCGAACAGTGGGCTCGGCTCCTGCAGACC
AGCAACATCACCAAAAGCGAACAGAAAAAGAACCCCCAGGCCGTGCTGGACGTGCTGGAG
TTCTATAATAGCAAAAAGACCAGCAATTCCCAGAAGTATATGTCCTTCACCGACAAAAGC

Linker:
GCGGCCGGCACGTCTGGCTCCGGGAAAGGCAGCGGGGAAGGCTCCACCAAGGGGACCTCC
GGGAGCGGCAAGGGGTCCGGCGAGGGAAGCACGAAAGGCGGCAGCGCTGCCGGCACATCT
GGAAGCGGCAAGGGCTCTGGGGAGGGGTCCACTAAAGGAGGGAGCGCGGCCGCT

miRFP670:
ATGGTGGCCGGCCACGCCAGCGGCAGCCCCGCCTTCGGCACCGCCAGCCACAGCAACTGC
GAGCACGAGGAGATCCACCTGGCCGGCAGCATCCAGCCCCACGGCGCCCTGCTGGTGGTG
AGCGAGCACGACCACCGCGTGATCCAGGCCAGCGCCAACGCCGCCGAGTTCCTGAACCTG
GGCAGCGTGCTGGGCGTGCCCCTGGCCGAGATCGACGGCGACCTGCTGATCAAGATCCTG
CCCCACCTGGACCCCACCGCCGAGGGCATGCCCGTGGCCGTGCGCTGCCGCATCGGCAAC
CCCAGCACCGAGTACTGCGGCCTGATGCACCGCCCCCCCGAGGGCGGCCTGATCATCGAG
CTGGAGCGCGCCGGCCCCAGCATCGACCTGAGCGGCACCCTGGCCCCCGCCCTGGAGCGC
ATCCGCACCGCCGGCAGCCTGCGCGCCCTGTGCGACGACACCGTGCTGCTGTTCCAGCAG
TGCACCGGCTACGACCGCGTGATGGTGTACCGCTTCGACGAGCAGGGCCACGGCCTGGTG
TTCAGCGAGTGCCACGTGCCCGGCCTGGAGAGCTACTTCGGCAACCGCTACCCCAGCAGC
ACCGTGCCCCAGATGGCCCGCCAGCTGTACGTGCGCCAGCGCGTGCGCGTGCTGGTGGAC
GTGACCTACCAGCCCGTGCCCCTGGAGCCCCGCCTGAGCCCCCTGACCGGCCGCGACCTG
GACATGAGCGGCTGCTTCCTGCGCAGCATGAGCCCCTGCCACCTGCAGTTCCTGAAGGAC
ATGGGCGTGCGCGCCACCCTGGCCGTGAGCCTGGTGGTGGGCGGCAAGCTGTGGGGCCTG
GTGGTGTGCCACCACTACCTGCCCCGCTTCATCCGCTTCGAGCTGCGCGCCATCTGCAAG
CGCCTGGCCGAGCGCATCGCCACCCGCATCACCGCCCTGGAGAGC

Linker:
GGCAGCGGCTCCGGGAGCGGGTCCGGAGGCGAATTC:

TC10:
ATGCCCGGAGCCGGCCGCAGCAGCATGGCTCACGGGCCCGGCGCGCTGATGCTCAAGTGC
GTGGTGGTCGGCGACGGGGCGGTGGGCAAGACGTGCCTACTCATGAGCTATGCCAACGAC
GCCTTCCCGGAGGAGTACGTGCCCACCGTCTTCGACCACTACGCAGTCAGCGTCACCGTG

GGGGGCAAGCAGTACCTCCTAGGACTCTATGACACGGCCGGACAGGAAGACTATGACCGT
CTGAGGCCTTTATCTTACCCAATGACCGATGTCTTCCTTATATGCTTCTCGGTGGTAAAT
CCAGCCTCATTTCAAAATGTGAAAGAGGAGTGGGTACCGGAACTTAAGGAATACGCACCA
AATGTACCCTTTTTATTAATAGGAACTCAGATTGATCTCCGAGATGACCCCAAAACTTTA
GCAAGACTGAATGATATGAAAGAAAAACCTATATGTGTGGAACAAGGACAGAAACTAGCA
AAAGAGATAGGAGCATGCTGCTATGTGGAATGTTCAGCTTTAACCCAGAAGGGATTGAAG
ACTGTTTTTGATGAGGCTATCATAGCCATTTTAACTCCAAAGAAACACACTGTAAAAAAA
AGAATAGGATCAAGATGTATAAACTGTTGTTTAATTACGTGA
